# A chimeric infection program and epigenetic conflict underpin endogenous giant virus latency

**DOI:** 10.64898/2026.10.09.758095

**Authors:** Uri Sheyn, Richard Heery, Maria P Erazo-Garcia, Lan Xu, Rory J Craig, Alex de Mendoza, Frank O Aylward

## Abstract

Giant endogenous viral elements (GEVEs) are ubiquitous in eukaryotic genomes, but how the virus adapts to the host and is capable of reactivation despite host defense mechanisms remains unclear. Here we examined this process in the green alga *Chlamydomonas reinhardtii* and its latent giant virus, punuivirus, which is integrated into the host genome. We reveal a chimeric genomic architecture in which punuivirus latency genes are enriched in spliceosomal introns and 6-methyladenine marks, mirroring host transcriptional regulation, while virulence genes are predominantly controlled by a conserved viral promoter. Using single-cell RNA-seq, we show that nearly 70% of cells express viral transcripts, but only a small subpopulation commits to viral production, expressing virulence genes with host transcriptional shutoff. Moreover, we find that the intact punuivirus GEVE remains refractory to host 5-methylcytosine (5mC) silencing, whereas a naturally occurring deletion variant is associated with widespread 5mC accumulation across the GEVE and elsewhere in the host genome. The deleted region encodes a highly expressed ten-eleven translocation dioxygenase (TET), and epigenetic profiling of intact and deletion strains implicates this GEVE gene in removing 5mC marks, thereby maintaining the virus in a transcriptionally permissive latent state. Our study provides novel insights into the molecular underpinnings of host-virus co-existence in a giant virus-alga interaction.

**Graphical Abstract:** 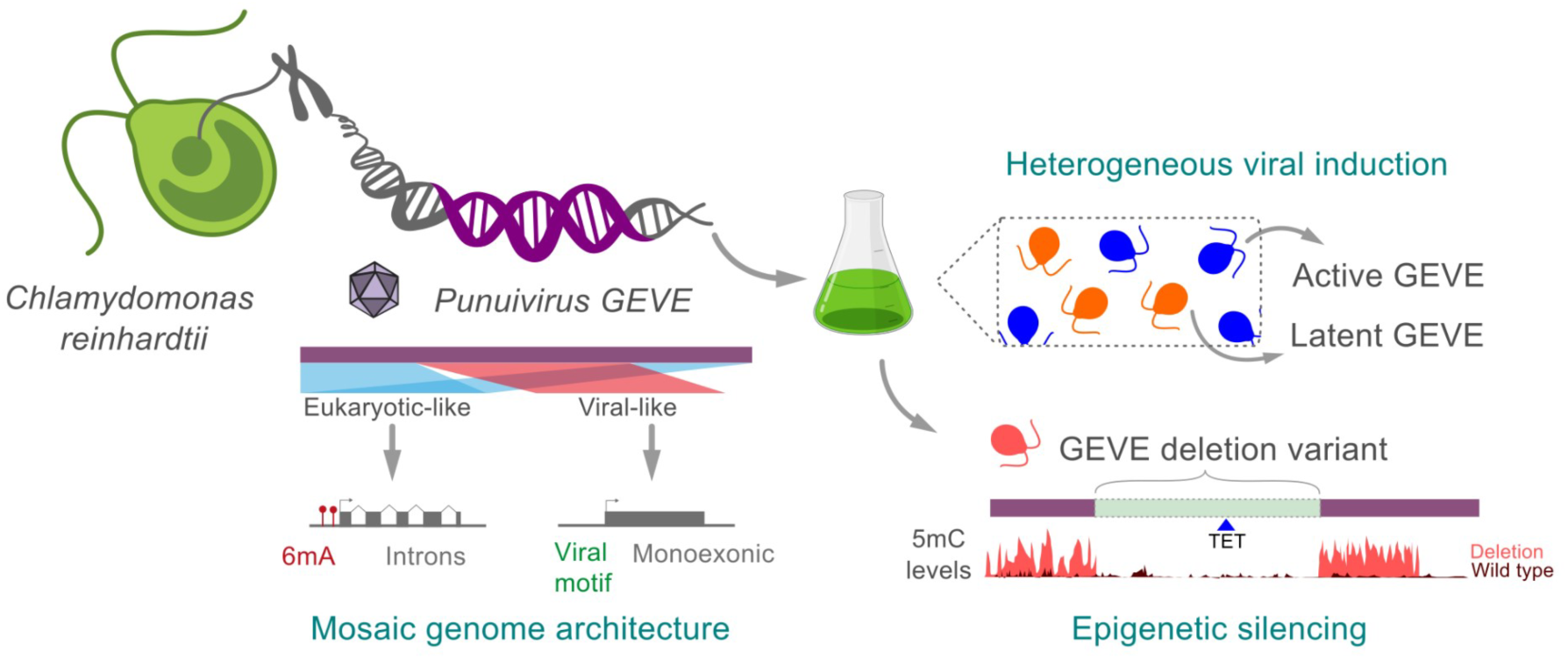

## Introduction

Latency is a common strategy employed by diverse viruses that often involves integration into the host genome^1^. This represents an alternative to lytic infection, allowing viruses to persist within host cells for prolonged periods while retaining the capacity to reactivate. Unlike obligately lytic viruses, latent viruses must balance two competing demands: remaining capable of future viral production while avoiding both host defense mechanisms and excessive damage to the host. How latent viruses achieve this balance remains poorly understood, particularly in eukaryotic viruses. Moreover, latency and host genome integration are important gateways for domestication and the transfer of novel genetic material into cellular genomes because much of the viral genetic material has already evolved to function in the transcriptional landscape of the host^2,3^. As such, understanding latent infection strategies and long-term host-virus coexistence has great importance for eukaryotic genome evolution.

*Giant <u>e</u>ndogenous <u>v</u>iral <u>e</u>lements* (GEVEs) derived from giant viruses are the largest integrated viral elements characterized to date and provide an exceptional opportunity to study latency, but they have been characterized only recently and their impact on host biology remains unclear. To date, the vast majority of cultivated giant viruses are obligately lytic^4^, but the recent discovery of widespread GEVEs suggests that latent infection programs are also commonly employed by these viruses in nature. Integrative giant viruses were studied in brown algae as early as the 1970s^5^, but their discovery in other lineages lagged far behind. Work in green algae provided the first evidence that GEVEs were widespread in nature^6^, and they were subsequently described in a range of lineages from across the eukaryotic tree of life, including euglenoids^7^, other brown algae^8^, fungi^9–11^, amoebae^2^, and unicellular relatives of animals^2^. In some protist genomes >15% of the genes are encoded within hundreds of different GEVEs^2^, and individual elements can reach lengths of 1.5 million base-pairs^9^ or form tandem arrays of elements^3^, underscoring their impact on eukaryotic genome evolution. Most GEVEs are degraded relics of past infections and are epigenetically silenced by the host^2,12–15^, however, and as a result there are few study systems available for examining interactions between hosts and virion-producing GEVEs.

Recently, a 617 kbp GEVE was discovered in the model green alga *Chlamydomonas reinhardtii* that can reactivate and form virions in late-stage cultures^16^. The virus produced by this GEVE, called punuivirus, represents the largest latent virus discovered to date and a unique model for studying latent infection dynamics in a giant virus. Prior work has shown that virions are produced in post-exponential cultures of *C. reinhardtii* that correspond to a shift in GEVE gene expression^16^, but it remains unclear how the punuivirus genome has evolved to effectively interact with host transcriptional machinery during periods of latency, or how many cells experience a switch to virulent infection. Moreover, it is unknown whether the host has evolved defense mechanisms to target punuivirus, or whether epigenetic modifications could be effective against an active GEVE. To date, host defense systems deployed against virulent giant viruses have remained elusive, with those described to date relying on third-party interactions with hyperparasitic viruses or intracellular bacteria^17–19^. In this study we address these questions by employing a suite of genomic, single-cell transcriptomic, and epigenomic approaches. Our results provide insight into the molecular underpinnings of giant virus latency and long-term host coexistence.

## Results and Discussion

### Mosaic genome architecture of punuivirus

To better define the genetic landscape of both punuivirus and the host alga, we generated a new gapless assembly of *C. reinhardtii* CC-2937 using Oxford Nanopore Technologies (ONT) sequencing. We then re-annotated the punuivirus genome using both RNA-seq and ONT data to better curate the gene boundaries and epigenetic features of this endogenous viral element. We used a hybrid gene prediction strategy that leveraged both Prodigal and BRAKER3 for virus-like and eukaryote-like reading frames, respectively. We manually verified gene boundaries using RNA-seq data, and identified modified bases such as N6-methyladenine (6mA) near transcription start sites by mapping of ONT reads (see Methods). This resulted in a total of 506 genes, of which 61 contained spliceosomal introns that were not identified in the initial genome annotation of this virus (**Figure 1**). Spliceosomal introns had a mean length of 176 bp, with an average of 2.5 introns per gene, and exhibited conserved features similar to those observed in *C. reinhardtii*, including 3′ cytosine-enriched ends and canonical GT–AG splice-site motifs^20^ (**Figure S1A,B**). In addition, we identified 53 genes with 6mA marks near transcriptional start sites (TSSs) (**Figure 1**).

**Figure 1.**
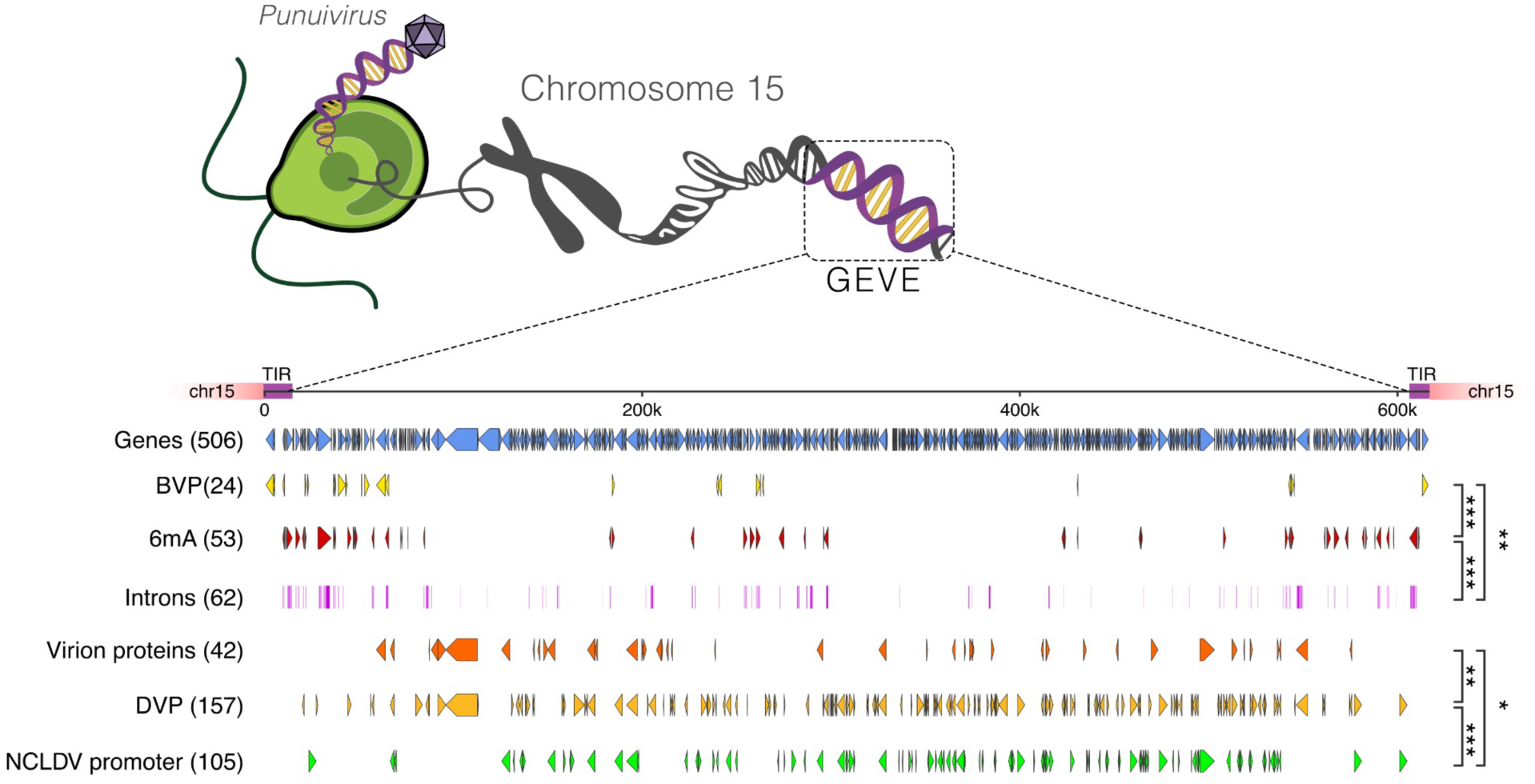
Linear map of the GEVE genome. Black/gray arrows show the 506 annotated genes (arrowhead = strand direction). TIR1/TIR2: terminal inverted repeats. Colored ticks below the gene track mark six binary genomic features (BVP, DVP, 6mA, Introns, NCLDV Promoter, Virion Proteins). Right-facing brackets connect feature pairs with significant positive co-occurrence (Fisher’s exact test; BH-FDR q < 0.05; OR > 1); bracket line width is proportional to log₂(OR). Significance level of each pair is annotated with asterisks (* q < 0.05, ** q < 0.01, *** q < 0.001).

In a previous study, we generated RNA-seq data from CC-2937 across different stages of a growth curve in which we observed virion production in stationary phase cultures. Using these data together with our new gene annotations, we defined sets of genes with enriched expression before viral production (BVP; n=24) and during viral production (DVP, n=157). Interestingly, we found that both intron-bearing genes and genes with upstream 6mA marks were strongly co-enriched (pairwise Fisher’s exact test; odds ratio (OR) = 14.25, q = 5.20 × 10⁻¹⁴). Of all genes that carry upstream 6mA modifications, 53% also contained spliceosomal introns, and both features were further co-enriched with BVP-cluster membership (6mA × BVP: q = 2.02 × 10⁻⁴; introns × BVP: OR = 4.96, q = 2.56 × 10⁻³), suggesting that they are linked to a common transcriptional program (**Figure S2**; Supplementary Table 5).

To identify punuivirus genes associated with virulent infection, we analyzed promoter regions immediately upstream of the TSSs and found that they exhibit a pronounced AT-rich composition, consistent with promoter sequences of other nucleocytoviruses (**Figure S1C**)^21^. Within these regions, we detected a conserved AAANAACATAAG motif (**Figure S1D**), closely resembling enriched promoter sequences of virulent giant viruses ^21,22^. This motif was present upstream of 105 genes, which are likely transcribed by the virus-encoded multi-subunit RNA polymerase^23^. These genes were significantly enriched within the DVP cluster (OR = 2.63; q = 1.93 × 10⁻⁴), confirming that these are expressed during periods of virion production. We next cross-referenced the 42 genes whose protein products were detected in our previous LC-MS/MS proteomics analysis of viral particles^16^. These virion-associated genes were significantly co-enriched with the DVP cluster (OR = 2.68; q = 5.60 × 10⁻³) and with genes carrying the nucleocytovirus-like promoter (OR = 2.31; q = 2.66 × 10⁻²), whereas none of them carried 6mA modifications (q = 2.61 × 10⁻²) (**Supplementary Table 5**, **Figure S2**, **Figure 1**). Together, these results delineate a set of genes enriched during virulent infection and late-stage virion assembly.

Collectively, these patterns define two modules within the punuivirus GEVE that are distinguished by their regulatory features. While the latent program is marked by constitutive gene expression, 6mA modification upstream of TSSs, and spliceosomal introns, the virulence program is characterized by expression during viral production, promoter elements characteristic of other virulent viruses, and virion structural protein genes (**Figure S2**). These co-occurrence patterns suggest that punuivirus retains a coherent, bipartite architecture that has evolved to exploit the regulatory environment of its host and adapt to its temperate infection strategy. In *C. reinhardtii*, host 6mA marks and spliceosomal introns are markers of active transcription^24,25^, so the tight co-occurrence of 6mA modification and spliceosomal introns within BVP-cluster genes suggests that latent-module loci are recognized by host transcriptional machinery as host-like genes, consistent with their elevated expression before viral production becomes detectable in culture. By contrast, genes expressed during virion production carry nucleocytovirus-type promoter motifs, which in other lytic giant viruses have been shown to follow an early/late temporal expression logic^21,22^.

### GEVE transcriptional activity escalates into a lytic gene program in a sub-population

Our identification of a chimeric genome architecture in punuivirus raises key questions regarding how the latent and virulent programs are regulated at a population-level. In particular, it remains unclear whether the low levels of viral production observed in algal cultures are due to low viral production across a large number of cells (e.g. budding) or a more virulent infection program in a small sub-population. To address these questions, we employed single-cell RNA sequencing (scRNA-seq) using a split-pool barcoding approach (Parse Biosciences; see STAR Methods). We applied this approach to the same biomass that we previously sampled for bulk RNA-seq, which spans 4 to 7 days post-inoculation (DPI) of the algal culture for which we identified clear viral production at later timepoints. Our scRNA-seq analysis recovered 22,421 QC-passing cells, of which 15,640 had at least one Unique Molecular Identifier (UMI) mapping to a GEVE gene (69.8%). Overall, cells with high-levels of GEVE expression were rare, however, with only 39 of the cells exceeding a threshold we defined as representing high GEVE activity (three standard deviations above mean GEVE UMI counts, or ≥43; **Figure 2A**; STAR Methods).

**Figure 2.**
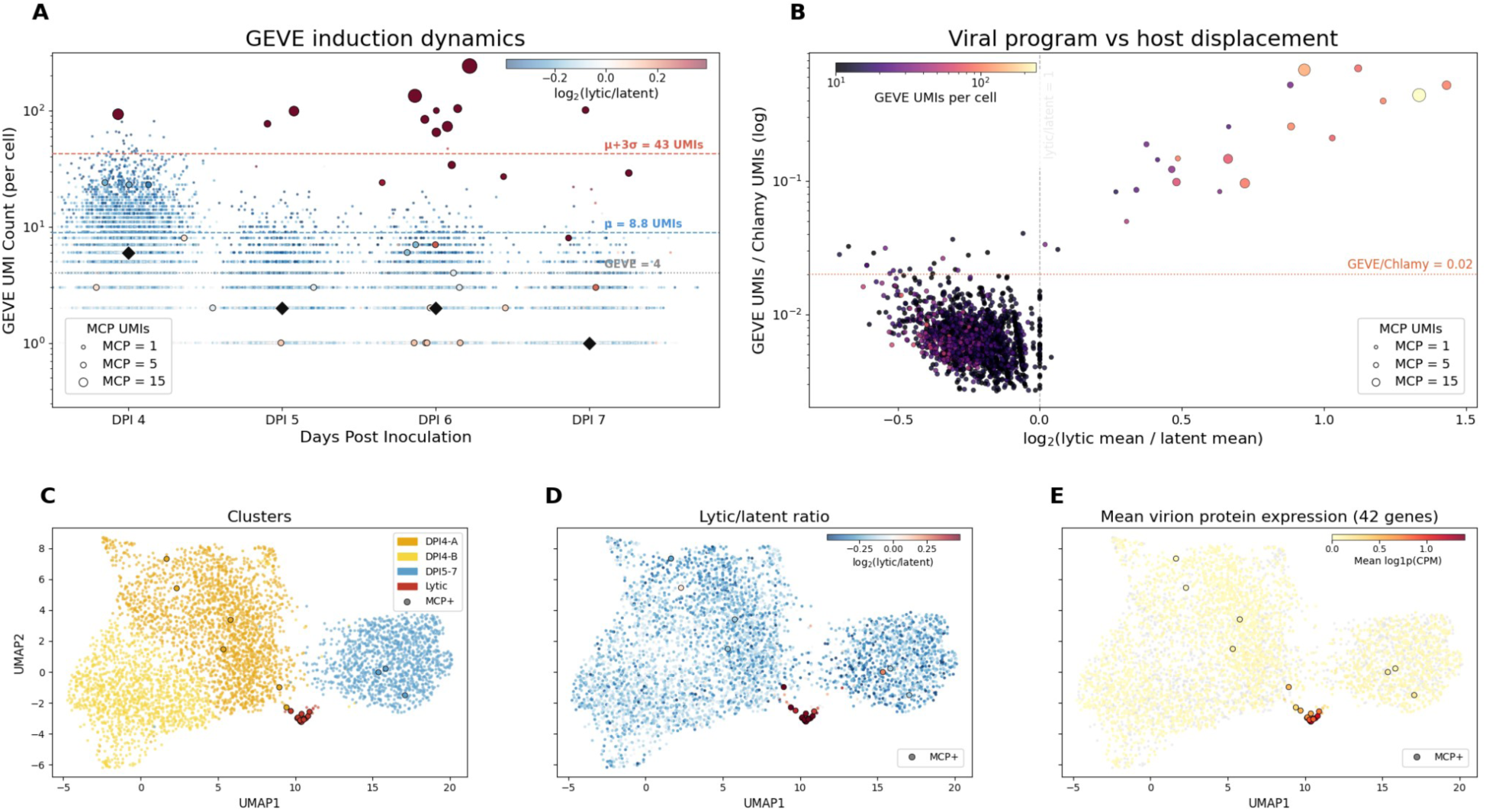
GEVE transcriptional dynamics and single-cell developmental trajectory. (**A**) GEVE UMI counts per cell across days post-inoculation (DPI 4–7), colored by lytic/latent gene-program ratio. Each dot represents one GEVE-expressing cell; GEVE-silent cells are omitted. Color encodes the log₂ ratio of mean log1p-normalized expression across 51 lytic-module genes versus 34 latent-module genes (diverging RdBu scale: red = lytic-dominant, blue = latent-dominant, white = balanced). Cells with detectable MCP expression are redrawn on top with a black outline, with dot size proportional to MCP UMI count. Filled black diamonds (◆) mark the per-DPI median GEVE UMI count among GEVE-expressing cells. Three horizontal reference lines mark the noise floor (GEVE = 4, gray dotted), the geometric mean of the log-normal fit to GEVE > 4 cells (μ = 8.8 UMIs, blue dashed), and the High-GEVE threshold (μ+3σ = 43 UMIs, red dashed). Numbers below each column show the count and percentage of GEVE-expressing cells at that DPI; the y-axis is log-scaled. (**B**) Scatter plot of lytic/latent ratio score (x-axis) against GEVE/Chlamy UMI ratio (y-axis, log scale) for all cells with GEVE ≥ 10 UMIs (n = 1,735), colored by total GEVE UMIs per cell (magma scale, log-normalized); dot size is proportional to MCP UMIs. The dashed vertical line marks a log₂ ratio of 0 (balanced lytic/latent expression); the y-axis ratio is computed as (GEVE UMIs + 0.5) / (Chlamy UMIs + 0.5). (**C-E**) UMAP of 4,688 cells with GEVE > 4 UMIs, computed on HVG + GEVE gene PCA; each dot is one cell. (**C**) Cells colored by cluster assignment: DPI4-A (dark orange), DPI4-B (yellow), DPI5-7 (blue), Lytic/Virocell (red). (**D**) Cells colored by lytic/latent module ratio score, computed as in (**B**) (diverging RdBu scale centered at 0). (**E**) Same UMAP colored by mean expression (log1p CPM) across the 42 annotated virion structural protein genes; cells with zero detected virion protein transcripts are shown in gray. In panels (**C-E**), cells with MCP UMIs > 0 are redrawn with a black outline.

We sought to understand whether high-GEVE cells simply contained more viral transcripts or represented a distinct lytic transcriptional state. To address this, we grouped the 39 cells by hierarchical clustering, identifying two sets: 12 virocells with high expression of DVP-cluster, nucleocytovirus promoter-driven, virion structural, and lytic module genes (including the major capsid protein, MCP), and 27 High-GEVE latent cells with enriched expression of latent module, BVP, and 6mA-marked genes (**see STAR Methods; Appendix**). Because high GEVE UMI counts alone do not reliably distinguish these states, we scored all GEVE-expressing cells using two additional metrics: the ratio of lytic to latent GEVE gene cluster UMIs, and the ratio of GEVE to host UMI counts (**see STAR Methods**). This yielded a final set of 21 virocells (**Figure 2B**), 20 of which were found in postexponential cultures (DPI 5-7; **Figure 2A**), consistent with viral production occurring primarily in later-stage cultures. Virocells exhibited coherent, high-intensity activation of DVP-cluster genes with nucleocytovirus promoters and virion structural proteins, while High-GEVE latent cells were enriched for the latent module, BVP, and 6mA-marked gene expression and lacked any signatures of the lytic program (**Figure S3**). The independent emergence of this chimeric expression program from the scRNA-seq data, recapitulating signatures defined in our earlier genomic analysis, underscores the stark mosaicism underpinning punuivirus transcriptional regulation.

To better characterize the global transcriptional landscape of GEVE-expressing cells, we performed UMAP dimensionality reduction on the 4,688 cells with GEVE > 4 UMIs, using the top 2,000 highly variable *C. reinhardtii* genes together with all expressed GEVE genes (**Figure 2C–D**). We resolved four stable clusters: two exponential-phase clusters enriched for DPI 4 cells (DPI4-A and DPI4-B), a cluster enriched for post-exponential DPI 5–7 cells, and a small, isolated cluster of cells with high GEVE expression and MCP detection. This isolated cluster showed a clear enrichment in lytic/latent ratio relative to the rest of the population (**Figure 2D**), consistent with our earlier finding of a sharp transcriptional shift to a lytic infection program. To validate that the virocell state corresponds to productive viral replication, we examined expression of genes encoding proteins previously identified in purified virions by LC-MS/MS. Twenty-seven of the forty-two virion protein genes were detected at significantly higher rates in virocells than in all latent GEVE cells (≥ 10 UMIs), whereas only one was preferentially detected in all latent GEVE cells (**Figure 2E**). Overall, the distinct transcriptional profile of virocells at the single-cell level together with their emergence specifically in later stage cultures is reminiscent of threshold-crossing dynamics in spontaneous prophage induction. This is consistent with viral bet-hedging strategies in which only a small subpopulation commits to productive replication^26,27^, as well as the small number of virion-containing cells reported in electron microscopy studies of algae associated with punuivirus relatives^16^.

### Viral induction is accompanied by transcriptional displacement of host mRNA

We next sought to determine how activation of the lytic program affects the overall transcriptional landscape of virocells, and specifically whether virocells experience the host transcript shutdown reported in virulent giant virus infections of other protist hosts ^28–30^. Grouping all 22,421 QC-passing cells by cell state (**Figure 3A; STAR Methods**), *C. reinhardtii* nuclear transcripts constituted 92.4% of total UMIs in GEVE-background cells (GEVE < 10 UMIs), with a modest rise in GEVE fraction to 0.6% in both GEVE-active and High-GEVE cells and a corresponding small decline in host transcripts. In virocells, by contrast, the GEVE fraction rose sharply to 17.3% of total UMIs while the host fraction fell to 78.4%, indicating that viral transcription accounts for roughly one in six sequenced molecules in these cells. This compositional shift was significant at the single-cell level: host UMI fraction was lower in virocells (median 80.0%) than in High-GEVE cells (median 89.2%; rank-biserial r = -0.78, p = 4.8 x 10^-6^), confirming that lytic viral transcription is accompanied by a displacement of host mRNA rather than a simple addition of viral transcripts on top of an unchanged host transcriptome (**Figure 3B**). Notably, this displacement was specific to cells with a high lytic/latent expression ratio: High-GEVE cells, despite having total GEVE UMI counts comparable to or exceeding those of many virocells, did not show a high lytic/latent ratio and correspondingly showed no meaningful reduction in host transcript fraction, indicating that displacement tracks with lytic program dominance rather than with GEVE UMI load alone.

**Figure 3.**
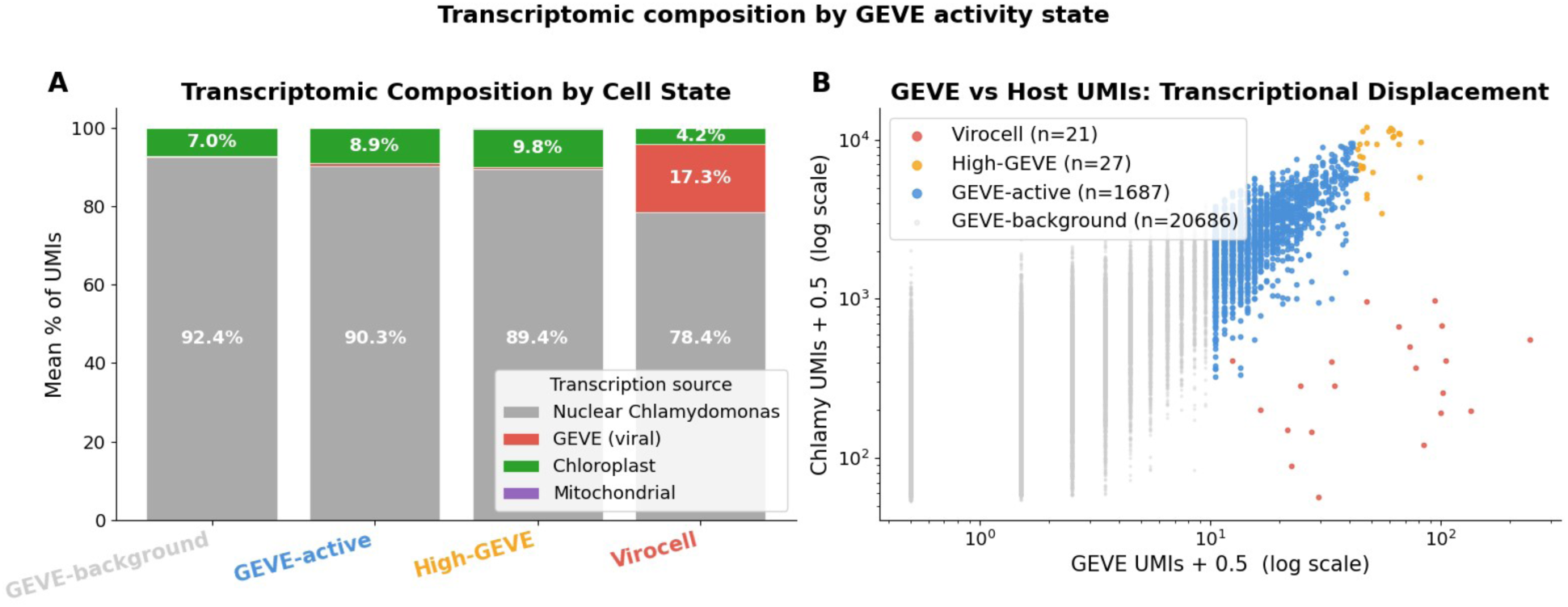
Transcriptomic composition by GEVE activity state. Cell states are defined by GEVE UMI count and lytic/latent program: GEVE-background (below 10 GEVE UMIs), GEVE-active (10–42 UMIs, not virocell; n = 1,687), High-GEVE (GEVE ≥ 43 UMIs, not virocell; n = 27), and Virocell (GEVE ≥ 10, log₂ lytic/latent > 0, GEVE/*C. reinhardtii* > 0.02; n = 21), across all 22,421 QC-passing cells. (**A**) Stacked bar chart of mean UMI percentage per gene type (Nuclear *C. reinhardtii* host, GEVE viral, chloroplast, mitochondrial) for each cell state. GEVE viral transcripts rise from < 1% in GEVE-background cells to ∼17% in virocells, with host UMI fraction significantly lower in virocells than High-GEVE cells (rank-biserial r = -0.78, p = 4.8 x 10^-6^, Mann-Whitney U test). (**B**) Scatter plot of raw GEVE UMIs vs raw *C. reinhardtii* UMIs per cell (log scale, +0.5 pseudocount), colored by cell state, showing virocells shifted toward lower host UMI counts relative to High-GEVE and GEVE-active cells at comparable GEVE UMI levels.

The rise of the viral transcript fraction to 17% of total UMIs and its negative correlation with host transcript counts within virocells is indicative of transcriptional displacement rather than simple addition, consistent with host shutoff described in other giant-virus systems. The genes preferentially upregulated in virocells include virion structural components, viral DNA replication machinery, and ankyrin repeat proteins likely involved in protein-protein interactions, which are all indicative of viral reprogramming of the host cell to support viral production^31^. Despite this clear transcriptional shift in virocells, the degree of host shutoff we observed is lower than that observed in virulent giant viruses, such as Emiliania huxleyi virus 1 (EhV1). For this virus, studies using scRNA-seq have reported almost complete viral takeover of the transcript pool during late-stage infection^28,30^. Bulk RNA-seq during giant virus infection in other host-virus systems, such as giant viruses that infect hosts in the genera *Chlorella* or *Acanthamoeba*, revealed more modest displacement of host mRNA in early stage infections, however (21-24% viral mRNA)^32,33^, and it remains unclear whether the dramatic host shutdown observed in infections of some virulent giant viruses are typical or simply one extreme across a wide spectrum.

Collectively, our scRNA-seq analyses show that the GEVE exists predominantly in a latent state that is actively maintained by a unique transcriptional program, and that only a small fraction of cells transition to virulent infection through activation of a distinct set of genes. The coexistence of a small population of virocells within a much larger pool of cells expressing a distinct viral transcriptional program demonstrates that latency is an actively maintained transcriptional state rather than simply a consequence of low viral activity. This motivated us to further examine the epigenetic landscape of the GEVE and clarify its potential role in mediating host-virus interactions and GEVE persistence over time.

### Partial GEVE deletion is associated with widespread reconfiguration of viral and host DNA methylation

To understand if host epigenetic silencing is regulating GEVE expression, we reanalyzed previously generated ONT data from three independent CC-2937 cultures^16^, calling both 5-methylcytosine (5mC) and 6-methyladenine (6mA). As reported previously^24,34–37^, the *C. reinhardtii* genome accumulates 5mC primarily at CG dinucleotides (mCG) within centromeres and subtelomeres, whereas 6mA is enriched at AT dinucleotides (mAT) around the transcription start sites of actively transcribed genes (see **Appendix**), just as we observe in latency-related genes of the punuivirus GEVE. Unexpectedly, one culture displayed a pronounced reduction in sequencing coverage across the central region of the GEVE, with split read mappings supporting a large internal deletion spanning ∼280 kbp and encompassing 277 genes (**Figure 4A**). This same culture also exhibited substantially higher levels of 5mC across the GEVE than the other two cultures (**Figure 4B**). Inspection of individual long reads revealed that all three cultures used for initial genome sequencing actually contained a mixture of full-length GEVE molecules (GEVE+) and molecules carrying the deletion (GEVE−), but at markedly different frequencies, ranging from approximately 0.5% to 58% of reads. In our scRNA-seq dataset, the majority of GEVE-expressing cells (66%) contained matches to transcripts found in the deleted region, indicating that GEVE– strains were not prevalent in this population. Collectively, these results indicate that the deletion variant segregates within the stock culture and can become highly abundant in some subcultures.

**Figure 4.**
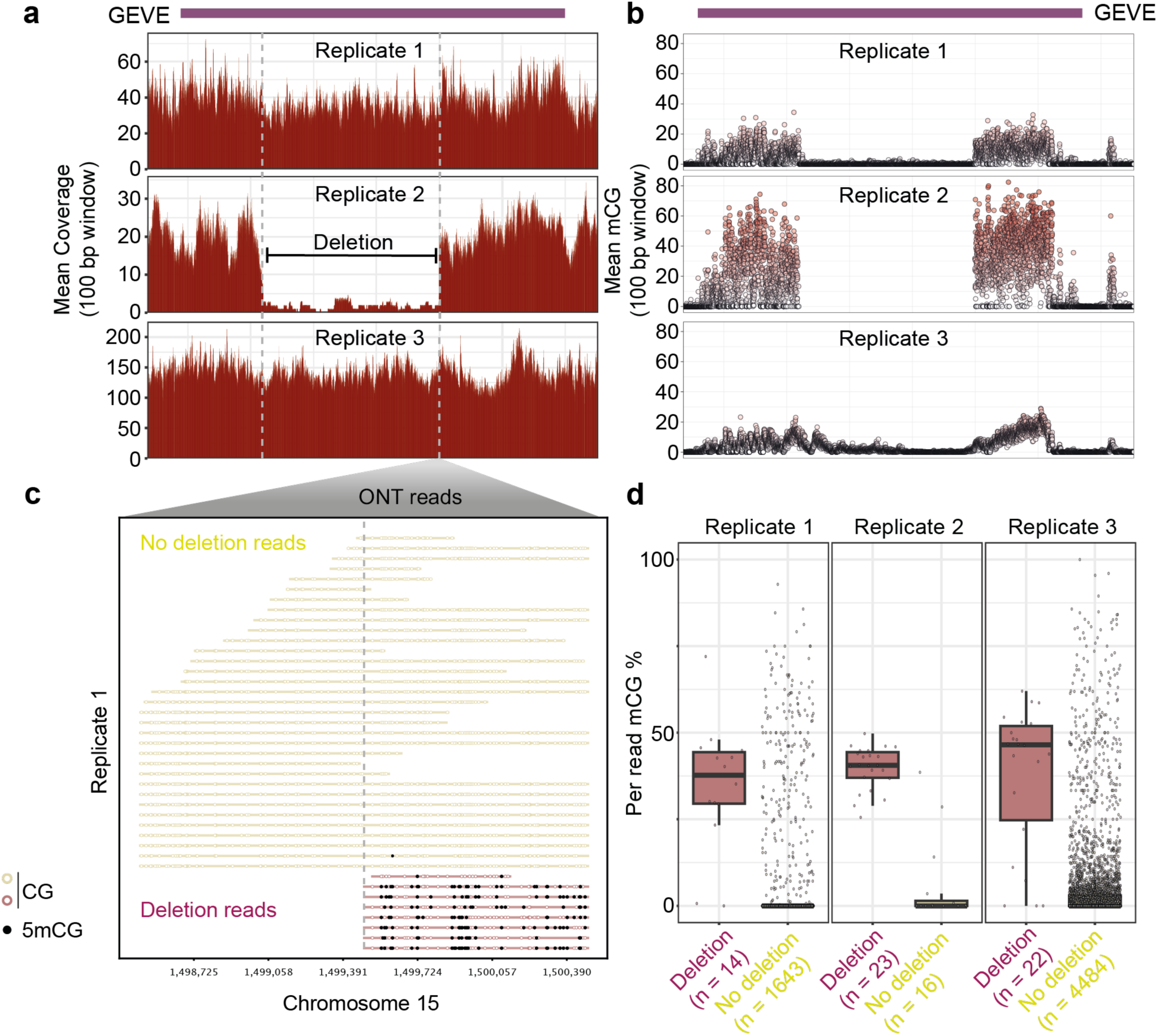
A naturally occurring GEVE deletion variant is associated with elevated DNA methylation. (A) Long-read coverage across the punuivirus GEVE in three independently sequenced CC-2937 cultures, revealing a recurrent internal deletion (∼280 kb). (B) Mean CpG methylation (5mC) across the GEVE in the same three mixed cultures. Cultures containing a higher proportion of deletion variants exhibit increased 5mC levels across the remaining GEVE sequence. (C) Per-read 5mC profiles from one representative mixed culture. Reads spanning the deleted interval (full-length GEVE molecules) show little CpG methylation, whereas reads terminating at the deletion breakpoint (deletion-GEVE molecules) display extensive CpG methylation across the retained GEVE sequence. The dashed vertical line marks the 3′ deletion breakpoint; filled circles indicate methylated CpGs and open circles unmethylated CpGs. (D) Distribution of per-read GEVE CpG methylation levels for deletion and full-length GEVE molecules across the three mixed cultures. Deletion-containing molecules consistently show substantially higher 5mC levels than full-length GEVE molecules.

To determine whether the elevated 5mC was associated specifically with either GEVE+ or GEVE– strains, we examined DNA methylation at the single-read level. Reads carrying the deletion were consistently hypermethylated for 5mC in all three cultures, whereas full-length GEVE reads remained largely unmethylated (**Figure 4C,D; Appendix**). To further validate this observation, we isolated clonal GEVE+ and GEVE− lines by single-colony picking followed by PCR screening across the deletion breakpoint (**Figure 5A**). Two independent isolates of each genotype were sequenced using ONT, confirming that the GEVE− lines consisted exclusively of the deletion allele, whereas the GEVE+ lines contained only full-length GEVE variants. The epigenetic landscape of the two genotypes differed strikingly. Whereas the full-length GEVE remained almost completely devoid of 5mC methylation, the remaining GEVE sequence in GEVE− isolates became extensively hypermethylated (**Figure 5B**). Thus, deletion of approximately half of the viral genome is accompanied by rapid acquisition of host-derived 5mC methylation across most of the remaining GEVE sequence, consistent with epigenetic silencing. Because the integrated GEVE remained largely unmethylated, we also assessed the methylation status of DNA in punuivirus virions using Enzymatic Methyl-seq (EM-seq). Viral reads were strongly enriched relative to residual host DNA (see **Appendix**), confirming sample purity, and the viral genome showed 5mC levels close to the assay’s non-conversion background (**Figure 5B**), indicating that punuivirus virions are essentially 5mC unmethylated and that 5mC gain in GEVE− is host derived.

**Figure 5.**
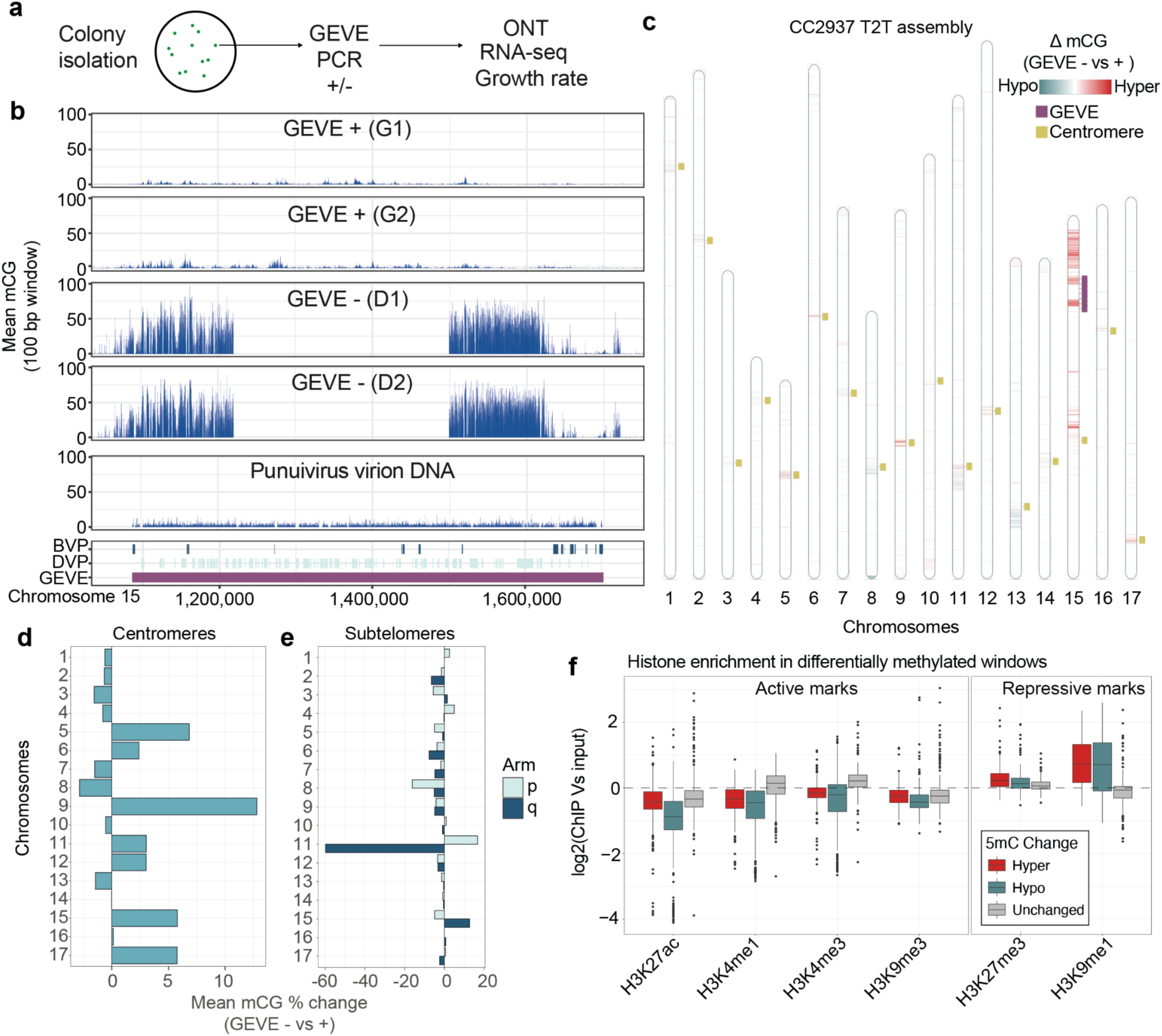
Loss of the full-length GEVE is associated with widespread redistribution of 5mC across the host genome. (A) Experimental strategy for isolating clonal GEVE+ (full-length GEVE) and GEVE− (deletion GEVE) strains by colony PCR, followed by Oxford Nanopore sequencing, RNA sequencing, and phenotypic characterization. (B) Mean CG methylation(mCG) across the GEVE region in two independent GEVE+ and two independent GEVE− isolates, together with purified punuivirus virion DNA measured by EM-seq. Deletion-containing isolates exhibit elevated 5mC across the remaining GEVE sequence, whereas full-length GEVE isolates and viral DNA remain largely unmethylated. (C) Genome-wide distribution of differential CG methylation between GEVE− and GEVE+ strains in 10-kb windows. Red indicates increased methylation in GEVE− strains and blue decreased methylation. Centromeres are indicated in yellow and the chromosome 15 GEVE locus in purple. (D,E) Mean changes in CG methylation across centromeric (D) and subtelomeric (E) regions of each chromosome between GEVE− and GEVE+ strains (p is the short arm, q the long arm). (F) Distribution of histone modification enrichment scores for genomic regions that gain or lose CpG methylation in GEVE− strains. Differentially methylated regions are preferentially associated with heterochromatic chromatin states.

Remarkably, the epigenetic effects of the deletion extended well beyond the GEVE itself. Genome-wide comparison of GEVE− relative to GEVE+ isolates identified substantially more genomic windows that gained 5mC (n=820) than lost it (n=151). Surprisingly, the strongest increase occurred on chromosome 15, in the same chromosome arm where the GEVE insertion site lies (**Figure 5C; Figure S4A, B**). However, methylation changes were also detected throughout the genome. Centromeric regions, which in *C. reinhardtii* are defined by centrophilic *ZeppL* LINE retrotransposons^38,39^, generally gained 5mC methylation in GEVE− strains (**Figure S4C**,**D**), although two centromeres flanked by large satellite arrays instead showed local methylation losses (**Figure S4E**). By contrast, the satellite-enriched subtelomeric regions displayed an overall reduction in 5mC methylation (**Figure 5E**). Together, these observations indicate that loss of the full-length GEVE is associated with widespread redistribution, rather than a simple global increase, of host DNA methylation in GEVE– strains.

To determine which chromatin environments were most susceptible to this epigenetic reconfiguration, we integrated our methylation maps with publicly available histone modification ChIP-seq datasets from *C. reinhardtii* ^40,41^. Despite originating from different strains (CC-5325), sequence conservation allowed reliable comparison across most of the genome beyond the GEVE. Regions undergoing differential 5mC methylation were preferentially associated with heterochromatic histone modifications, particularly H3K27me3 and H3K9me1, whereas promoter-associated H3K4me3 and other euchromatic marks were depleted (**Figure 5F**). These observations suggest that the existing chromatin state strongly influences where 5mC can be gained or lost following GEVE deletion.

In contrast to the extensive redistribution of 5mC methylation, 6mA remained remarkably stable between GEVE+ and GEVE− strains (**Figure S5A**), both within the GEVE and across the host genome.

Furthermore, regions carrying promoter-associated 6mA were strikingly resistant to 5mC gain. Within the GEVE, windows containing 6mA accumulated little or no additional 5mC in GEVE− strains (**Figure S5B**), a pattern that was particularly evident at the transcription start sites of latency-associated genes, where narrow 6mA peaks remained protected despite extensive methylation of the surrounding sequence (**Figure S5C,D**). A similar pattern was observed in the second, highly degenerated 48 Kbp GEVE relict in chromosome 16, whose overall hypermethylated 5mC state was unchanged between GEVE+ and GEVE− strains. However, the relict retained local depletion of 5mC around residual 6mA-marked genes (**Figure S5D**). Together, these results suggest that 6mA-associated euchromatin is refractory to 5mC deposition, restricting the spread of host silencing into transcriptionally permissive regions.

### Partial GEVE deletion remodels host transcription and transposable element activity

To determine whether the extensive redistribution of 5mC was accompanied by transcriptional changes, we performed RNA-seq on all four clonal isolates at two stages of culture growth (3 and 5 days post-inoculation), encompassing the period of viral genome replication (**Figure S7A**). Subsequent detection of extracellular virions by PCR and flow cytometry confirmed that both GEVE+ strains underwent viral induction (**Figure S7A-C**). Within the GEVE, most differentially expressed genes were downregulated in GEVE− strains (10 downregulated versus 3 upregulated; **Figure 6A**). Eight of the ten repressed genes also gained 5mC, consistent with epigenetic silencing. In contrast, many genes belonging to the late viral production module remained silenced or expressed at very low levels in both genotypes and accumulated 5mC, suggesting that 5mC preferentially accumulates at inactive loci (**Figure S6B**). By comparison, 6mA-marked GEVE genes maintained similar expression levels in both strains, consistent with their resistance to 5mC (**Figure S6B**).

**Figure 6.**
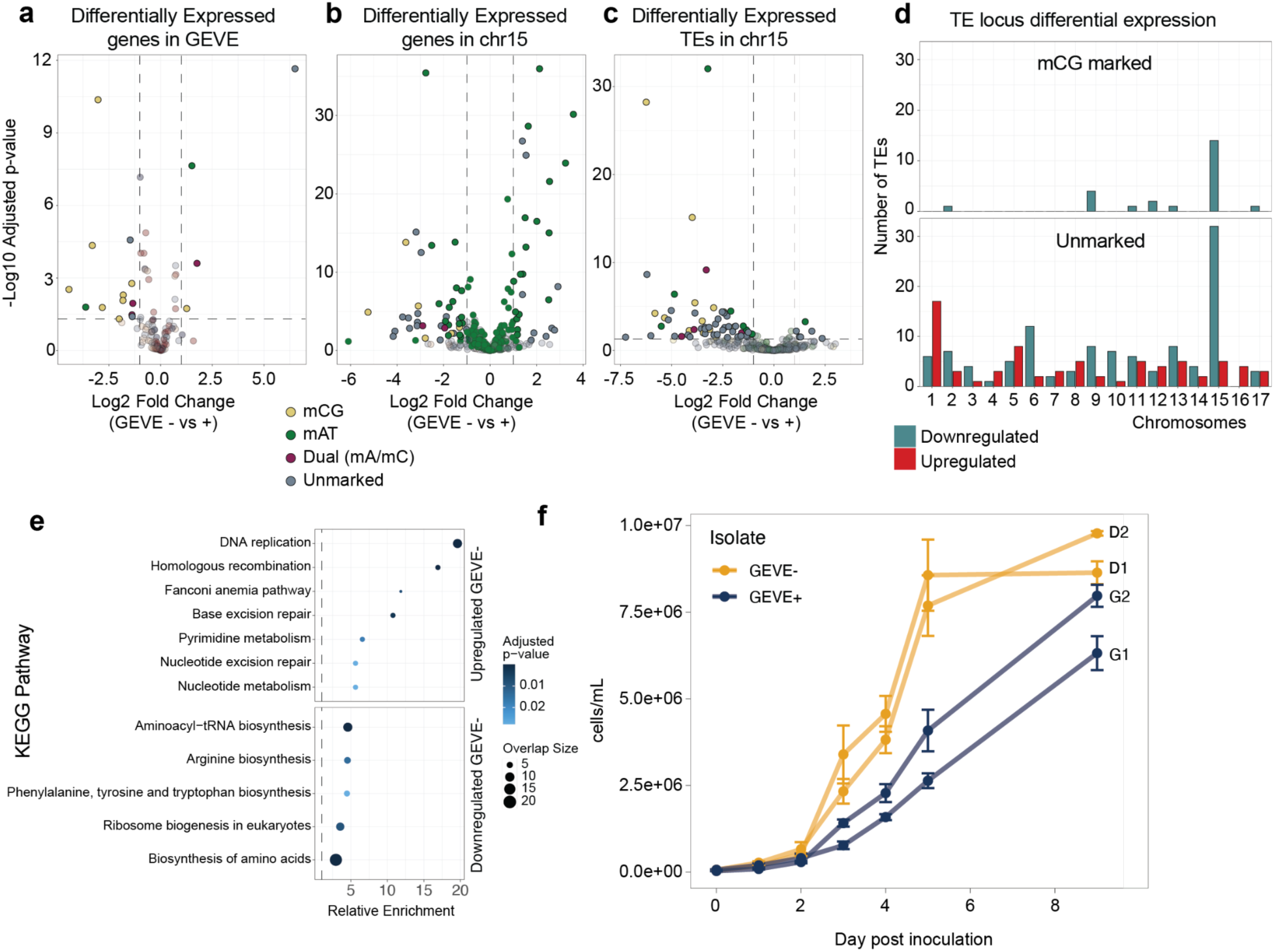
Partial GEVE deletion is associated with transcriptional and physiological remodeling of the host. (A) Volcano plot showing differential expression of GEVE genes between GEVE− (deletion) and GEVE+ (full-length) strains (DESeq2, adjusted P < 0.05). Genes are coloured according to DNA methylation status (mCG-marked, mAT-marked, dual-modified, or unmodified). Genes carrying mCG are predominantly found among the downregulated fraction, consistent with transcriptional repression by CpG methylation. (B) Differentially expressed protein-coding genes along chromosome 15, which harbours the GEVE locus. Downregulated genes include relatively few mCG-marked loci, indicating that transcriptional changes extend beyond genes directly targeted by DNA methylation. (C) Differentially expressed transposable elements along chromosome 15, the majority of which are transcriptionally repressed in GEVE− strains. (D) Number of differentially expressed transposable element insertions on each chromosome, stratified by CpG methylation status and direction of expression change. Transposable elements that acquire mCG are exclusively downregulated, whereas numerous unmethylated transposable elements on chromosome 15 are also repressed, indicating that transcriptional effects are not fully explained by local DNA methylation. (E) KEGG pathway enrichment analysis of differentially expressed host genes. A total of 971 upregulated genes (147 annotated) and 978 downregulated genes (218 annotated) were included in the analysis. (F) Growth curve of GEVE+ and GEVE-strains in liquid media, showing faster exponential growth in GEVE-strains.

The transcriptional effects were not restricted to the GEVE. Chromosome 15, which undergoes the greatest increase in 5mC following GEVE deletion, contained eight host genes that both gained 5mC and became transcriptionally repressed (**Figure 6B**). Only one additional gene elsewhere in the genome showed the same relationship (**Figure S6C**). Nevertheless, many host genes were differentially expressed despite showing no detectable changes in DNA methylation (**Figure 6B; Figure S6C**), indicating that much of the transcriptional response is likely not directly linked to 5mC changes. This is also consistent with the observation that most 5mC changes occur within heterochromatic and therefore gene-poor regions of the genome.

The strongest transcriptional effects were observed in transposable elements (TEs). In the GEVE– strains, every TE that acquired 5mC became transcriptionally repressed (n=24), both on chromosome 15 and elsewhere in the genome (**Figure 6C,D**). However, repression was not exclusively linked to 5mC, as numerous unmethylated TE insertions (particularly on chromosome 15) were also downregulated (**Figure 6D**). At the family level, *ZeppL* retrotransposons accounted for much of the genome-wide gain in 5mC, reflecting their exclusive presence within centromeric regions (**Figure S8A**). However, differential expression analysis using the TEtranscripts tool detected little change in *ZeppL* transcript abundance, while they were consistently among the most highly expressed TE subfamilies (**Figure S8B,C**). Instead, the largest transcriptional changes were observed among Helitron and Gypsy transposable elements (**Figure S8B**), without a clear directionality. Together, these results suggest that the presence of the intact GEVE promotes a more transcriptionally permissive environment for a subset of TEs, particularly in chromosome 15, whereas partial deletion of the GEVE is accompanied by widespread TE repression through both DNA methylation-dependent and -independent mechanisms.

Beyond TEs, GEVE deletion was also associated with broader changes in host physiology. KEGG pathway enrichment analysis of differentially expressed protein-coding genes showed that GEVE− strains preferentially upregulated pathways involved in mitosis, DNA replication, and DNA repair (**Figure 6E**), consistent with their faster growth in both liquid and solid media (**Figure 6F and Figure S7D**). Conversely, genes upregulated in GEVE+ strains were enriched for metabolic pathways, which may reflect either metabolic reprogramming that favors viral production or an indirect consequence of reduced investment in cell proliferation. Notably, despite these transcriptional and growth differences, GEVE+ strains do not undergo widespread viral production and cultures still reach stationary phase (**Figure 6F**), indicating that maintenance of an unmethylated, transcriptionally permissive GEVE is not sufficient to trigger large-scale GEVE induction. Additional regulatory mechanisms or environmental cues must therefore govern the transition from latency to productive infection.

### A viral TET dioxygenase provides a candidate mechanism for maintaining an unmethylated GEVE

The widespread acquisition of 5mC following GEVE deletion suggested that the missing genomic region encodes a factor that actively protects the integrated virus from host epigenetic silencing. We therefore searched the deleted interval for genes with potential chromatin regulatory functions and identified a Ten-Eleven Translocation (TET)-like dioxygenase (GEVEv2_173). TET enzymes are the principal mediators of active DNA demethylation in animals^42^, and a recently described TET homolog in *C. reinhardtii* (CMD1) catalyzes a rare form of 5mC oxidation in this alga^43^. The viral TET gene contains eight introns and 6mA marks near the TSS (**Figure 7A**), consistent with host-mediated regulation during periods of latency. It is among the most highly expressed punuivirus genes overall, ranked 33 and 32 across bulk RNA-seq and scRNA-seq datasets, respectively (**Figure 7B**), and clusters within the constitutively expressed BVP module (see **Appendix**). Importantly, the ∼280-kbp deletion encompasses the TET gene and only 5 other constitutively expressed BVP genes located within the central region of the GEVE. Consequently, GEVE− strains completely lack TET expression. We therefore propose that continuous expression of this viral enzyme counteracts host 5mC deposition during latency, maintaining the integrated GEVE in a transcriptionally permissive state (**Figure 7D**). Loss of the TET-containing region would then allow host DNA methyltransferases to rapidly methylate the remaining viral genome, explaining the extensive 5mC hypermethylation observed in GEVE− strains, yet having no effect on 6mA.

**Figure 7.**
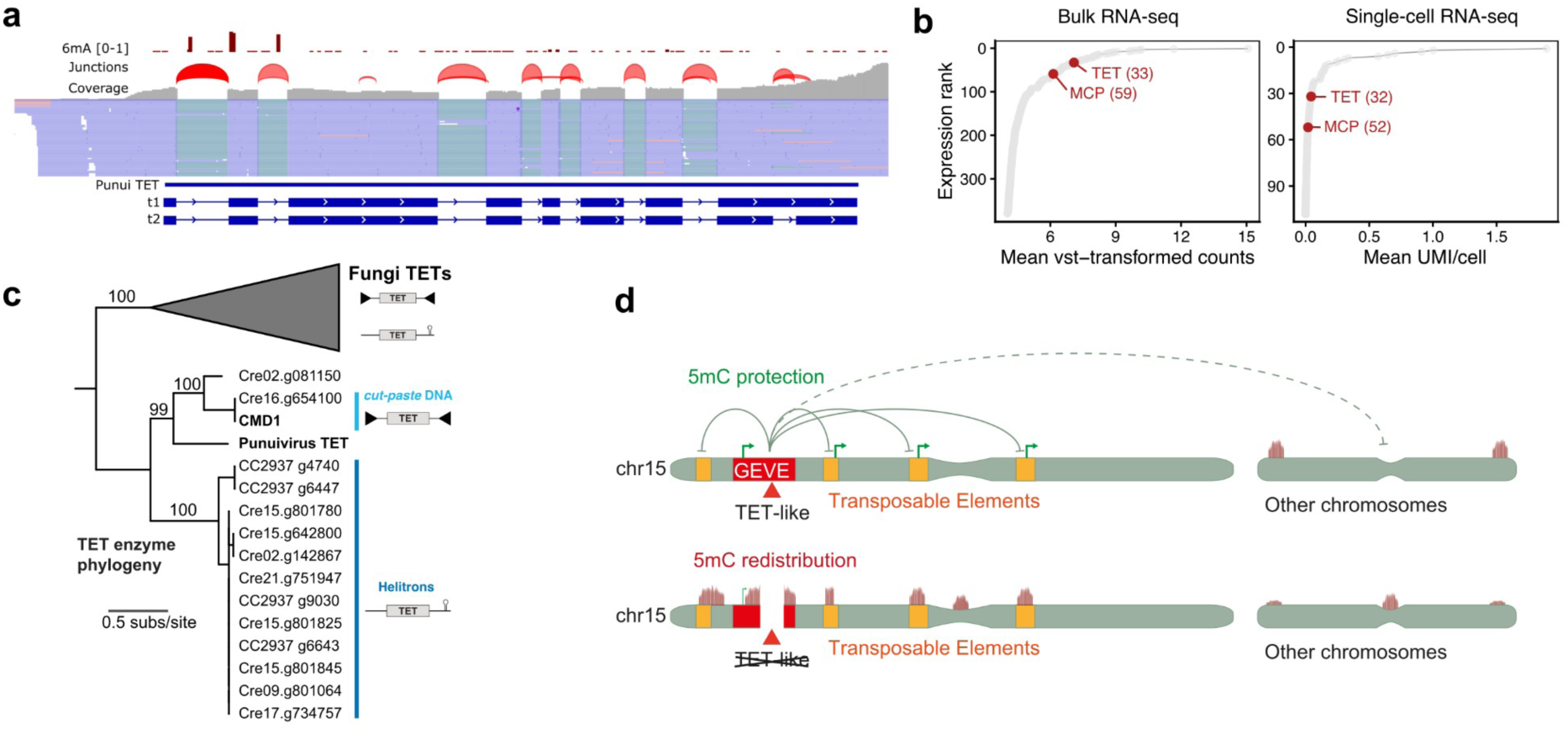
A GEVE-encoded TET enzyme is a potential candidate for epigenetic manipulation. (A) Punuivirus GEVEv2_173 Ten-Eleven Translocation (TET) gene model predicted by BRAKER3, supported by RNA-seq mapped reads. Splice junctions were confirmed by Portcullis (see Methods). (B) Mean viral TET expression across all samples in bulk RNA-seq (left, vst-transformed) and single-cell RNA-seq (right, log1p transformed). Values were ranked using the dense ranking method. Mean expression of the major capsid protein is shown for comparison. (C) A phylogenetic tree of TET proteins, highlighting the *C. reinhardtii* clade. Fungi outgroups obtained from Iyer et al.^45^ and rooted following ^44^. Sequences found within TEs are highlighted, note that in fungi TETs are carried by multiple different groups of *cut-paste* DNA transposons ^45,46^. Proteins with Cre IDs belong to the reference genome strain CC-4532. Transposon pictograms are simplified and all elements in *C. reinhardtii* carry multiple genes (see Appendix). (D) Proposed model for the epigenetic consequences of GEVE deletion. In full-length GEVE strains, the viral TET dioxygenase maintains a hypomethylated GEVE and limits 5mC at other genomic regions. Loss of the TET-containing genomic region permits 5mC to accumulate at the remaining GEVE while redistributing methylation across the host genome, leading to widespread changes in transposable element and host gene expression.

We next investigated the evolutionary origin of the viral TET relative to other *C. reinhardtii* TET genes. *CMD1* is one of eleven TET homologs in the *C. reinhardtii* reference genome (strain CC-4532). We found clear support that 10 of 11 TET genes are carried by two specific TEs: two copies of *CMD1* are encoded by a large cut-and-paste DNA transposon, whereas 8 other homologs are part of a Helitron (**Figure 7C, Appendix**). Furthermore, all 11 of these TET genes are missing from the CC-2937 genome, suggesting that they segregate as presence-absence polymorphisms in the species. Supporting this, we detected four distinct TET copies in CC-2937 that are carried by polymorphic Helitron insertions. Thus, it appears that the situation in *C. reinhardtii* mirrors that described for the related TET genes of fungi^44^, which are carried as cargo by a vast array of DNA transposons^45,46^. Intriguingly, the viral enzyme is phylogenetically placed within the diversity of the *C. reinhardtii* DNA transposon TETs (**Figure 7C**), suggesting that punuivirus may have captured the TET gene from a transposon present in its host. Importantly, the punuivirus TET is also closely related to the functionally characterized CMD1 (60% amino acid identity in the core TET-JBP domain and conservation of all functionally characterized sites^47^) (see **Appendix**), further supporting its potential role as a catalytically active enzyme capable of DNA demethylation.

Collectively, these findings suggest that the constitutively-expressed viral TET is a key component of the latent infection program. Rather than allowing the host to progressively silence the integrated virus, punuivirus appears to maintain a dynamic equilibrium in which viral demethylation continuously counteracts host epigenetic defense (**Figure 7D**). If similar chromatin regulators are widespread among giant endogenous viruses, modulation of host epigenetic landscapes may represent a general strategy by which these viruses achieve long-term coexistence within eukaryotic genomes.

### Conclusions

With a 617 kbp genome, punuivirus is the largest latent virus discovered to date, and our results highlight the dramatic impact of its cryptic infection cycle on both its genome evolution as well as the physiology and epigenomics of its host. Our refined genome assembly and re-annotation demonstrates that punuivirus has a wide range of latency-related genes that are expressed during periods in which no viral production can be detected. We show that these latency genes have acquired numerous spliceosomal introns, and they are commonly associated with TSS-associated 6mA marks, mirroring the genetic architecture and regulatory themes that define the host green alga, where these elements play key roles in gene expression^24,25^. Historically, large DNA viruses have been thought to be intron-poor, and only recently have metagenomic surveys recovered a broader diversity of intron-rich viral lineages^48^. Indeed, widespread introns were previously identified in GEVEs, but they were considered to be signatures of genomic erosion in inactivated viral relics rather than components of a functional genetic machinery^6^. Relatives of punuivirus are found throughout a variety of green algae^6^, and the broader punuivirus lineage therefore represents a valuable system for examining the process of viral intron acquisition as well as the long-term consequences of maintaining diverse, intron-rich genes on host range and persistence. In addition, although diverse epigenetic marks have been found in the genomes of virulent giant viruses^49,50^, research in this area has focused mainly on virus encoded restriction modification systems, but our finding of 6mA marks linked with latency gene expression thereby expands the breadth of roles these epigenetic marks play during infection. While punuivirus lacks a 6mA methyltransferase, its host-like regulatory architecture may allow viral genes to recruit the host transcriptional machinery and, in doing so, co-opt the host 6mA pathway.

By resolving the punuivirus expression program at single-cell resolution, we were able to further demarcate the transcriptional modules involved in both latency and virulence, and quantify the number of virocells present at different stages of culture growth. Punuivirus expresses mainly latency-related transcripts in most algal cells, and a distinct transcriptional shift to the virulence module occurs in only a small subpopulation of virocells. The low number of virocells we detect is consistent with microscopy-based observations of punuivirus relatives that have found at most 3% of cells had detectable intracellular virions^16^. Interestingly, early electron microscopy studies of green algae also reported few or even single cells with intracellular virions among hundreds surveyed^51^. At the time these initial studies were published, it remained unclear why only a small number of cells would exhibit signatures of infection in an otherwise healthy culture, but the presence of widespread cryptic infections now appears likely. Our observation of widespread expression of viral latency genes in the majority of the algal population also highlights the impact of harboring a GEVE even without viral production. Indeed, our results stand in stark contrast to latency in other eukaryotic DNA viruses, such as herpesviruses, where few or no transcripts are produced during periods of latency^52^. For example, in human cytomegalovirus, latency appears to be regulated simply through muted expression of virulence genes^53^.

Although the majority of cells in a population do not experience the switch to virulent infection, our epigenetic analysis demonstrates that simply carrying a full-length punuivirus GEVE has a dramatic impact on the epigenetic status of the host genome as well as overall host physiology. Indeed, hosts carrying a partially-deleted GEVE exhibit a dramatic remodeling of their genome-wide 5mC profile, including altered methylation of transposable elements, centromeres and subtelomeric regions. We propose that these changes may reflect an epigenetic sink effect in which the intact GEVE alters the distribution or activity of host methylation machinery, and its disruption consequently rebalances 5mC across genomic compartments^54^. How the viral TET-like gene exerts such chromosome-scale effects remains unclear, as the protein lacks an obvious sequence-specific DNA-binding domain and, following translation in the cytoplasm, would need to return to the nucleus and preferentially act on the GEVE in chromosome 15. TET-like demethylases are common across diverse endogenous viruses, as recently reported for diverse Polinton-like viruses^55^, suggesting that these host-evading strategies are not limited to punuivirus. Maintaining an unmethylated and transcriptionally permissive GEVE is not sufficient to trigger productive infection, however, as most cells carrying the full-length GEVE do not become virocells. Punuivirus therefore illustrates how long-term viral latency can emerge from a molecular compromise, in which viral persistence depends on integration into host regulatory systems while simultaneously reshaping the cellular environment in which it resides.

Integration into host transcriptional programs may also have consequences beyond viral persistence. By acquiring host-compatible gene architectures and epigenetic features, latency-associated viral genes become primed for functional domestication by the host genome, especially because they remain transcriptionally active even after the GEVE itself begins to degenerate. At the same time, the ability of punuivirus to influence epigenetic regulation far beyond its own genomic boundaries shows that the consequences of latency need not be restricted to the viral sequence itself. Such interactions may therefore be more complex than a simple host–virus arms race. Over millions of years of coexistence, recurrent exchange of genes and regulatory activities between giant viruses and their hosts could become an important source of both genetic and epigenetic innovation in eukaryotic genomes.

## STAR Methods

### Resource Availability

#### Lead Contacts

Further information and requests for resources and reagents should be directed to and will be fulfilled by the lead contacts, Corresponding Authors Alex de Mendoza and Frank O Aylward

## Materials Availability

### Data and Code Availability

All data reported in this paper will be shared upon reasonable request to the lead contact. Nucleotide assembly and annotation files for the resequencing of CC-2937 can be found on Zenodo (https://doi.org/10.5281/zenodo.22310362). Single-cell RNA-seq data (raw and processed) have been deposited in the NCBI Gene Expression Omnibus (GEO) under accession GSE347098; the processed AnnData object of GEVE-expressing cells used for clustering and UMAP analyses is available from the same Zenodo record. Code to reproduce all single-cell transcriptomic analyses, figures and supplementary tables is available at https://github.com/urisheyn/GEVE-single-cell. POD5 files containing raw Oxford Nanopore signal data along with bedMethyl files containing per-base 5mC and 6mA methylation summaries for the samples generated in this study are available from EBI BioStudies^56^ under the accession S-BSST3314. POD5 files for the samples generated in Erazo-Garcia et al.^16^ are available from SquiDBase^57^ under the accessions SQB000017, SQB000018 and SQB000019. Bulk RNA-seq raw sequencing data for the full-GEVE and GEVE-deletion isolates have been deposited in the NCBI Sequence Read Archive (SRA) under BioProject PRJNA1528683, which links both bulk RNA-seq and the single-cell transcriptome datasets. ONT data used to generate the gapless CC-2937 assembly have been deposited in the NCBI SRA under BioProject PRJNA1531727. Processed DNA methylation data for all samples along with code to reproduce DNA methylation analyses are available at https://github.com/AlexdeMendoza/Chlamydomonas_GEVE. The *C. reinhardtii* subclones described in this study are available at the Chlamydomonas Resource Center. Any additional information required to reanalyze the data reported in this paper is available from the lead contact upon request.

### Method Details

#### Punuivirus GEVE gene model reannotation

RNA-seq reads generated by Erazo-Garcia et al.^16^ (from all samples, one replicate) were combined and mapped to the existing *C. reinhardtii* CC-2937 genome using STAR v2.7.9a^58^ in two-pass mode with the parameters “–alignIntronMax 5000” and “--outSAMstrandField intronMotif”. The resulting BAM file was sorted and indexed with SAMtools v1.22.1^59^ and used to refine the gene annotations within the GEVE region using BRAKER3^60^. A curated protein database containing *Nucleocytoviricota* and *C. reinhardtii* CC-4532 v6.1 protein sequences was also incorporated to generate intron-aware gene models. These predictions were merged with the previous Prodigal-based annotation, using BRAKER3 gene models as the primary reference. The resulting unified annotation comprised 506 genes, each assigned an identifier with the prefix “GEVEv2”. Gene models were transferred to the 48 kbp and 5 kbp GEVE relicts using Liftoff v1.6.3 followed by manual curation. RNA-seq reads were subsequently remapped to the updated GEVE transcript annotation, and the differential expression analysis was repeated as described previously^16^. Viral genes were reclassified according to their expression profile into clusters of genes enriched Before Viral Production (BVP) or During Viral Production (DVP).

### GEVE intron characterization

A total of 61 genes were predicted to carry spliceosomal introns using BRAKER3. In addition, a group IC1 self-splicing intron was annotated within the MCP gene (GEVEv2_329) using Infernal v1.1.5^61^ and covariance models for group I/II introns^62^. Because it is not a host-type spliceosomal intron, the MCP gene was excluded from all intron-based analyses. Intron splice junctions were also manually curated in IGV and validated using Portcullis. To examine the nucleotide composition surrounding the 3′ splice site, intron sequences of at least 100 bp were aligned relative to their 3′ end, and nucleotide frequencies were calculated using a sliding window of 25 bp. Splice sites were visualized using WebLogo3 3.9.0^63^ after aligning intron 5’ and 3’ ends.

### Protein functional annotation

In addition to annotations methods used previously, protein sequences were annotated by generating multiple sequence alignments (MSAs) using HHBlits^64^ and the UniRef30 database (release 2023/02) with three iterations, a minimum coverage of 20% and an E-value threshold of 1x10^-3^. The resulting MSAs were used to search against the PDB70 database using HHsearch^64^, and only hits with a probability of at least 80% were retained for functional annotation.

### Promoter motif analysis

We searched for putative promoter motifs 6 to 12 bp in length within the GEVE by retrieving the 200 bp upstream region of each predicted gene and performing motif discovery with STREME v5.5.9^65^ using a p-value threshold of 0.05. A Markov background model was generated with the MEME Suite function fasta-get-markov using GEVE intergenic regions of at least 50 bp. Intergenic regions located between convergently transcribed genes were excluded from the background model generation, resulting in the following nucleotide frequencies: A/T = 0.219 and G/C = 0.281. GEVE genes located within repetitive regions were also excluded from the analysis (GEVEv2_205–209, GEVEv2_235–246, and GEVEv2_457– 463). The best motif was then searched using FIMO v5.5.9^66^ against the upstream sequences, yielding 111 motif occurrences (q-value <0.05) in a total of 105 genes.

### Re-assembly of CC-2937 genome and repeat annotation

The gapless genome assembly was produced using a subclone of CC-2937 that is being used for the Chlamydomonas Pan-Genome Project, which was obtained from the Chlamydomonas Resource Center as CC-5826. This subclone originally had an intact GEVE (determined by PacBio sequencing, unpublished data) but must have experienced a unique GEVE deletion event prior to ONT sequencing.

High molecular weight DNA for genome assembly was extracted from a 75 ml liquid TAP culture maintained under constant light (90 μmol m-2 s-1) with shaking at 160 rpm for 3d. Extraction was performed using the Macherey-Nagel NucleoBond HMW DNA kit, following the supplier’s instructions with the exception of cell lysis, which was performed at 65 °C for 30m in 2.5 ml of the supplied lysis buffer supplemented with 1.4 ml 5M NaCl, 1 ml CTAB 10% / PVP 40,000 5% and 50 μL 0.5M EDTA. A standard 1X ratio AMPure XP Bead clean-up was performed prior to ONT library preparation.

High molecular weight genomic DNA was subject to ONT sequencing on a P2 Solo using the Native Barcoding V14 library prep (SQK-NBD114.24) and an R10.4.1 flow cell. Adaptive sampling was performed to deplete organelle reads by providing assemblies of the *C. reinhardtii* plastome and mitogenome. Basecalling was performed using Dorado v1.4.0 and the dna_r10.4.1_e8.2_400bps_sup@v5.2.0 model. Assembly was performed using hifiasm v0.24.0-r703 (-l0 – ont) ^67^ after filtering out reads of less than 10 kb, which resulted in ∼6.4 Gb of data. The assembly yielded 17 megabase-scale contigs that were confirmed to correspond to chromosomes by manual comparison to the *C. reinhardtii* reference genome^35^. 19 small contigs containing redundant ribosomal DNA arrays, subtelomeric repeats (see ref. 36) or organellar sequence were removed. Inspection of chromosome 15 revealed that this subclone harbored a complete GEVE deletion (see below), and the existing GEVE assembly was therefore manually inserted to restore the ancestral viral integration. The final assembly consists of 17 gapless chromosomes, all of which terminate in either telomeres (n=25), rDNA arrays (n=2) or subtelomeric repeats (n=7).

The gene and GEVE annotations were transferred to the new gapless assembly by using Liftoff^68^ (version 1.6.3), using the `-polish` option for genes. Repeat annotation was performed by passing the curated library of *C. reinhardtii* repeats^38^ to RepeatMasker v4.0.9 (https://www.repeatmasker.org/). Satellites identified from the curated library were merged with automated prediction using Tandem Repeats Finder v4.09.1^69^ run with the options `2 5 7 80 10 50 2000 -d -ngs` and filtered for repeats with monomer lengths of 10 or more bp. Centromeres were defined as regions spanning from the first to the last *ZeppL* retrotransposon copy^35^, and subtelomeres were defined as any sequence beyond the first nucleotide of the *Sultan* satellite arrays^36^. Individual TE copies were predicted using the One code to find them all Perl script (--unknown --strict)^70^.

### Single-cell RNA sequencing library preparation

#### Cell biomass collection and fixation

*C. reinhardtii* CC-2937 cultures were grown to a density of 5 × 10⁶ cells/mL, and 50 mL aliquots (2.5 × 10⁸ cells) were harvested by centrifugation and fixed using the Evercode Cell Fixation v2 kit (Parse Biosciences, Seattle, WA, USA). The manufacturer’s protocol was used, with the exception that cells were sequentially treated with a prefixation buffer, a fixation solution, and a permeabilization solution prior to barcoding. Immediately following permeabilization and prior to neutralization, cells were treated with subtilisin (Alcalase; derived from *Bacillus licheniformis*) at 3 Anson units/mL, the highest concentration tested by Hwang et al.^71^ who showed this broad-specificity serine protease effectively disrupts the algal cell wall of *C. reinhardtii*. Cells were then neutralized, resuspended in DMSO-supplemented cell buffer, and stored at −80°C.

#### Single-cell combinatorial barcoding

Fixed cells were processed using the Evercode WT Mini v2 kit (Parse Biosciences) according to the manufacturer’s protocol. Briefly, cells were counted and distributed across 12 wells of a Round 1 barcoding plate, where a well-specific barcode was introduced by an in-cell reverse transcription reaction using barcoded primers. Cells were pooled and distributed across two further 96-well barcoding rounds, in which a second and third well-specific barcode were appended to each transcript via sequential in-cell ligation reactions. After the third round of barcoding, cells were pooled, strained, and split into sublibraries prior to lysis.

#### cDNA amplification

Barcoded cDNA was captured on streptavidin binder beads via a biotin moiety introduced during the third barcoding round. Bead-bound cDNA underwent a template-switching reaction to append a primer-binding site to the 3′ end, followed by PCR amplification (5 initial cycles plus the maximum number of second-cycle repeats specified for the corresponding sublibrary cell count, per the manufacturer’s guidelines) to generate double-stranded, whole-transcriptome amplified cDNA. Amplified cDNA was purified by SPRI bead cleanup, and concentration and fragment size distribution were assessed by fluorometric quantification and capillary electrophoresis.

#### Sequencing library preparation

For each sublibrary, 100 ng of amplified cDNA (or the entire yield when below 100 ng) was fragmented, end-repaired, and A-tailed, followed by a double-sided SPRI size selection. Illumina sequencing adaptors were ligated to the fragmented cDNA, and a second SPRI cleanup was performed. A sublibrary index PCR then appended the fourth, sublibrary-level barcode together with the P5 and P7 Illumina adaptors, with the number of PCR cycles adjusted according to the amount of cDNA input into fragmentation. Final libraries underwent a double-sided SPRI size selection targeting a fragment size of 400 to 500 bp and were quantified by fluorometric and capillary electrophoresis assays.

#### Sequencing

Sublibraries were pooled and sequenced on an Illumina NovaSeq 6000 instrument with paired-end reads (Read 1 = 74 cycles, i7 index = 6 cycles, Read 2 = 86 cycles) and a 5% PhiX spike-in, per the manufacturer’s recommended run configuration. Sublibraries were demultiplexed on their 6-bp i7 sublibrary index.

#### Algal sub-strain isolation

*Chlamydomonas reinhardtii* CC-2937 was obtained from the Chlamydomonas Resource Center and maintained on Tris-Acetate-Phosphate (TAP) agar slants supplemented with yeast extract (4 g L^-1^) at 23 °C under 120 μmol m^-2^ s^-1^ light intensity. We evaluated the punuivirus GEVE integrity in individual colonies using a PCR assay targeting the MCP^72^ as well as a previously defined ∼280–kilo–base pair (kbp) internal deletion spanning the MCP locus. This deletion was characterized using long-read sequencing data from our previous study^16^ (SRA accession: SRS21938379) and detected using the primers GEVE_DEL_Fw (5′– GTT GCA GAC ACC ATT GGG–3′) and GEVE_DEL_Rv (5′–ACA GCT CAG CAA GAG GC–3′) which flank the deletion boundaries. Conditions for amplification can be found in Moniruzzaman et al^72^. Two GEVE+ (mcp-positive) and two GEVE-colonies (deletion-positive) were selected and restreaked through three rounds of single-colony isolation to ensure purity. The GEVE+ and GEVE-monoclonal isolates were designated G1 and G2, and D1 and D2, respectively, and were inoculated onto slants for long-term maintenance.

#### DNA extraction of algal cells and viral particles

Liquid cultures of GEVE+ and GEVE-strains were started from cells grown on TAP agar plates and transferred into 100 mL of TAP medium in Erlenmeyer flasks. Flasks were incubated for ten days under a 12:12 h light:dark photoperiod, agitated in an orbital shaker at 150 rpm, and exposed to 100 μmol m-2 s-1 light intensity. These cultures were then used to inoculate 100 mL TAP medium at a starting density of 5 x 10^4^ cells mL^-1^. After five days of incubation under the same conditions, 50 mL of culture was harvested and centrifuged at 4,500 g for 1 h at 4 °C. The resulting biomass was used for DNA extraction as previously described^16^.

The remaining supernatant from the GEVE+ cultures was used for viral particle isolation. Approximately 400 mL of supernatant was filtered through a 5 μm PES membrane and concentrated to ∼ 20 mL by tangential flow filtration using a 30 kDa PES Vivaflow 200 TFF unit. The retentate was distributed into microfuge tubes and centrifuged at 20,000g for 2 h at 4 °C. Pellets were resuspended in 50 μL molecular-grade water and pooled into a single sample, which was passed through a 0.8 μm PES syringe filter. Following the methods described in Erazo-Garcia et al.^16^, viral-like particles were stained and visualized by flow cytometry. Unencapsidated cellular DNA was then digested by treating the sample with 6U of DNase for 1 h at 37 °C prior to viral DNA extraction^16^.

#### Enzymatic methyl-seq library preparation and analysis

4 ng of punuivirus DNA was spiked with unmethylated lambda phage DNA and a pUC19 CpG-methylated plasmid to assess unmethylated cytosine conversion efficiency and enzymatic protection of methylated cytosines, respectively. DNA was sheared with the NEBNext UltraShear enzyme (NEB, no. M7634) at 37°C for 10 minutes and a paired-end EM-seq library was constructed using the NEBNext Enzymatic Methyl-seq v2 Kit (NEB, no. E8015) following the manufacturer’s instructions. The resulting library was sequenced on an Illumina NovaSeq X Plus platform.

The EM-seq converted reads were trimmed using fastp^73^ (version 0.22.0) and overlapping read pairs were merged using BBMerge^74^ (BBTools version 38.18) and treated as single-end reads. Merged and unmerged reads were then mapped to the gapless CC2937 genome assembly with pUC19 and lambda phage DNA sequences appended (downloaded from https://raw.githubusercontent.com/nebiolabs/EM-seq/refs/heads/master/assets/methylation_controls.fa) using BS-Seeker2^75^ (version 2.1.7) with Bowtie 2 (version 2.2.8) as the backend aligner in end-to-end alignment mode. Single-end merged reads and paired-end unmerged reads separately. The BS-Seeker2 parameters `-e 300 -X 3000 -m 4` were used to align unmerged reads, whereas `-e 400 -m 6` were used for merged reads. Overlapping portions of read pairs in the BAM file for the unmerged reads were clipped using bamUtil clipOverlap^76^ (version 1.0.15). BAM files for merged and unmerged reads were then merged and deduplicated using Sambamba^77^ (version 0.6.6) and methylation calling was subsequently performed with CGmapTools^78^ (version 0.1.3). GEVE CpG sites had a mean coverage of 48× and a mean methylation level of 3.1%, with phage lambda cytosines and pUC19 CpG sites showing mean methylation levels of 3.2% and 98.7%, respectively. The resulting CGmap file was filtered for CpG sites and processed using R.

#### Oxford Nanopore sequencing and methylation calling

Oxford Nanopore sequencing libraries were prepared for the clonal GEVE+ and GEVE− lines generated for this study from 1 µg of input DNA per sample using the SQK-LSK114 Ligation Sequencing Kit, following the manufacturer’s instructions. Libraries were sequenced on a PromethION sequencer using R10.4.1 chemistry flow cells. Basecalling was performed using Dorado (version 1.2.0) with the sup,5mCG_5hmCG and sup,6mA models for calling 5mC and 6mA modifications, respectively. The resulting reads were aligned to the gapless CC2937 genome assembly using minimap2^79^ (version 2.24-r1122) with the map-ont preset. Nanopore reads for the samples generated in Erazo-Garcia et al.^16^ were re-basecalled from the original POD5 files using Dorado (version 0.9.1) with the hac,5mCG_5hmCG model and then mapped to the CC-2937 T2T genome assembly.

Modkit (version 0.4.1) was used to quantify the frequency of 5mC and 6mA calls at individual CpG and ApT sites across the gapless CC2937 assembly using the `modkit pileup` command. For CpG sites, we used the sample-specific call probability pass threshold automatically calculated by Modkit for 5mC and unmodified cytosine calls, ignoring 5hmC calls and combining calls from both strands at each site (parameters `--cpg --ignore h --combine-strands`). For ApT sites, we used a stringent threshold of 0.995 for 6mA calls and the automatically calculated threshold for unmodified adenine calls, again combining calls from both strands at each site (parameters `--motif AT 0 --mod-threshold a:0.995 --combine-strands`). The resulting bedMethyl files were processed using R.

For analysis of per-read base modifications, the `modkit extract calls` command was used to extract base modification calls for reads aligning to chromosome 15. Calls at CpG sites were filtered to retain those passing the automatically calculated call probability threshold (parameters `--region chromosome_15 --cpg --pass-only --mapped-only`), while calls at ApT sites were filtered to retain those passing a 0.995 probability threshold (parameters `--motif AT 0 --pass-only --mapped-only --filter-threshold A:0.995`). The resulting files were processed in R and filtered to retain calls with a canonical basecall quality score greater than 20. Methylartist^80^ (version 1.5.2) was used to generate plots of modified base calls in individual nanopore reads **(Figure 4C** and **Appendix**).

mAT sites were defined as AT sites with over 50% methylation in both G1 and G2 samples. mAT-marked genes were defined as those with an mAT site located within 400 bp upstream to 1,000 bp downstream of their TSS. Genes gaining mCG in GEVE-samples were defined as those with a mean mCG value over 15 percentage points higher in GEVE– samples than GEVE+ samples across the region extending from 400 bp upstream of their TSS along the full gene body. mAT-marked TEs were defined as those overlapping an mAT site, while TEs gaining mCG in GEVE-samples were defined as those with a mean mCG value over 15 percentage points higher in GEVE– samples than GEVE+ samples. Windows gaining or losing mCG in GEVE− samples were defined as non-overlapping 1 kb windows with mean mCG values differing by more than 15 percentage points between GEVE− and GEVE+ samples.

#### ChIP-seq Analysis

ChIP-seq data from two different studies profiling histone marks in the *C. reinhardtii* strain CC-5325 were downloaded from NCBI (NCBI BioProject accessions PRJNA255778 and PRJNA1029152). Reads were aligned to the gapless CC2937 genome assembly with Bowtie 2 (version 2.5.4) in end-to-end alignment mode with the `--very-sensitive` preset and allowing a maximum of one alignment per read (`-k 1`). The resulting BAM files were converted to bigWig files using bamCoverage (deepTools^81^ version 3.5.6), normalizing read depth using counts per million (CPM). Histone mark signal was then normalized relative to input controls using bigwigCompare (deepTools) with the parameters `--operation log2 --pseudocount 1 --skipZeroOverZero --binSize 50`. The UCSC Genome Browser utility bigWigAverageOverBed (version 2) was used to calculate the ChIP-seq signal in non-overlapping 1 kb windows, with non-covered bases counted as zeroes.

#### Temporal analysis of viral production and RNA-seq in algal strains

Cells from each GEVE+ and GEVE– strain were transferred from slants into 100 mL of TAP medium and grown for five days. These cultures were then used to inoculate 100 mL TAP medium, in duplicate, at a starting density of 5 x 10^4^ cells mL^-1^. Cultures were incubated following the conditions described before, and cell counts were recorded by flow cytometry using the CytoFLEX-S Flow Cytometer^16^. Between three (early exponential) and five (late exponential) days post-inoculation, cell biomass and supernatant were collected by centrifuging 1.5 mL aliquots at 2,500g for 5 min. Two biomass aliquots per sample were flash-frozen with liquid nitrogen and stored at -80 °C for subsequent RNA and DNA extraction. The supernatant was filtered through a 0.45 μm PES syringe filter and treated with DNase prior to PCR detection of viral particles using viral (MCP) and host (ITS1-5.8S-ITS2) markers as previously described^16^. An additional supernatant aliquot was collected on day ten, as viral particles were not yet detected at or before day five using PCR.

Total DNA was extracted from the GEVE+ strains cell biomass by scaling down the extraction protocol used previously. Briefly, 350 μL of preheated SDS buffer (50 mM Tris-HCl pH 8, 200 mM NaCl, 20 mM EDTA, 2% SDS) and 350 μL of CTAB buffer (100 mM Tris-HCl pH 8, 20 mM EDTA pH 8, 1.4 M NaCl, 2% CTAB, 1% PVP M.W. 40,000) were added to each sample along with 5μL Proteinase K (20 mg ml−1), followed by incubation at 65°C for 30 min. RNase A (5μL; 100 mg mL−1) was then added and samples were incubated for an additional 10 min at room temperature. Samples were centrifuged at 4,500 g for 5 min, and the supernatant was extracted once with an equal volume of phenol-chloroform (1:1), followed by one extraction with one volume of chloroform. DNA was precipitated with 1/10 volume 3M sodium acetate and two volumes of 100% ethanol. Samples were incubated at -20 °C for 1 h, and the DNA was pelleted by centrifugation at 20,000g for 30 min at 4 °C. Pellets were rinsed with 70% ethanol and resuspended in 40 ul of molecular-grade water. Samples were further diluted to ∼ 10 ng μL-1 prior to qPCR.

To assess viral genome replication, qPCR was performed in three technical replicates using 2 μL of algal biomass DNA, 18 μL Platinum™ SYBR™ Green qPCR SuperMix (ThermoFisher), and 0.2 μM of each primer. Primers targeted the viral MCP gene (see primer sequences in cite) and the single-copy host gene CBLP: CBLP_qPCR_Fw (5′–GGA TGT GCA CGC GGT TTA CCC A–3′) and CBLP_qPCR_Rv (5′–GGT GTT GAC GTA GCC GTG GTG G–3′). Amplification was performed on a CFX96 real-time PCR system (Bio-Rad) with the following cycling conditions: 50 °C for 2 min; 95 °C for 2 min; 45 cycles of 95 °C for 15 s followed by 60 °C for 30 s; followed by a melting curve from 65° to 95°C. Relative viral-to-host copy number was calculated using the following equation, considering the primer efficiency (E) for each primer.

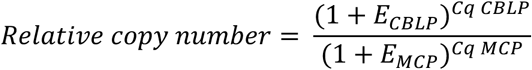

Total RNA from all GEVE strains was extracted from day-3 and day-5 timepoints using TRIzol, following Molnár et al. protocol with minor modifications^82^. Briefly, 1 mL TRIzol and 200 µL chloroform were used per sample, and an additional chloroform extraction step was incorporated after the initial TRIzol phase separation. RNA was precipitated with 1 volume of isopropanol, 1/10 volume of 3 M sodium acetate, and 1 µL glycogen (20 mg mL-1), incubated at -20 °C for 1 h, and recovered by centrifugation at 20,000 g for 30 min at 4 °C. Pellets were rinsed with 1 mL freshly prepared 80% ethanol and resuspended in 50 µL RNase-free water. Directional mRNA libraries were prepared by Novogene (Novogene, Beijing, China) using a poly(A)-enrichment workflow. Sequencing was performed on the NovaSeq X Plus platform (PE150).

At ten days post-inoculation, GEVE+ cultures were filtered, and the supernatant was concentrated using tangential flow filtration. Viral particles were pelleted at 20,000g for 15 min at 4 °C, resuspended in molecular-grade water, filtered through a 0.8 μm PES syringe filter, stained and analyzed by flow cytometry.

### RNA-Seq data analysis

Reads were trimmed using TrimGalore v0.6.11^83^, and rRNA-derived reads were removed using SortMeRNA v4.4.0^84^. Paired-end reads were aligned to the gapless CC-2937 assembly using STAR (version 2.7.11b)^58^ with the parameters `--outFilterMultimapNmax 100 --winAnchorMultimapNmax 100`. Gene and TE subfamily expression were quantified using TEcount (version 2.2.4) from the TEToolkit suite^85^. TElocal (version 1.1.3), also from the TEToolkit suite, was used to quantify the expression of individual TE loci. Differential expression testing was performed using DESeq2 (version 1.49.4)^86^. Genes and TEs were defined as differentially expressed if they had a Benjamini-Hochberg adjusted *p*-value < 0.05 and an absolute log2 fold change > 1. Variance-stabilized expression values calculated using the DESeq2 `vst` function were used for principal component analysis (PCA), while counts normalized using the median-of-ratios method^87^ implemented by DESeq2 were used for all other analyses requiring normalized RNA-seq counts.

### Quantification and Statistical Analysis

#### Genomic feature annotation and co-occurrence analysis

Six genomic features were annotated across all 506 canonical GEVE genes: BVP-cluster and DVP-cluster membership (defined previously^16^); presence of N6-methyladenine (6mA) DNA modification (identified from ONT); presence of spliceosomal introns (annotated during gene prediction; 61 genes, excluding the self-splicing intron of the MCP gene); possession of an NCLDV-type late promoter motif; and identity as a virion structural protein gene (determined by prior LC-MS/MS proteomics of isolated GEVE virions). See section “Punuivirus GEVE gene model reannotation” for details.

Pairwise co-occurrence among all six features was assessed by two-sided Fisher’s exact tests across all 14 non-exclusive feature pairs (the BVP/DVP pair was excluded as mutually exclusive by construction). Raw p-values were corrected for multiple testing using the Benjamini-Hochberg false discovery rate (BH-FDR) procedure [SciPy 1.13.0] across all 14 tests. Odds ratios (OR) and BH-adjusted q-values are reported in **Supplementary Table 5**.

#### Single-cell data processing and GEVE expression states

Sequencing reads were processed with split-pipe (Parse Biosciences, version 1.1.2) against a custom *Chlamydomonas reinhardtii* CC-2937 and GEVE reference, annotated with a GTF file that defined gene, exon, and intron coordinates. The two sublibraries were merged into a single gene expression matrix. Unless otherwise noted, analyses used library-size-normalized, log1p-transformed expression values [Scanpy 1.10.1]^88^. GEVE expression states were defined from the distribution of per-cell GEVE UMI counts. Cells in the upper tail of a log-normal fit to this distribution (≥ 43 GEVE UMIs; n = 39) were classified as High-GEVE, and cells with 10–42 GEVE UMIs (n = 1,696, including 9 virocells) as Active GEVE Expressing Cells. Median sequencing depth differed across culture stages, from more than 1,000 UMIs per cell at DPI 4 to fewer than 500 at DPI 6–7 (see **Appendix**). The 85 GEVE genes detected at higher rates in High-GEVE cells than in Active GEVE Expressing Cells were grouped by hierarchical clustering into a lytic module (51 genes) and a latent module (34 genes). The transcriptional program of each cell was summarized as the log₂ ratio of mean lytic-module to latent-module expression. Full details of reference construction, read processing, normalization, threshold selection, and module identification are provided in the **Appendix** [SciPy 1.13.0, Seaborn 0.13.2, Matplotlib 3.8.4]^89,90^.

#### Cell-state classification: virocells and latent cells

For each GEVE-expressing cell, the lytic/latent module ratio (computed on raw UMI counts; see **Appendix** Methods) and the GEVE/host UMI ratio, defined as (GEVE UMIs + 0.5) / (*C. reinhardtii* UMIs + 0.5), were used to classify cells. Virocells were defined as cells satisfying all three criteria simultaneously: total GEVE UMIs ≥ 10, log₂ lytic/latent module ratio > 0, and GEVE/host UMI ratio > 0.02; 21 cells met this definition (1, 3, 12 and 5 cells at DPI 4, 5, 6 and 7, respectively). All other cells with at least 10 GEVE UMIs were classified as latent GEVE cells; together they form the group all latent GEVE cells (≥ 10) (n = 1,714; **Appendix**). For comparisons with virocells at similar viral load, two subsets of latent GEVE cells were used: High-GEVE latent cells (GEVE ≥ 43 UMIs; n = 27; **Appendix**, Supplementary Table 8) and the 21 High-GEVE latent cells with the highest total GEVE UMI counts (top-21 High-GEVE latent cells; Figure S3, **Appendix**). All High-GEVE latent cells came from DPI 4, whereas 20 of 21 virocells came from DPI 5–7. Because median sequencing depth was several-fold higher at DPI 4 (see **Appendix**), detection-rate differences between virocells and High-GEVE latent cells may partly reflect culture stage and sequencing depth [pandas 2.2.2, NumPy 1.26.4].

#### Transcriptional displacement analysis

Host UMI fraction (*C. reinhardtii* UMIs / total UMIs per cell) was compared between High-GEVE and virocell populations by two-sided Mann-Whitney U test, with effect size as rank-biserial correlation [SciPy 1.13.0, NumPy 1.26.4]. Two additional depth-corrected checks were run to rule out a sequencing-depth confound: a partial Spearman correlation between GEVE and host UMIs controlling for total UMI count (permutation-tested)[SciPy 1.13.0, NumPy 1.26.4], and a regression-residual test [statsmodels 0.14.1] comparing observed to depth-predicted host UMI counts, fit on the depth-matched GEVE-background and GEVE-active populations. All three tests agreed in direction: host UMI fraction was lower in virocells (median 80.0%) than in High-GEVE cells (median 89.2%; rank-biserial r = −0.78, p = 4.8 × 10⁻⁶); the partial correlation between GEVE and host UMIs was significantly negative in virocells (r = −0.736, permutation p = 5.0 × 10⁻⁴) but not in High-GEVE cells (r = 0.097, p = 0.57); and virocells fell significantly below depth-predicted host counts (p = 2.9 × 10⁻⁶), significantly more than High-GEVE cells did (between-group p = 1.2 × 10⁻⁶).

#### Statistical Analysis of 5mC patterns

Statistical analyses were performed using R (version 4.5.1). The enrichment of TE subfamilies among the hypermethylated and hypomethylated windows outside the GEVE in the GEVE-samples was tested using a genomic-region permutation test. The randomizeRegions function from the regioneR R/Bioconductor package^91^ (version 1.41.3) was used to randomly redistribute the windows across the genome, excluding the GEVE, and the number of overlaps with each TE subfamily was counted. This process was repeated 10,000 times for the hypermethylated and hypomethylated windows separately. For each TE subfamily, the number of randomizations where the number of overlaps was equal to or greater than the actual observed number of overlaps was used to calculate a permutation-based p-value. The Benjamini-Hochberg procedure was then used for multiple-testing correction.

The over-representation of KEGG pathways containing at least 10 genes among differentially expressed genes was tested using a one-sided Fisher’s exact test, with all genes tested for differential expression by DESeq2 and associated with at least one KEGG pathway used as the background gene set. The Benjamini-Hochberg procedure was then used for multiple-testing correction.

#### Phylogenetic analysis of TET dioxygenases

TET dioxygenases from the *C. reinhardtii* reference genome^43^ and CC-2937 (one from punuivirus, four from Helitron transposons) were supplemented with transposon-encoded fungal TET proteins identified by Iyer et al.^45^ The fungal TETs were clustered using CD-HIT v4.8.1^92^ so that proteins with at least 60% sequence similarity spanning at least 60% of the smaller protein in the comparison were reduced to a single representative (options -d 0 -aS 0.6 -c 0.6 -G 0 -g 1 -n 3). Alignment was performed with MAFFT v7.526^93^ using the auto flag. Sequences missing the core TET domain were manually removed, and alignment columns with more than 30% gaps were subsequently filtered out using trimAl v1.5.rev1^94^ with the flag -gt 0.7. A maximum-likelihood phylogeny was produced using IQ-TREE v3.1.2^95^ using ModelFinder^96^ and 1000 replicates of ultrafast bootstrapping (-m MFP -bb 1000).

## Acknowledgments

We acknowledge use of the Virginia Tech Advances Research Computing center and Queen Mary University of London Apocrita HPC facilities for the computational work performed in this study. We are grateful to Assaf Vardi and Talia Sima Shaler for comments on an earlier version of the single-cell RNA-seq analysis. We thank members of the Aylward lab for comments on an earlier version of the manuscript. US, MPEG, and FOA were supported by National Institutes of Health R35 grant no. 1R35GM147290. AdM and RH were supported by a ERC-Starting Grant (950230).

## Author Contributions

Conceptualization: F.O.A. and A.d.M.; formal analysis: U.S., R.H., M.P.E.G., and R.J.C.; funding acquisition: F.O.A. and A.d.M.; investigation: U.S., L.X., M.P.E.G., and R.J.C.; methodology: U.S., R.H., M.P.E.G., and R.J.C.; visualization: U.S., R.H., M.P.E.G., A.d.M. and R.J.C.; supervision: F.O.A. and A.d.M.; writing – original draft: F.O.A., U.S. and A.d.M.; and writing – review & editing: R.H., M.P.E.G., R.J.C., F.O.A., U.S. and A.d.M.

## Declaration of Interests

The authors declare no competing interests.

## Extended Data Figures

**Figure S1.**
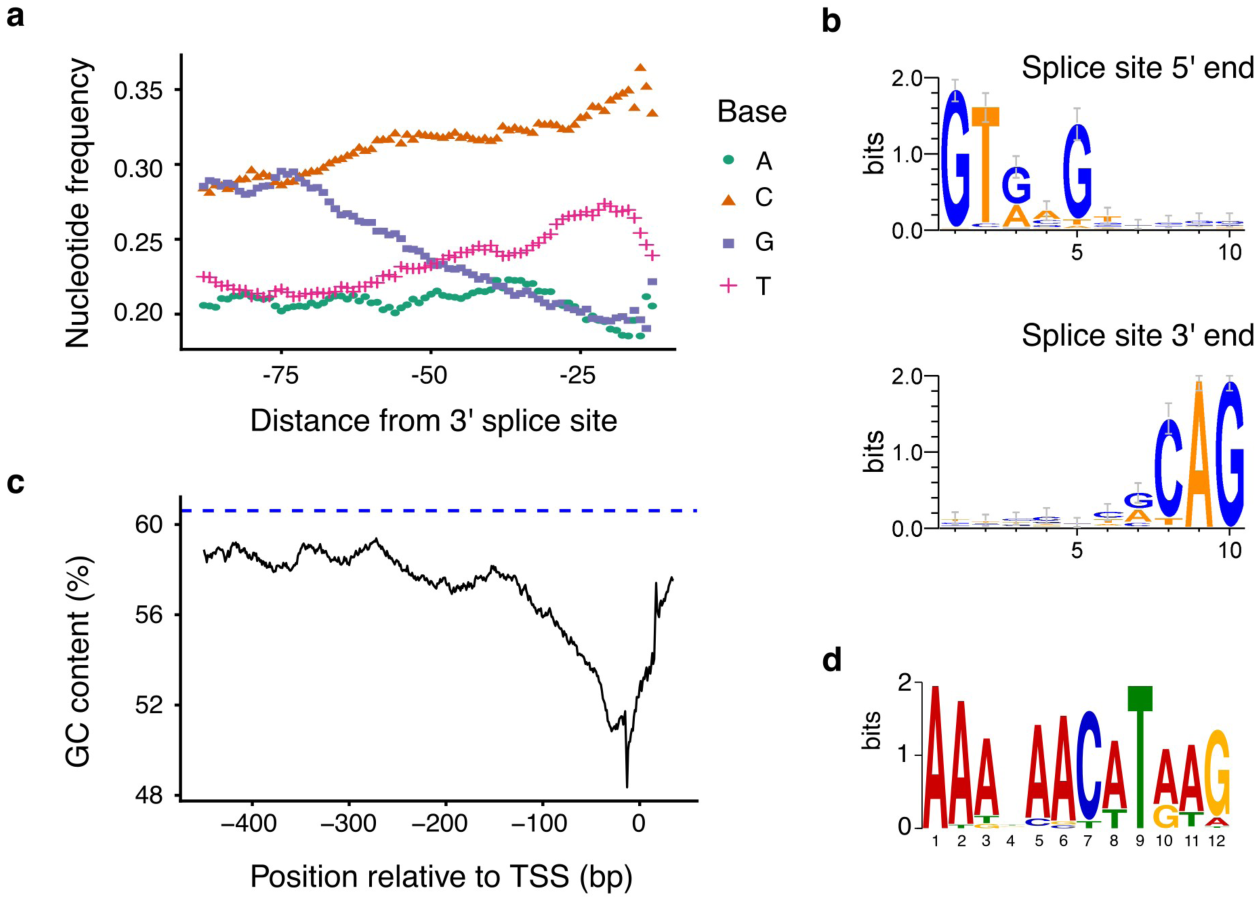
Sequence composition of GEVE introns and upstream gene regions. (A) Nucleotide composition calculated using a 25-bp sliding window across the aligned 3′ ends of Punuvirus introns. (B) Sequence logos showing canonical splice-site motifs at the aligned 5′ and 3′ ends of punuivirus introns. (C) GC content calculated using a 30-bp sliding window across regions upstream of the transcription start sites (TSS) of punuivirus genes. The dashed line indicates the average GC content of the GEVE region. (D) Enriched sequence motif identified in the upstream regions of GEVE genes.

**Figure S2.**
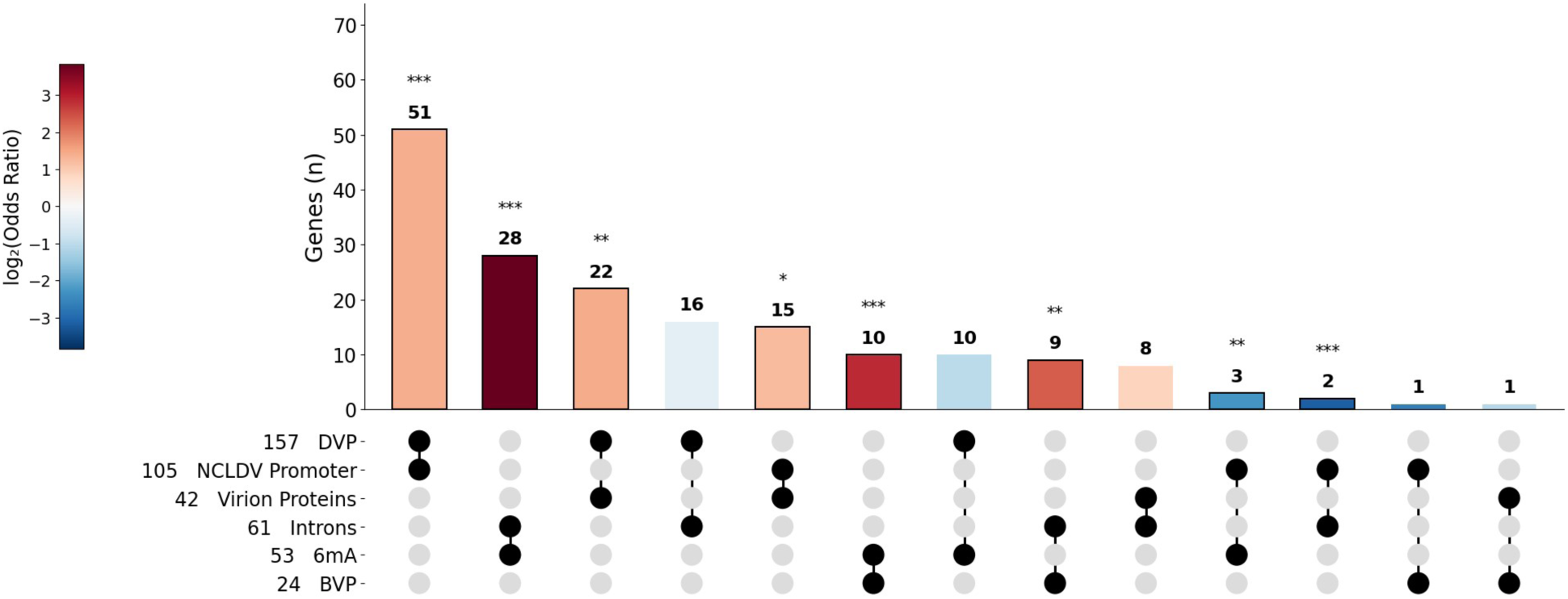
Pairwise co-occurrence among six binary GEVE genomic features (BVP, DVP, 6mA, Introns, NCLDV Promoter, Virion Proteins) across 506 annotated GEVE genes. Each feature’s total set size is printed to the left of its name. Bars show the number of genes carrying both features of each pair (inclusive count n(A & B), printed above each bar; genes can carry additional features, so bars do not sum to 506), sorted by descending count. Bar color encodes log₂(odds ratio) from a two-sided Fisher’s exact test for co-occurrence of the two features (**Supplementary Table 5**); asterisks mark BH-FDR-adjusted significance (* q < 0.05, ** q < 0.01, *** q < 0.001), and bars outlined in black also reach q < 0.05. The BVP & DVP pair is absent (mutually exclusive by definition), and the 6mA & Virion Proteins pair is absent because no gene carries both features. Introns denote spliceosomal introns (61 genes).

**Figure S3.**
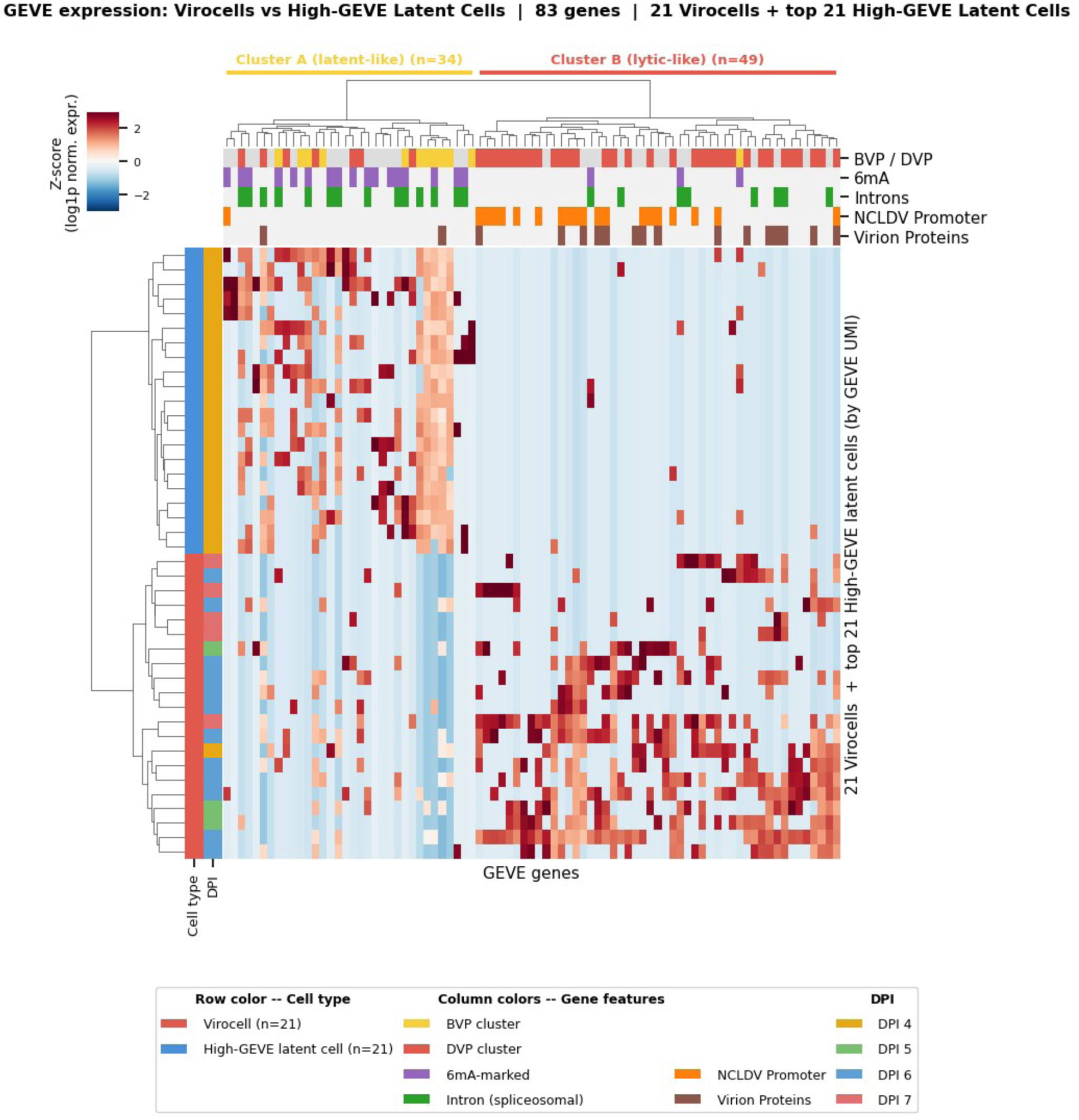
GEVE expression heatmap comparing virocells and top-21 High-GEVE latent cells. Virocells (n = 21) were defined by three simultaneous criteria: GEVE ≥ 10 UMIs, log₂ lytic/latent module ratio > 0, and GEVE-to-*C. reinhardtii* UMI ratio > 0.02. The comparison group comprises the 21 High-GEVE latent cells (GEVE ≥ 43 UMIs, not virocells) with the highest total GEVE UMI counts; all are from DPI 4. Restricted to 83 genes detected in > 20% of virocells or > 10% of top-21 High-GEVE latent cells. Z-scored log1p-normalized expression (10k UMIs), clipped at ±3. Hierarchical clustering (Ward / Euclidean). Column color bars indicate BVP/DVP cluster, 6mA, NCLDV promoter, spliceosomal intron, and virion protein annotation. Two gene clusters emerge: Cluster A (latent-like, n = 34 genes, BVP/6mA-enriched) and Cluster B (lytic-like, n = 49 genes, DVP/NCLDV promoter/virion-enriched).

**Figure S4.**
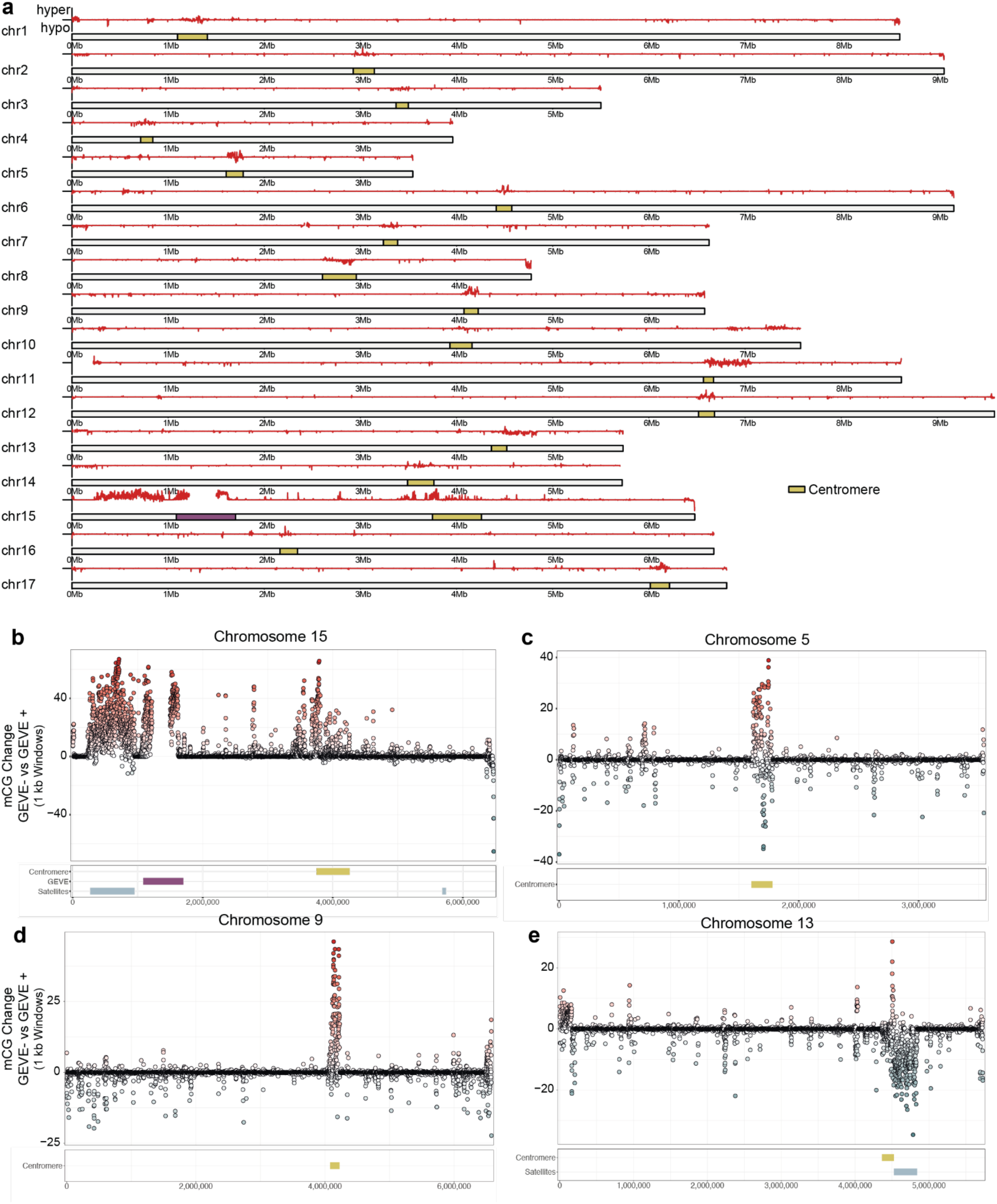
Genome-wide redistribution of CG methylation following GEVE deletion. (A) Karyotype showing the average difference in CG methylation between GEVE− and GEVE+ strains across 10-kb genomic windows. Red indicates increased methylation and blue decreased methylation in GEVE− strains. Chromosome positions of centromeres and the chromosome 15 GEVE locus are indicated. (B–E) Detailed views of chromosomes 15 (B), 5 (C), 9 (D), and 13 (E), illustrating representative examples of regional gains and losses of CG methylation. Differentially methylated regions occur as discrete chromosomal domains rather than as uniform genome-wide changes, indicating extensive epigenomic redistribution following GEVE deletion.

**Figure S5.**
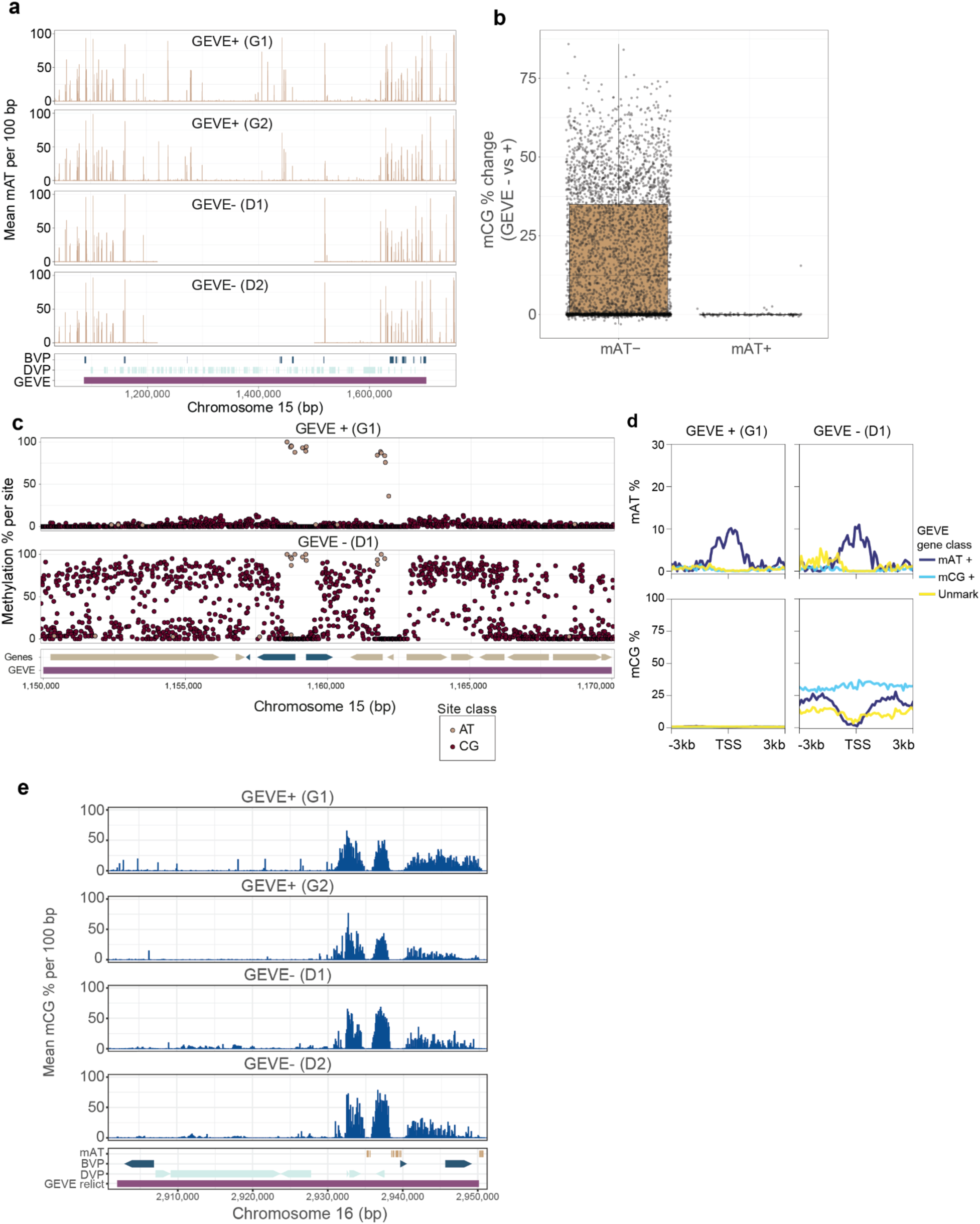
Adenine methylation is associated with protection from CG methylation within the GEVE. (A) 6mA (mAT) methylation on GEVE in both strain types remains largely unchanged beyond the deletion. (B) Distribution of changes in 5mC across the GEVE, calculated in 100-bp windows and stratified according to the presence or absence of 6mA marks. Windows containing mAT show no 5mC gains following GEVE deletion, unlike previously unmarked windows. (C) Representative examples of 6mA-marked BVP genes illustrating the relationship between the two DNA modifications. CpG methylation accumulates following GEVE deletion but remains depleted around promoter-associated 6mA peaks. (D) Average DNA methylation profiles centered on transcription start sites (TSSs) of mAT-marked genes, mCG-marked genes, and unmodified genes. Promoter-associated 6mA regions remain protected from CpG methylation, despite the overall increase in mCG across the GEVE. (E) 5mC landscape of the chromosome 16 GEVE relic. Although overall 5mC remains unchanged, regions carrying promoter-associated 6mA continue to exhibit local depletion of 5mC, indicating that the antagonistic relationship between the two DNA modifications is maintained in degenerated viral loci.

**Figure S6.**
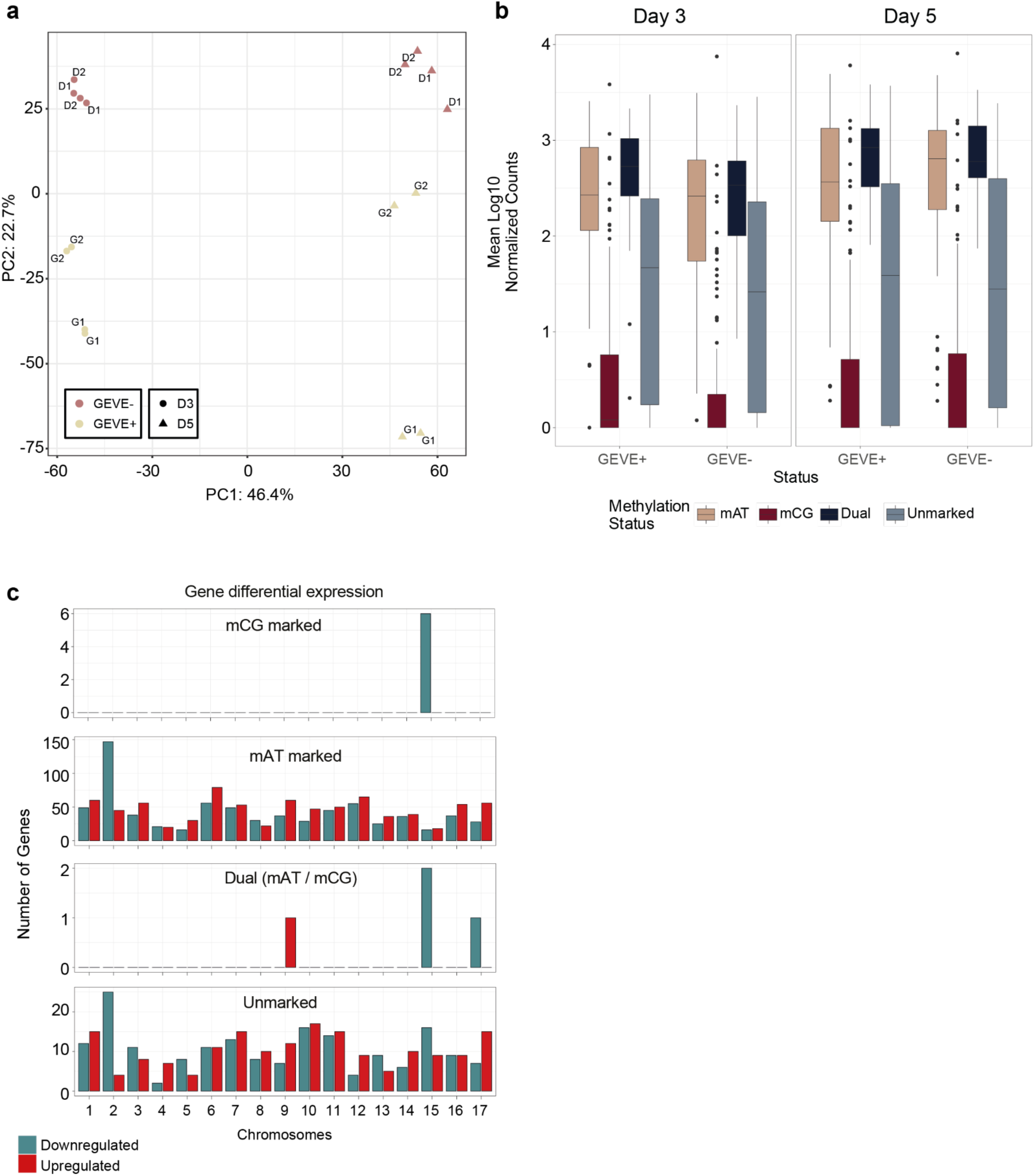
Transcriptional changes following GEVE deletion extend beyond loci that undergo CpG methylation changes. (A) Clustering of RNA-seq samples of each genotype and culture day post inoculation, showing most variation is derived from days in culture (PC1) and then genotype (PC2). GEVE-strains: D1 and D2. GEVE+ strains: G1 and G2. (B) Distribution of average expression levels in Day 3 and Day 5 of GEVE genes stratified by methylation status, showing that genes that gain 5mC are seldom expressed in any condition, whereas 6mA marked genes remain highly expressed even after GEVE deletion. (C) Number of differentially expressed host protein-coding genes between GEVE− and GEVE+ strains, stratified by chromosome, 5mC status, and direction of expression change. Although chromosome 15 is enriched for downregulated mCG-marked genes, other chromosomes (particularly chromosome 2) display substantial transcriptional changes despite limited alterations in 5mC, indicating that many expression changes are indirect.

**Figure S7.**
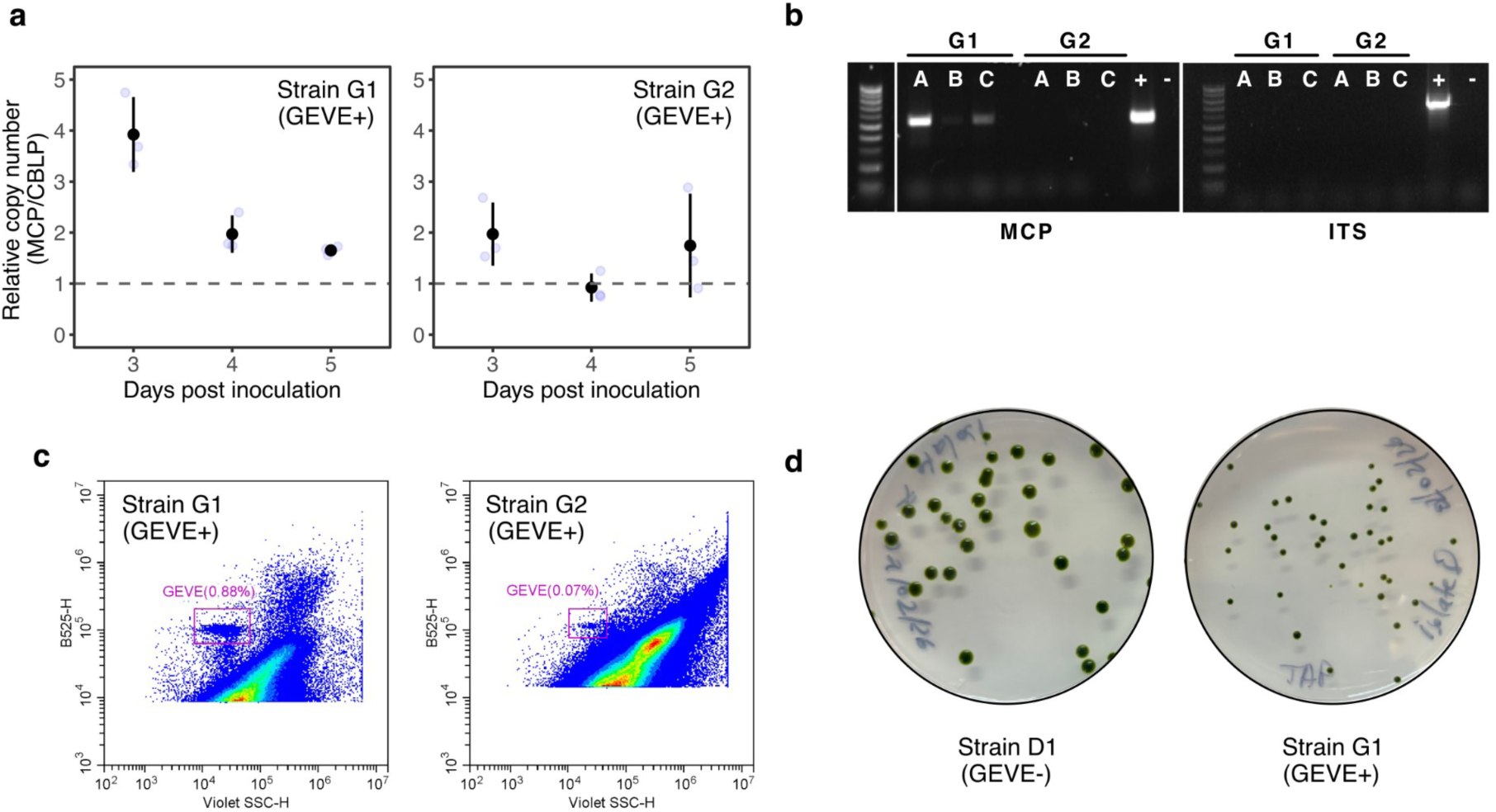
Assessment of viral induction and phenotypic differences among GEVE strains. (A) Viral genome replication quantified by qPCR as the relative copy number of the viral major capsid protein (MCP) gene normalized to the single-copy host gene CBLP. (B) PCR detection of extracellular virions in DNase-treated supernatants collected 10 days post-inoculation. For each GEVE + strain (G1 and G2), three biological replicates (A–C) were assayed by amplification of the MCP gene, with host ITS serving as a negative control for contaminating host DNA. Lane order: 100-bp ladder; GEVE + G1 replicates A– C; GEVE + G2 replicates A–C; positive control; non-template control. Left: MCP amplification. Right: ITS amplification. (C) Flow cytometry detection of virions concentrated from culture supernatants of both GEVE + strains. Plots display green fluorescence (B525) versus Violet Side Scatter (Violet SSC). (D) Representative colony size from exponential-phase cultures of GEVE - D1 and GEVE + G1 strains after 11 days of growth.

**Figure S8.**
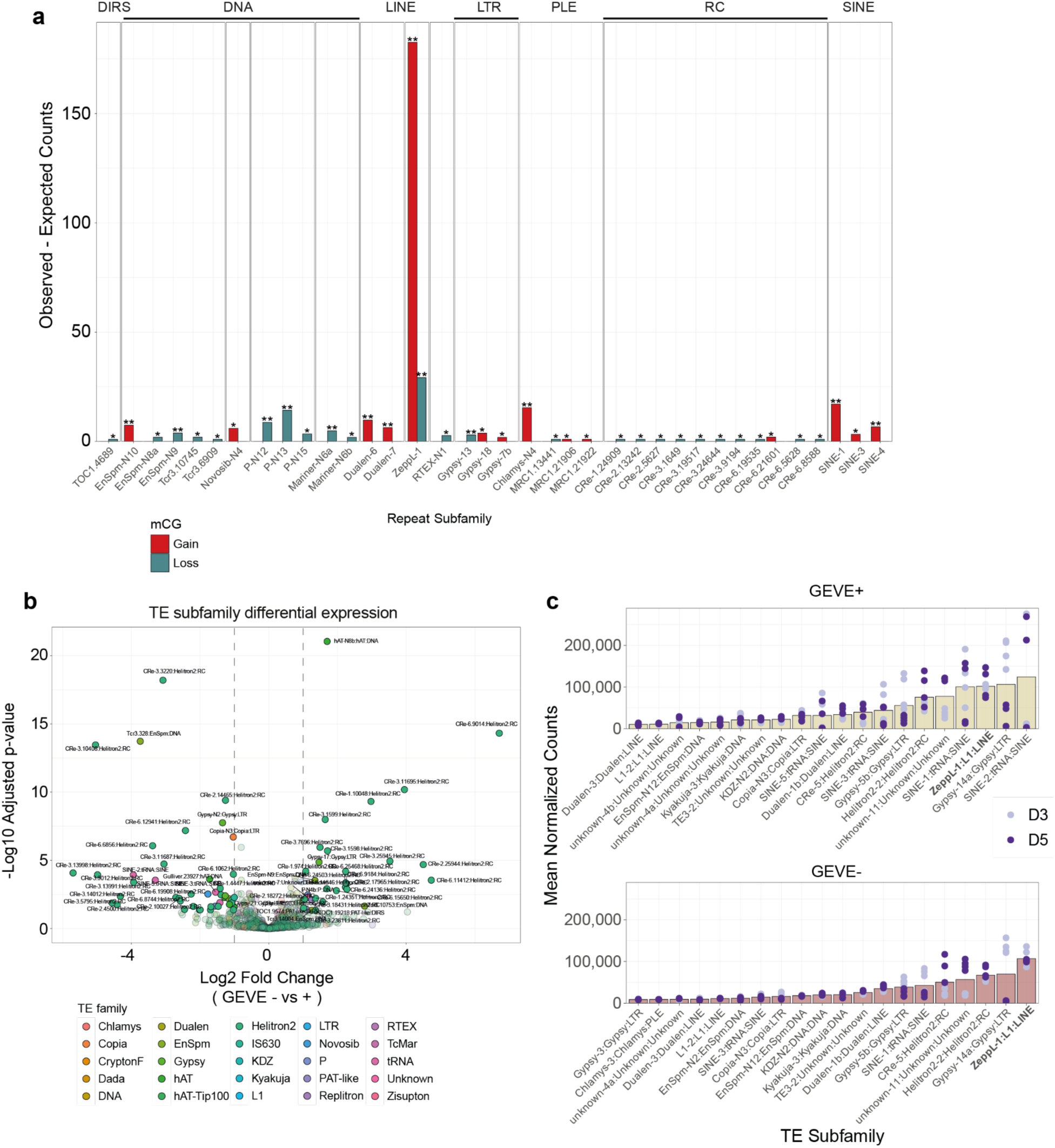
Transposable Element subfamily epigenetic and transcriptional changes. (A) Enrichment of transposable element (TE) subfamilies within genomic windows that gain (red) or lose (blue) 5mC in GEVE− relative to GEVE+ strains. The most pronounced gain in 5mC methylation occurs over ZeppL retrotransposons, consistent with the extensive methylation changes observed at centromeres, whereas only a limited number of TE subfamilies are associated with methylation loss. Asterisks indicate significant enrichment after multiple testing correction using the Benjamini-Hochberg procedure (* < 0.05; ** < 0.01; permutation test: details in Methods). (B) Differential expression of subfamily level TEs between GEVE− and GEVE+ strains. Each point represents a single subfamily colored by TE family. Dashed lines indicate differential expression thresholds (log2 fold-change ±1). (C) Normalized expression counts of the most transcribed TE subfamilies at days 3 and 5 of culture growth (D3 and D5) in GEVE+ and GEVE– strains. ZeppL retrotransposons are among the top 3 most expressed TEs in each condition.

## Supplementary Tables

**Supplementary Table 1. Complete annotation of the 506 canonical GEVE genes**. Each row is one annotated GEVE gene. Columns: genomic coordinates (contig, start, end, strand), BVP/DVP cluster assignment, binary feature flags (6mA modification, spliceosomal intron, NCLDV-type late promoter [FIMO q < 0.05], virion structural protein [LC-MS/MS]), number of introns per gene, virion protein accession, and functional annotations (NCLDV marker, Pfam domain, GVOG family, EggNOG description, HHpred top hit).

**Supplementary Table 2. GEVE genes bearing spliceosomal introns.** The 61 GEVE genes containing spliceosomal introns. Columns: BVP/DVP/Unclassified cluster assignment, genomic coordinates, number of introns per gene, 6mA modification flag, NCLDV-type late promoter motif (FIMO q < 0.05), virion structural protein flag, and functional annotations. Of 61 intron-bearing genes, 28 also carry 6mA modification and 9 are in the BVP cluster, consistent with co-enrichment of introns with the latent gene program.

**Supplementary Table 3. GEVE genes bearing N6-methyladenine (6mA) modification.** The 53 GEVE genes with upstream N6-methyladenine (6mA) modification, called from Oxford Nanopore Technologies (ONT) reads. Columns: BVP/DVP/Unclassified cluster assignment, genomic coordinates, spliceosomal intron flag, NCLDV-type late promoter motif (FIMO q < 0.05), virion structural protein flag, and functional annotations.

**Supplementary Table 4. BVP and DVP cluster gene list.** All 181 GEVE genes assigned to the Before Viral Production (BVP; n = 24) or During Viral Production (DVP; n = 157) transcriptional cluster (ErazoGarcia 2025). Binary feature flags: 6mA modification, spliceosomal intron, NCLDV-type late promoter motif (FIMO q < 0.05), virion structural protein identity. Annotation: first available functional annotation (NCLDV marker, GVOG family, Pfam domain, or EggNOG description).

**Supplementary Table 5. Pairwise Fisher’s exact test statistics for co-occurrence of six binary GEVE genomic features.** All non-exclusive combinations of BVP, DVP, 6mA, Introns, NCLDV Promoter, and Virion Proteins across 506 annotated GEVE genes (14 tested pairs; BVP vs. DVP excluded as mutually exclusive). Columns: co-occurrence count n(A & B), marginal counts, Odds Ratio, raw two-sided p-value, and BH-FDR-corrected q-value. Nine of 14 pairs reach significance at q < 0.05: six positive associations (6mA–Introns, DVP–NCLDV Promoter, BVP–6mA, BVP–Introns, DVP–Virion Proteins, NCLDV Promoter–Virion Proteins) and three negative associations (Introns–NCLDV Promoter, 6mA–NCLDV Promoter, 6mA–Virion Proteins).

**Supplementary Table 6. GEVE genes bearing an NCLDV-type late promoter motif, identified by FIMO.** The 105 GEVE genes with a significant FIMO hit (q < 0.05, BH-FDR) for the STREME-1 motif (consensus AAANAACATAAG; 12 bp) in the 200 bp upstream sequence of each gene. Columns: Gene ID, BVP/DVP/Unclassified cluster, FIMO score, p-value, q-value, matched sequence, virion structural protein flag, and first available functional annotation.

**Supplementary Table 7. GEVE virion structural protein genes identified by LC-MS/MS proteomics.** The 42 GEVE genes encoding proteins detected in isolated GEVE virions by LC-MS/MS^16^. Columns: Gene ID, strand, BVP/DVP/Unclassified cluster, NCLDV-type late promoter flag, 6mA modification flag, spliceosomal intron flag, virion protein accession, and functional annotation.

**Supplementary Table 8. GEVE genomic feature enrichment: virocells versus High-GEVE latent cells**. Detection-rate enrichment of eight GEVE genomic feature categories in 21 virocells versus 27 High-GEVE latent cells (GEVE ≥ 43 UMIs, not meeting the virocell criteria; all from DPI 4), tested across all 506 canonical GEVE genes using Fisher’s exact test with Benjamini-Hochberg FDR correction across all 8 tests. Intron-bearing genes are split by BVP-cluster, DVP-cluster, or unclassified membership. Columns: Category, n genes, detection % in High-GEVE latent cells, detection % in virocells, odds ratio (virocell / High-GEVE latent detection), FDR, significance. OR > 1 = enriched in virocells (lytic program); OR < 1 = enriched in High-GEVE latent cells (latent program).

## Notes

### Competing Interest Statement

The authors have declared no competing interest.

## References

1. Lieberman, P.M. (2016). Epigenetics and Genetics of Viral Latency. Cell Host Microbe 19, 619– 628.

2. Sarre, L.A., Kim, I.V., Ovchinnikov, V., Olivetta, M., Suga, H., Dudin, O., Sebé-Pedrós, A., and de Mendoza, A. (2024). DNA methylation enables recurrent endogenization of giant viruses in an animal relative. Sci Adv 10, eado6406.

3. Schultz, S., Minch, B., Mimick, E., and Moniruzzaman, M. (2026). Extensive array of endogenous giant viral elements in a polar alga shows dynamic transcriptional response to abiotic stress. Curr Biol 36, 1811–1824.e3.

4. Bosmon, T., Abergel, C., and Claverie, J.-M. (2025). 20 years of research on giant viruses. Npj Viruses 3, 9.

5. Duchêne, C., Wang, L., and Coelho, S.M. (2026). Revisiting giant virus-host dynamics in brown algae: old stories and new perspectives. EMBO J 45, 1822–1835.

6. Moniruzzaman, M., Weinheimer, A.R., Martinez-Gutierrez, C.A., and Aylward, F.O. (2020). Widespread endogenization of giant viruses shapes genomes of green algae. Nature 588, 141–145.

7. Jivaji, A.M., Karki, S., Franken, E., Erazo-Garcia, M.P., Barth, Z.K., and Aylward, F.O. (2025). Giant endogenous viral elements in the genome of the model protist reveal past interactions with giant viruses. J Virol 99, e0071325.

8. Denoeud, F., Godfroy, O., Cruaud, C., Heesch, S., Nehr, Z., Tadrent, N., Couloux, A., Brillet-Guéguen, L., Delage, L., Mckeown, D., et al. (2024). Evolutionary genomics of the emergence of brown algae as key components of coastal ecosystems. Cell 187, 6943–6965.e39.

9. Zhao, H., Zhang, R., Wu, J., Meng, L., Okazaki, Y., Hikida, H., and Ogata, H. (2023). A 1.5-Mb continuous endogenous viral region in the arbuscular mycorrhizal fungus. Virus Evol 9, vead064.

10. Whitner, S.K., Amend, A.S., and Culley, A.I. (2026). Marine fungi harbor, and are co-diversified with, remnants of giant endogenous viral elements. npj Viruses, 1–11.

11. Myers, J.M., Schulz, F., Rahimlou, S., Yadav, V., Amses, K.R., Simmons, D.R., Sun, S., Orozco-Quime, M., Heitman, J., Stajich, J.E., et al. (2026). *Mycodnaviridae* is a clade of giant viruses that persistently infect zoosporic fungi. PLoS Biol 24, e3003937.

12. Liu, P., Vigneau, J., Craig, R.J., Barrera-Redondo, J., Avdievich, E., Martinho, C., Borg, M., Haas, F.B., Liu, C., and Coelho, S.M. (2024). 3D chromatin maps of a brown alga reveal U/V sex chromosome spatial organization. Nat Commun 15, 9590.

13. Blais, C., Colp, M.J., Sarre, L.A., de Mendoza, A., and Archibald, J.M. (2025). Epigenetic silencing and genome dynamics determine the fate of giant virus endogenizations in *Acanthamoeba*. BMC Biol 23, 171.

14. Sarre, L.A., Gastellou Peralta, G.A., Romero Charria, P., Ovchinnikov, V., and de Mendoza, A. (2025). Repressive cytosine methylation is a marker of viral gene transfer across divergent eukaryotes. Mol. Biol. Evol. 42, msaf176.

15. Bourdareau, S., Tirichine, L., Lombard, B., Loew, D., Scornet, D., Wu, Y., Coelho, S.M., and Cock, J.M. (2021). Histone modifications during the life cycle of the brown alga *Ectocarpus*. Genome Biol. 22, 12.

16. Erazo-Garcia, M.P., Sheyn, U., Barth, Z.K., Craig, R.J., Wessman, P., Jivaji, A.M., Ray, W.K., Svensson-Coelho, M., Cornwallis, C.K., Rengefors, K., et al. (2025). Cryptic infection of a giant virus in a unicellular green alga. Science 388, eads6303.

17. Arthofer, P., Delafont, V., Willemsen, A., Panhölzl, F., and Horn, M. (2022). Defensive symbiosis against giant viruses in amoebae. Proc Natl Acad Sci U S A 119, e2205856119.

18. Fischer, M.G., and Hackl, T. (2016). Host genome integration and giant virus-induced reactivation of the virophage mavirus. Nature 540, 288–291.

19. La Scola, B., Desnues, C., Pagnier, I., Robert, C., Barrassi, L., Fournous, G., Merchat, M., Suzan-Monti, M., Forterre, P., Koonin, E., et al. (2008). The virophage as a unique parasite of the giant mimivirus. Nature 455, 100–104.

20. Labadorf, A., Link, A., Rogers, M.F., Thomas, J., Reddy, A.S., and Ben-Hur, A. (2010). Genome-wide analysis of alternative splicing in *Chlamydomonas reinhardtii*. BMC Genomics 11, 114.

21. Oliveira, G.P., Andrade, A.C.D.S.P., Rodrigues, R.A.L., Arantes, T.S., Boratto, P.V.M., Silva, L.K.D.S., Dornas, F.P., Trindade, G. de S., Drumond, B.P., La Scola, B., et al. (2017). Promoter Motifs in NCLDVs: An Evolutionary Perspective. Viruses 9. 10.3390/v9010016.

22. Suhre, K., Audic, S., and Claverie, J.-M. (2005). Mimivirus gene promoters exhibit an unprecedented conservation among all eukaryotes. Proc Natl Acad Sci U S A 102, 14689–14693.

23. Aylward, F.O., Moniruzzaman, M., Ha, A.D., and Koonin, E.V. (2021). A phylogenomic framework for charting the diversity and evolution of giant viruses. PLoS Biol 19, e3001430.

24. Fu, Y., Luo, G.-Z., Chen, K., Deng, X., Yu, M., Han, D., Hao, Z., Liu, J., Lu, X., Dore, L.C., et al. (2015). N6-methyldeoxyadenosine marks active transcription start sites in *Chlamydomonas*. Cell 161, 879–892.

25. Baier, T., Jacobebbinghaus, N., Einhaus, A., Lauersen, K.J., and Kruse, O. (2020). Introns mediate post-transcriptional enhancement of nuclear gene expression in the green microalga *Chlamydomonas reinhardtii*. PLoS Genet 16, e1008944.

26. Zeng, L., Skinner, S.O., Zong, C., Sippy, J., Feiss, M., and Golding, I. (2010). Decision making at a subcellular level determines the outcome of bacteriophage infection. Cell 141, 682–691.

27. Veening, J.-W., Smits, W.K., and Kuipers, O.P. (2008). Bistability, epigenetics, and bet-hedging in bacteria. Annu Rev Microbiol 62, 193–210.

28. Ku, C., Sheyn, U., Sebé-Pedrós, A., Ben-Dor, S., Schatz, D., Tanay, A., Rosenwasser, S., and Vardi, A. (2020). A single-cell view on alga-virus interactions reveals sequential transcriptional programs and infection states. Sci Adv 6, eaba4137.

29. Fromm, A., Hevroni, G., Vincent, F., Schatz, D., Martinez-Gutierrez, C.A., Aylward, F.O., and Vardi, A. (2024). Single-cell RNA-seq of the rare virosphere reveals the native hosts of giant viruses in the marine environment. Nat Microbiol 9, 1619–1629.

30. Hevroni, G., Vincent, F., Ku, C., Sheyn, U., and Vardi, A. (2023). Daily turnover of active giant virus infection during algal blooms revealed by single-cell transcriptomics. Sci Adv 9, eadf7971.

31. Moniruzzaman, M., Erazo Garcia, M.P., Farzad, R., Ha, A.D., Jivaji, A., Karki, S., Sheyn, U., Stanton, J., Minch, B., Stephens, D., et al. (2023). Virologs, viral mimicry, and virocell metabolism: the expanding scale of cellular functions encoded in the complex genomes of giant viruses. FEMS Microbiol Rev 47. 10.1093/femsre/fuad053.

32. Rowe, J.M., Jeanniard, A., Gurnon, J.R., Xia, Y., Dunigan, D.D., Van Etten, J.L., and Blanc, G. (2014). Global analysis of *Chlorella variabilis* NC64A mRNA profiles during the early phase of Paramecium bursaria chlorella virus-1 infection. PLoS One 9, e90988.

33. Bickels Nuri, R., Feldmesser, E., Fridmann-Sirkis, Y., Keren-Shaul, H., Nevo, R., Minsky, A., and Reich, Z. (2024). *Acanthamoeba polyphaga* de novo transcriptome and its dynamics during Mimivirus infection. Sci Rep 14, 25894.

34. Romero Charria, P., Navarrete, C., Ovchinnikov, V., Xu, L., Sarre, L.A., Shabardina, V., Ksiezopolska, E., Casacuberta, E., Lara-Astiaso, D., Sebé-Pedrós, A., et al. (2025). Adenine DNA methylation associated with transcriptionally permissive chromatin is widespread across eukaryotes. Nat Genet 57, 3126–3136.

35. Craig, R.J., Gallaher, S.D., Shu, S., Salomé, P.A., Jenkins, J.W., Blaby-Haas, C.E., Purvine, S.O., O’Donnell, S., Barry, K., Grimwood, J., et al. (2023). The *Chlamydomonas* Genome Project, version 6: Reference assemblies for mating-type plus and minus strains reveal extensive structural mutation in the laboratory. Plant Cell 35, 644–672.

36. Chaux-Jukic, F., O’Donnell, S., Craig, R.J., Eberhard, S., Vallon, O., and Xu, Z. (2021). Architecture and evolution of subtelomeres in the unicellular green alga *Chlamydomonas reinhardtii*. Nucleic Acids Res. 49, 7571–7587.

37. Lopez, D., Hamaji, T., Kropat, J., De Hoff, P., Morselli, M., Rubbi, L., Fitz-Gibbon, S., Gallaher, S.D., Merchant, S.S., Umen, J., et al. (2015). Dynamic Changes in the Transcriptome and Methylome of *Chlamydomonas reinhardtii* throughout Its Life Cycle. Plant Physiol. 169, 2730– 2743.

38. Craig, R.J., Hasan, A.R., Ness, R.W., and Keightley, P.D. (2021). Comparative genomics of *Chlamydomonas*. Plant Cell 33, 1016–1041.

39. Liu, D., Wang, M., Gent, J.I., Sun, P., Dawe, R.K., and Umen, J. (2025). Two CENH3 paralogs in the green alga *Chlamydomonas reinhardtii* have a redundantly essential function and associate with ZeppL-LINE1 elements. Plant J. 122, e70153.

40. Ngan, C.Y., Wong, C.-H., Choi, C., Yoshinaga, Y., Louie, K., Jia, J., Chen, C., Bowen, B., Cheng, H., Leonelli, L., et al. (2015). Lineage-specific chromatin signatures reveal a regulator of lipid metabolism in microalgae. Nat Plants 1, 15107.

41. Ge, T., Gui, X., Xu, J.-X., Xia, W., Wang, C.-H., Yang, W., Huang, K., Walsh, C., Umen, J.G., Walter, J., et al. (2024). DNA cytosine methylation suppresses meiotic recombination at the sex-determining region. Sci. Adv. 10, eadr2345.

42. Ross, S.E., and Bogdanovic, O. (2019). TET enzymes, DNA demethylation and pluripotency. Biochem. Soc. Trans. 47, 875–885.

43. Xue, J.-H., Chen, G.-D., Hao, F., Chen, H., Fang, Z., Chen, F.-F., Pang, B., Yang, Q.-L., Wei, X., Fan, Q.-Q., et al. (2019). A vitamin-C-derived DNA modification catalysed by an algal TET homologue. Nature 569, 581–585.

44. Iyer, L.M., Zhang, D., Burroughs, A.M., and Aravind, L. (2013). Computational identification of novel biochemical systems involved in oxidation, glycosylation and other complex modifications of bases in DNA. Nucleic Acids Res. 41, 7635–7655.

45. Iyer, L.M., Zhang, D., de Souza, R.F., Pukkila, P.J., Rao, A., and Aravind, L. (2014). Lineage-specific expansions of TET/JBP genes and a new class of DNA transposons shape fungal genomic and epigenetic landscapes. Proc. Natl. Acad. Sci. U. S. A. 111, 1676–1683.

46. Kojima, K.K. (2025). Heterogeneous lineages of DNA transposons encode a TET/JBP dioxygenase in fungi. Biology (Basel) 14, 1741.

47. Li, W., Zhang, T., Sun, M., Shi, Y., Zhang, X.-J., Xu, G.-L., and Ding, J. (2021). Molecular mechanism for vitamin C-derived C-glyceryl-methylcytosine DNA modification catalyzed by algal TET homologue CMD1. Nat Commun 12, 744.

48. Medvedeva, S., Guyet, U., Pelletier, E., Ruscheweyh, H.-J., Sunagawa, S., Ogata, H., Aylward, F.O., Gaïa, M., Yutin, N., Koonin, E.V., et al. (2026). Widespread and intron-rich mirusviruses are predicted to reproduce in nuclei of unicellular eukaryotes. Nat Microbiol 11, 228–239.

49. Agarkova, I.V., Dunigan, D.D., and Van Etten, J.L. (2006). Virion-associated restriction endonucleases of chloroviruses. J Virol 80, 8114–8123.

50. Jeudy, S., Rigou, S., Alempic, J.-M., Claverie, J.-M., Abergel, C., and Legendre, M. (2020). The DNA methylation landscape of giant viruses. Nat Commun 11, 2657.

51. Dodds, J.A. (1979). Viruses of marine algae. Experientia 35, 440–442.

52. Cohen, J.I. (2020). Herpesvirus latency. J Clin Invest 130, 3361–3369.

53. Shnayder, M., Nachshon, A., Krishna, B., Poole, E., Boshkov, A., Binyamin, A., Maza, I., Sinclair, J., Schwartz, M., and Stern-Ginossar, N. (2018). Defining the Transcriptional Landscape during Cytomegalovirus Latency with Single-Cell RNA Sequencing. mBio 9. 10.1128/mBio.00013-18.

54. Murphy, P.J., and Berger, F. (2023). The chromatin source-sink hypothesis: a shared mode of chromatin-mediated regulations. Development 150, dev201989.

55. Bellas, C., and Sommaruga, R. (2026). A framework for Polinton-like virus diversity reveals links to multiple viral classes and *Nucleocytoviricota*. bioRxiv. 10.64898/2026.06.19.733378.

56. Sarkans, U., Gostev, M., Athar, A., Behrangi, E., Melnichuk, O., Ali, A., Minguet, J., Rada, J.C., Snow, C., Tikhonov, A., et al. (2018). The BioStudies database-one stop shop for all data supporting a life sciences study. Nucleic Acids Res. 46, D1266–D1270.

57. Cuypers, W.L., Ceylan, H., Turcksin, E., Raes, L., de Vrij, N., Michiels, J., Coppens, S., de Block, T., Jansen, D., Ariën, K.K., et al. (2026). SquiDBase: a community resource of raw nanopore data from microbes. NAR Genom. Bioinform. 8, lqaf213.

58. Dobin, A., Davis, C.A., Schlesinger, F., Drenkow, J., Zaleski, C., Jha, S., Batut, P., Chaisson, M., and Gingeras, T.R. (2013). STAR: ultrafast universal RNA-seq aligner. Bioinformatics 29, 15–21.

59. Li, H., Handsaker, B., Wysoker, A., Fennell, T., Ruan, J., Homer, N., Marth, G., Abecasis, G., Durbin, R., and 1000 Genome Project Data Processing Subgroup (2009). The Sequence Alignment/Map format and SAMtools. Bioinformatics 25, 2078–2079.

60. Gabriel, L., Brůna, T., Hoff, K.J., Ebel, M., Lomsadze, A., Borodovsky, M., and Stanke, M. (2024). BRAKER3: Fully automated genome annotation using RNA-seq and protein evidence with GeneMark-ETP, AUGUSTUS, and TSEBRA. Genome Res. 34, 769–777.

61. Nawrocki, E.P., and Eddy, S.R. (2013). Infernal 1.1: 100-fold faster RNA homology searches. Bioinformatics 29, 2933–2935.

62. Kalvari, I., Nawrocki, E.P., Argasinska, J., Quinones-Olvera, N., Finn, R.D., Bateman, A., and Petrov, A.I. (2018). Non-Coding RNA Analysis Using the Rfam Database. Curr Protoc Bioinformatics 62, e51.

63. Crooks, G.E., Hon, G., Chandonia, J.-M., and Brenner, S.E. (2004). WebLogo: a sequence logo generator. Genome Res. 14, 1188–1190.

64. Steinegger, M., Meier, M., Mirdita, M., Vöhringer, H., Haunsberger, S.J., and Söding, J. (2019). HH-suite3 for fast remote homology detection and deep protein annotation. BMC Bioinformatics 20, 473.

65. Bailey, T.L. (2021). STREME: accurate and versatile sequence motif discovery. Bioinformatics 37, 2834–2840.

66. Grant, C.E., Bailey, T.L., and Noble, W.S. (2011). FIMO: scanning for occurrences of a given motif. Bioinformatics 27, 1017–1018.

67. Cheng, H., Concepcion, G.T., Feng, X., Zhang, H., and Li, H. (2021). Haplotype-resolved de novo assembly using phased assembly graphs with hifiasm. Nat Methods 18, 170–175.

68. Shumate, A., and Salzberg, S.L. (2021). Liftoff: accurate mapping of gene annotations. Bioinformatics 37, 1639–1643.

69. Benson, G. (1999). Tandem repeats finder: a program to analyze DNA sequences. Nucleic Acids Res 27, 573–580.

70. Bailly-Bechet, M., Haudry, A., and Lerat, E. (2014). “One code to find them all”: a perl tool to conveniently parse RepeatMasker output files. Mob. DNA 5, 13–13.

71. Hwang, H.-J., Kim, Y.T., Kang, N.S., and Han, J.W. (2018). A Simple Method for Removal of the Chlamydomonas reinhardtii Cell Wall Using a Commercially Available Subtilisin (Alcalase). J Mol Microbiol Biotechnol 28, 169–178.

72. Moniruzzaman, M., Erazo-Garcia, M.P., and Aylward, F.O. (2022). Endogenous giant viruses contribute to intraspecies genomic variability in the model green alga. Virus Evol 8, veac102.

73. Chen, S., Zhou, Y., Chen, Y., and Gu, J. (2018). fastp: an ultra-fast all-in-one FASTQ preprocessor. Bioinformatics 34, i884–i890.

74. Bushnell, B., Rood, J., and Singer, E. (2017). BBMerge - Accurate paired shotgun read merging via overlap. PLoS One 12, e0185056.

75. Guo, W., Fiziev, P., Yan, W., Cokus, S., Sun, X., Zhang, M.Q., Chen, P.-Y., and Pellegrini, M. (2013). BS-Seeker2: a versatile aligning pipeline for bisulfite sequencing data. BMC Genomics 14, 774.

76. Jun, G., Wing, M.K., Abecasis, G.R., and Kang, H.M. (2015). An efficient and scalable analysis framework for variant extraction and refinement from population-scale DNA sequence data. Genome Res. 25, 918–925.

77. Tarasov, A., Vilella, A.J., Cuppen, E., Nijman, I.J., and Prins, P. (2015). Sambamba: fast processing of NGS alignment formats. Bioinformatics 31, 2032–2034.

78. Guo, W., Zhu, P., Pellegrini, M., Zhang, M.Q., Wang, X., and Ni, Z. (2018). CGmapTools improves the precision of heterozygous SNV calls and supports allele-specific methylation detection and visualization in bisulfite-sequencing data. Bioinformatics 34, 381–387.

79. Li, H. (2018). Minimap2: pairwise alignment for nucleotide sequences. Bioinformatics 34, 3094–3100.

80. Cheetham, S.W., Kindlova, M., and Ewing, A.D. (2022). Methylartist: tools for visualizing modified bases from nanopore sequence data. Bioinformatics 38, 3109–3112.

81. Ramírez, F., Dündar, F., Diehl, S., Grüning, B.A., and Manke, T. (2014). deepTools: a flexible platform for exploring deep-sequencing data. Nucleic Acids Res. 42, W187–W191.

82. Molnár, A., Schwach, F., Studholme, D.J., Thuenemann, E.C., and Baulcombe, D.C. (2007). miRNAs control gene expression in the single-cell alga *Chlamydomonas reinhardtii*. Nature 447, 1126–1129.

83. Krueger, F. TrimGalore: Consistent adapter and quality trimming for NGS, with extra functionality for RRBS data (Github).

84. Kopylova, E., Noé, L., and Touzet, H. (2012). SortMeRNA: fast and accurate filtering of ribosomal RNAs in metatranscriptomic data. Bioinformatics 28, 3211–3217.

85. Jin, Y., Tam, O.H., Paniagua, E., and Hammell, M. (2015). TEtranscripts: a package for including transposable elements in differential expression analysis of RNA-seq datasets. Bioinformatics 31, 3593–3599.

86. Love, M.I., Huber, W., and Anders, S. (2014). Moderated estimation of fold change and dispersion for RNA-seq data with DESeq2. Genome Biol. 15, 550.

87. Anders, S., and Huber, W. (2010). Differential expression analysis for sequence count data. Derm. Helv., 1–1.

88. Wolf, F.A., Angerer, P., and Theis, F.J. (2018). SCANPY: large-scale single-cell gene expression data analysis. Genome Biol 19, 15.

89. Waskom, M. (2021). seaborn: statistical data visualization. J. Open Source Softw. 6, 3021.

90. Virtanen, P., Gommers, R., Oliphant, T.E., Haberland, M., Reddy, T., Cournapeau, D., Burovski, E., Peterson, P., Weckesser, W., Bright, J., et al. (2020). SciPy 1.0: fundamental algorithms for scientific computing in Python. Nat Methods 17, 261–272.

91. Gel, B., Díez-Villanueva, A., Serra, E., Buschbeck, M., Peinado, M.A., and Malinverni, R. (2016). regioneR: an R/Bioconductor package for the association analysis of genomic regions based on permutation tests. Bioinformatics 32, 289–291.

92. Fu, L., Niu, B., Zhu, Z., Wu, S., and Li, W. (2012). CD-HIT: accelerated for clustering the next-generation sequencing data. Bioinformatics 28, 3150–3152.

93. Katoh, K., and Standley, D.M. (2013). MAFFT multiple sequence alignment software version 7: improvements in performance and usability. Mol Biol Evol 30, 772–780.

94. Capella-Gutiérrez, S., Silla-Martínez, J.M., and Gabaldón, T. (2009). trimAl: a tool for automated alignment trimming in large-scale phylogenetic analyses. Bioinformatics 25, 1972–1973.

95. Wong, T.K.F., Ly-Trong, N., Ren, H., Demotte, P., Baños, H., Roger, A.J., Susko, E., Bielow, C., De Maio, N., Goldman, N., et al. (2026). IQ-TREE 3: phylogenomic inference software using complex evolutionary models. Mol Biol Evol 43. 10.1093/molbev/msag117.

96. Kalyaanamoorthy, S., Minh, B.Q., Wong, T.K.F., von Haeseler, A., and Jermiin, L.S. (2017). ModelFinder: fast model selection for accurate phylogenetic estimates. Nat Methods 14, 587–589.

