## Supplementary appendix for "A chimeric infection program and epigenetic conflict underpin endogenous giant virus latency"

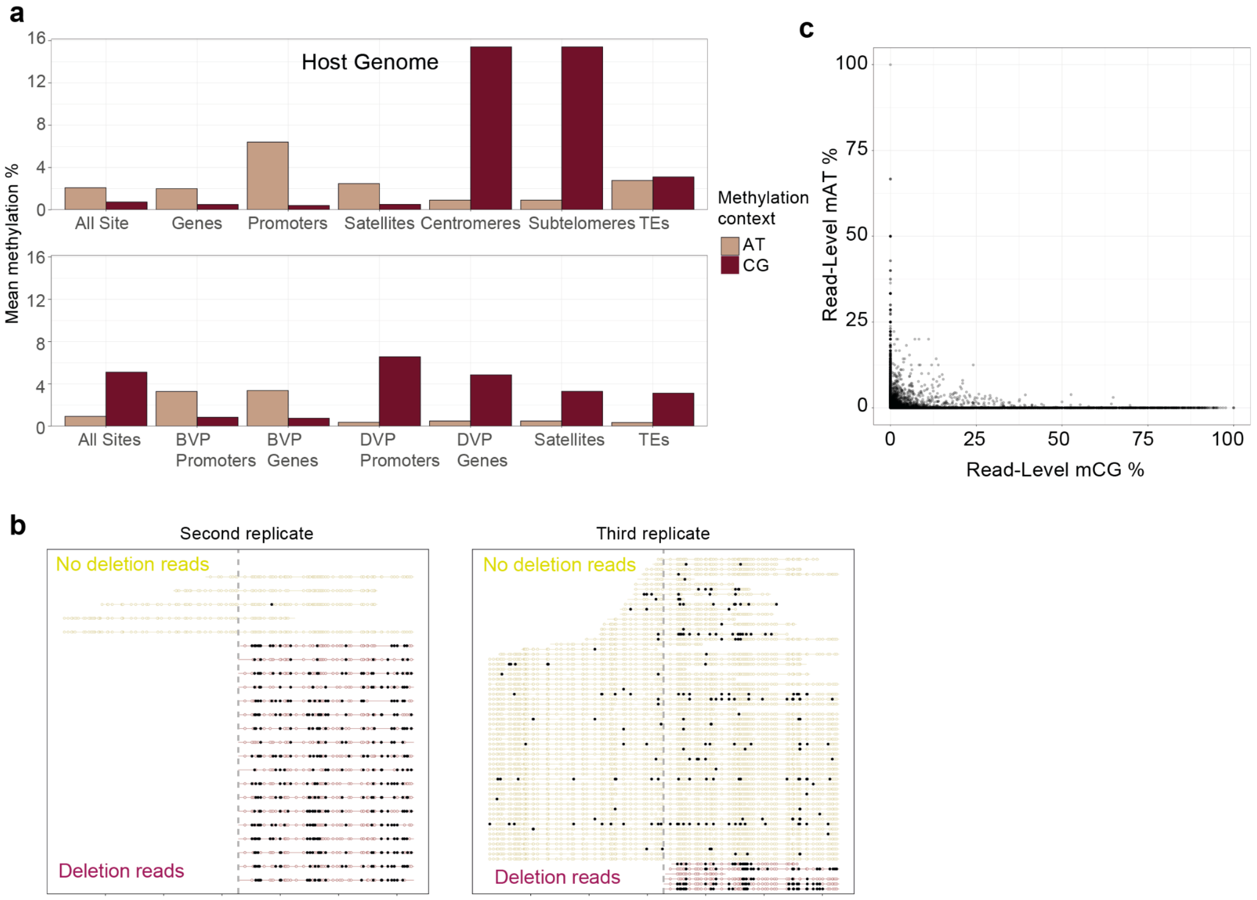


**Appendix Figure 1. Distribution of DNA base modifications across the *C. reinhardtii* and GEVE genomes.** (A) Mean levels of 5mC (mCG) and 6mA (mAT) across major genomic features in GEVE+ strains, including host genes, transposable elements, and the GEVE. The two DNA modifications show largely complementary genomic distributions. (B) Representative per-read 5mC profiles across the GEVE from one mixed culture, stratified by reads spanning the internal deletion (full-length GEVE) or terminating at the deletion breakpoint (deletion GEVE). The dashed vertical line marks the 3′ deletion breakpoint; filled circles indicate methylated CpGs and open circles unmethylated CpGs. Deletion-containing molecules show substantially higher 5mC across the retained GEVE sequence. (C) Distribution of per-read methylation levels overlapping the GEVE at 5mC and 6mA sites in GEVE+ strains. Only reads containing at least 10 CG sites and 10 AT sites were included. The two modifications occur largely in distinct subsets of molecules, consistent with the mutually exclusive molecular distribution of 5mC and 6mA.

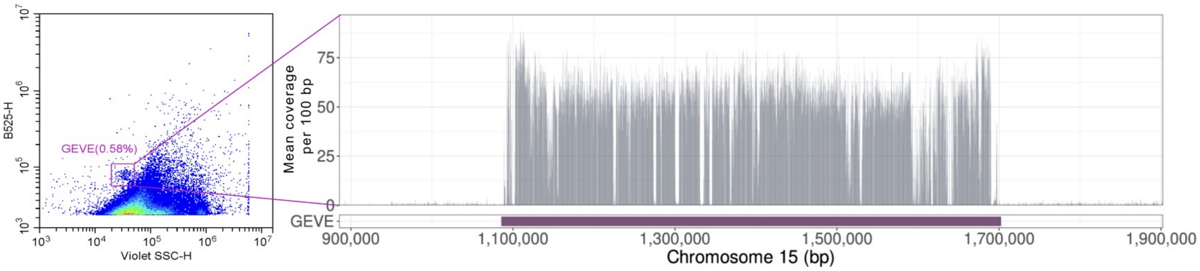


**Appendix Figure 2.** Punuivirus virion isolation and Enzymatic Methyl-seq (EM-seq) coverage of the viral genome. The left panel shows stained punuivirus particles isolated from GEVE+ cultures detected by flow cytometry. Virion samples were DNase treated prior to viral DNA extraction. The right panel displays EM-seq coverage across chromosome 15, with viral reads strongly enriched over the GEVE region relative to residual host DNA.

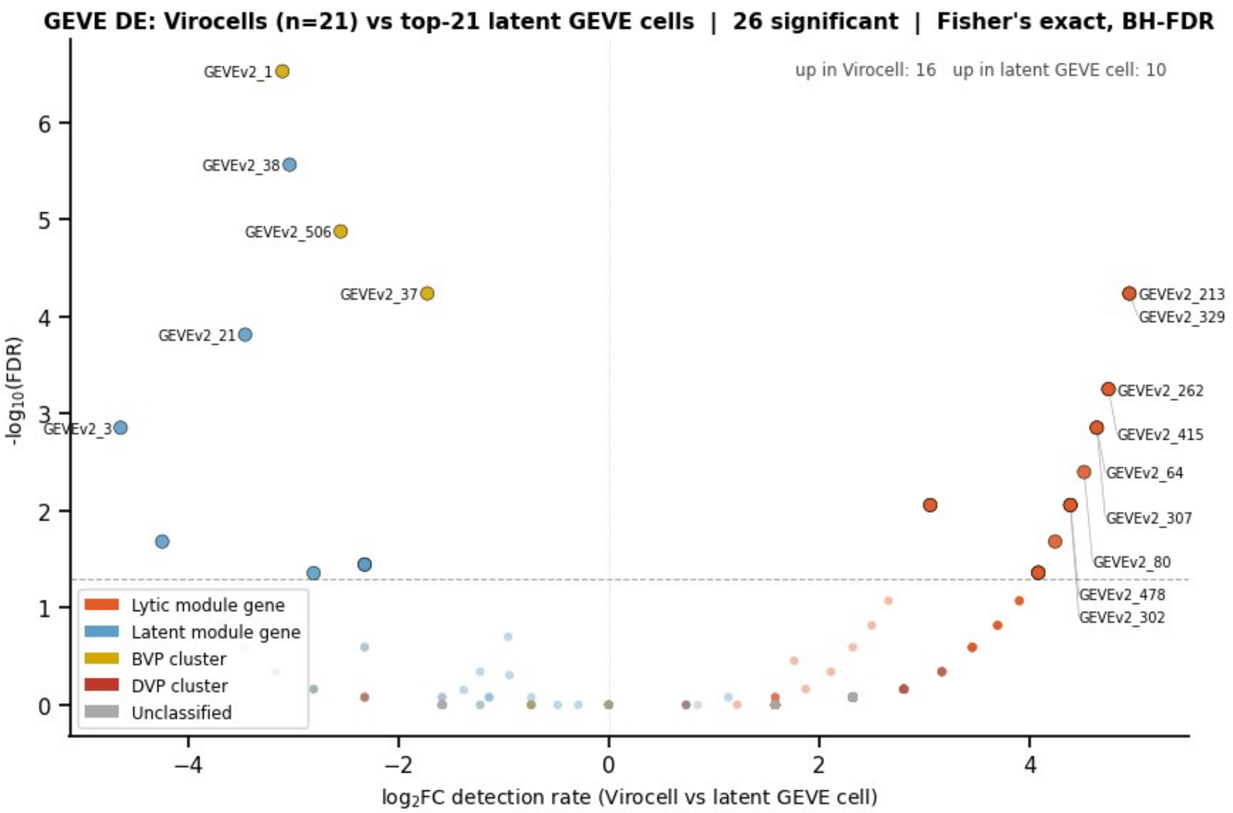


**Appendix Figure 3. Volcano plot of GEVE gene differential expression: Virocells versus top-21 High-GEVE latent cells.** Fisher's exact test on per-gene detection rates for 315 GEVE genes in 21 virocells versus the 21 High-GEVE latent cells with the highest total GEVE UMI counts (top-21 High-GEVE latent cells; all from DPI 4; not matched to virocells). x-axis: $\log_2$ fold-change in detection rate (positive = higher in Virocells). y-axis: $-\log_{10}$(BH-FDR). Dashed line marks FDR = 0.05. Genes colored by lytic-module (orange), latent-module (blue), BVP (gold), DVP (red), or unclassified (gray) membership. Of 315 GEVE genes tested, 26 reached FDR < 0.05: 16 more detected in virocells and 10 in the comparison group.

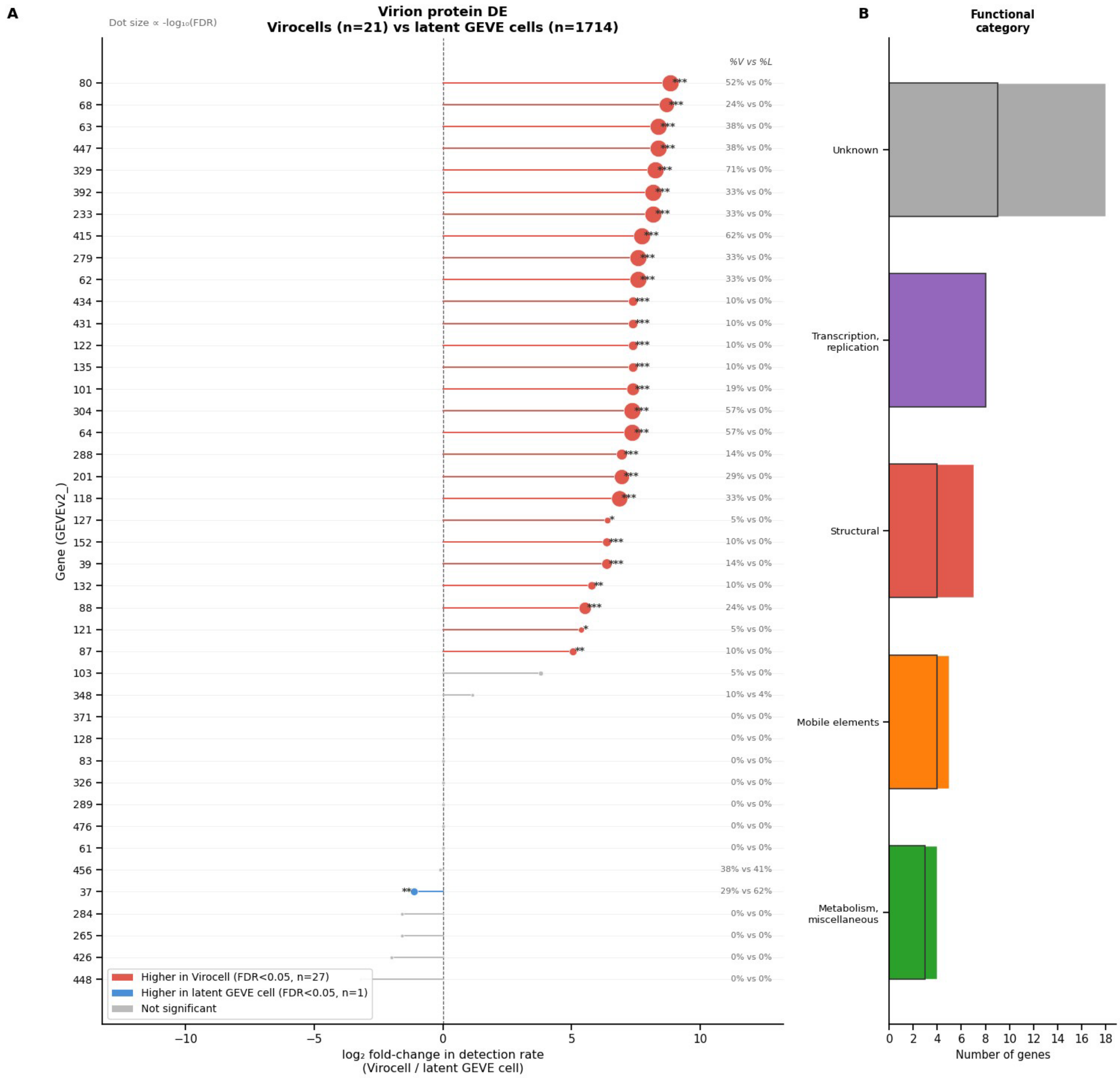


**Appendix Figure 4. Differential detection of GEVE virion protein genes in virocells versus all latent GEVE cells.** (A) Lollipop chart of $\log_2$ fold-change in per-gene detection rates (virocells, n = 21 / all latent GEVE cells, GEVE ≥ 10 UMIs and not virocell, n = 1,714) for all 42 virion protein genes (LC-MS/MS from Erazo-Garcia et al.[1]). Red = significantly more detected in virocells (FDR < 0.05); blue = higher detection in all latent GEVE cells; gray = not significant. Dot size proportional to $-\log_{10}$(FDR). Of 42 genes tested, 27 were significantly more detected in virocells and one (GEVEv2_37) in all latent-GEVE cells. (B) Distribution of virion protein genes across functional categories.

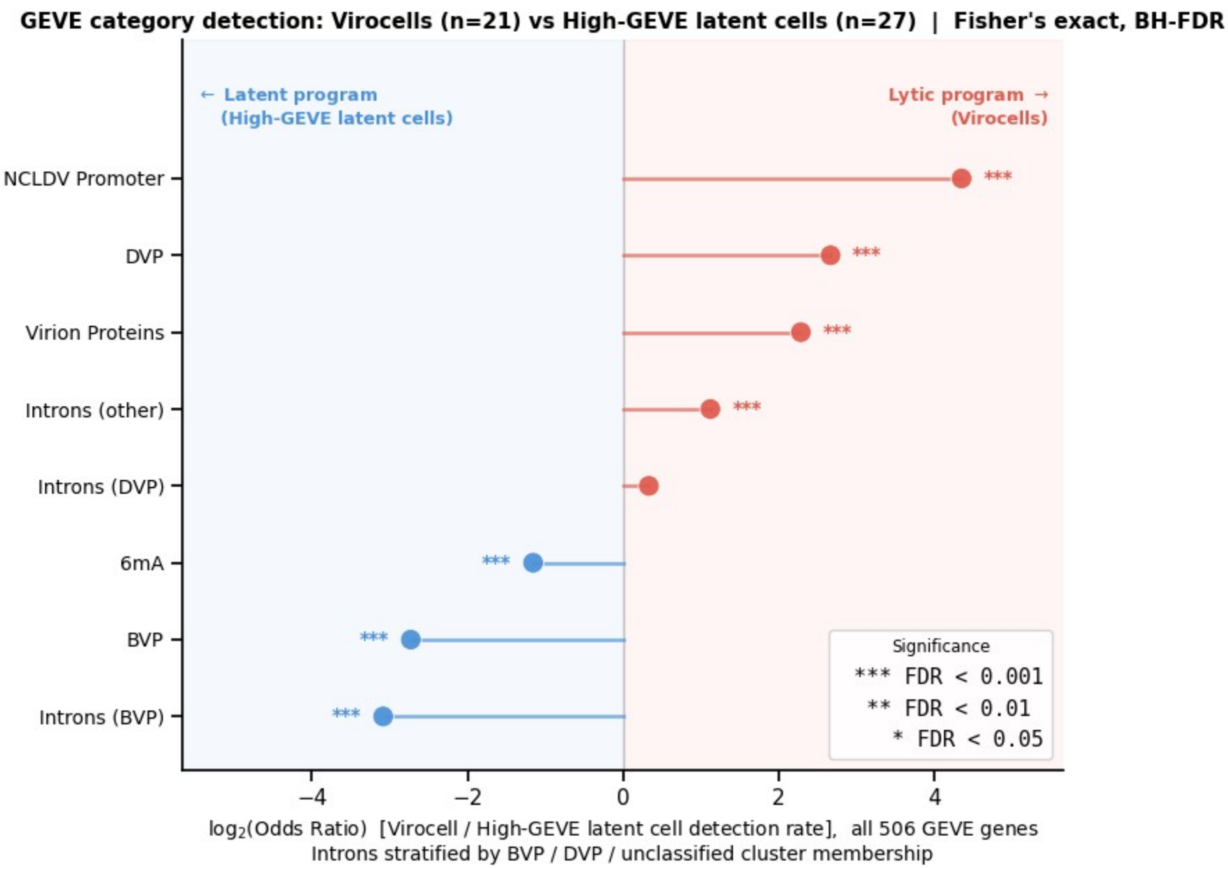


**Appendix Figure 5. GEVE genomic feature categories distinguish the transcriptional states of virocells and High-GEVE latent cells.** Diverging lollipop plot showing $\log_2$(OR) for detection-rate enrichment of eight GEVE genomic feature categories in virocells versus High-GEVE latent cells, tested across all 506 canonical GEVE genes (21 virocells; 27 High-GEVE latent cells, GEVE ≥ 43 UMIs, all from DPI 4; Fisher's exact test, BH-FDR). Intron-bearing genes are stratified by BVP, DVP, or unclassified cluster. $\log_2$(OR) > 0 = enriched in virocells (lytic); $\log_2$(OR) < 0 = enriched in High-GEVE latent cells (latent). DVP-cluster (OR = 6.36), NCLDV-type promoter (OR = 20.4), and virion structural protein (OR = 4.88) genes, as well as unclassified intron-bearing genes (OR = 2.18), were enriched in virocells; DVP-cluster intron-bearing genes did not differ (OR = 1.26, FDR = 0.36). BVP-cluster genes (OR = 0.151), BVP-cluster intron-bearing genes (OR = 0.118), and 6mA-modified genes (OR = 0.449, FDR = $1.2 \times 10^{-8}$) were enriched in High-GEVE latent cells. Full statistics in **Supplementary Table 8**.

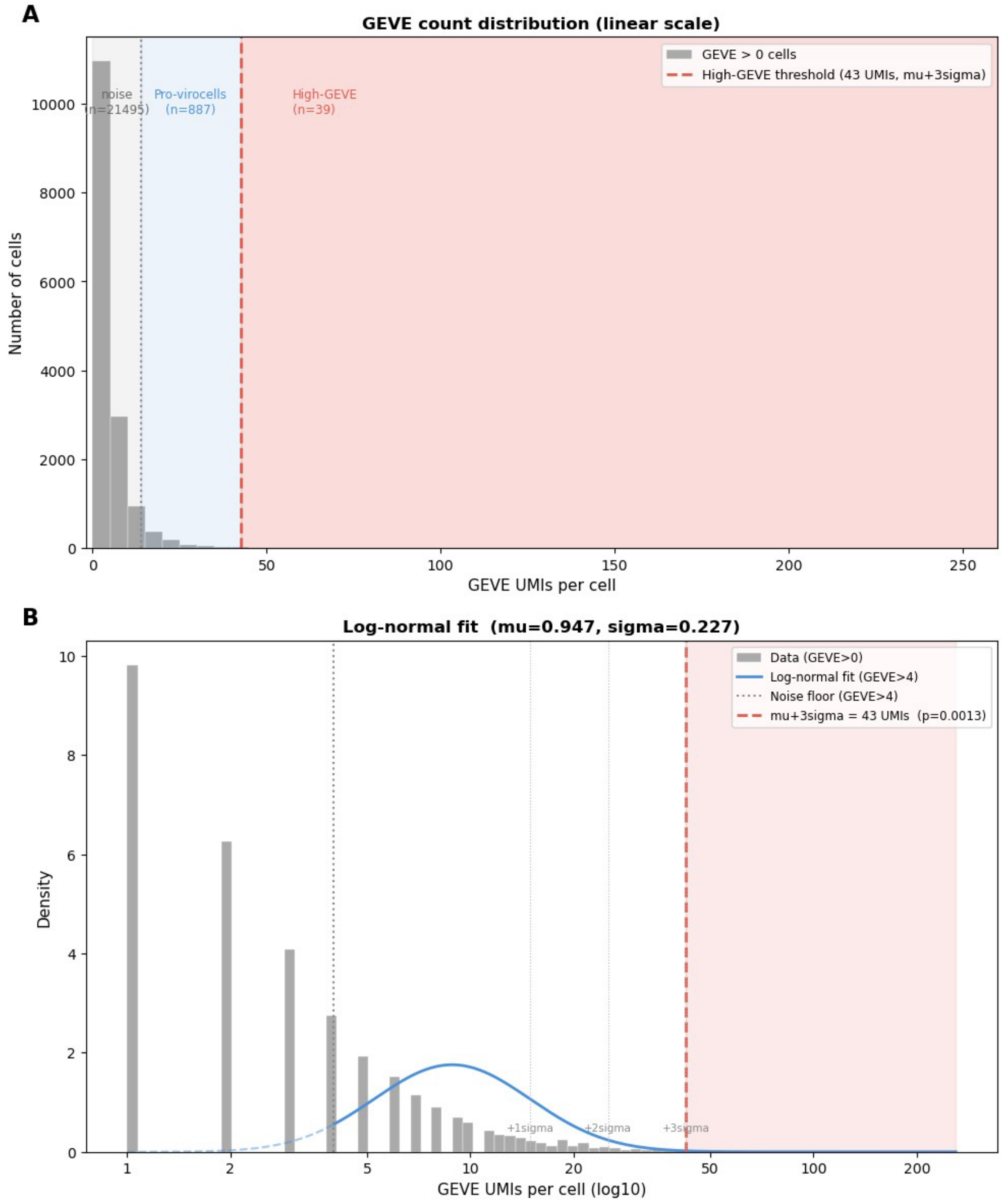


**Appendix Figure 6. Data-driven definition of cell-state boundaries from the GEVE UMI distribution.** (A) Histogram of GEVE UMIs per cell (linear scale; 5-UMI bins) for all cells with at least one GEVE UMI. Shaded regions mark the noise floor (gray; GEVE $\leq 4$), the Active GEVE Expressing Cells range (blue; 10–42 UMIs), and High-GEVE range (red). Dashed red line = High-GEVE threshold ($\mu + 3\sigma$). (B) Same distribution on $\log_{10}$ axis (density-normalized), overlaid with fitted log-normal (blue solid; fit to GEVE > 4; dashed extension below noise floor). Dotted vertical lines mark $+1\sigma$, $+2\sigma$, $+3\sigma$ from fitted mean. $\mu + 3\sigma = 43$ UMIs (one-tailed $p = 0.0013$). An earlier manual threshold of 50 UMIs ($\approx \mu + 3.3\sigma$, $p = 0.0005$) selected a subset (24 cells) of the same outlier population.

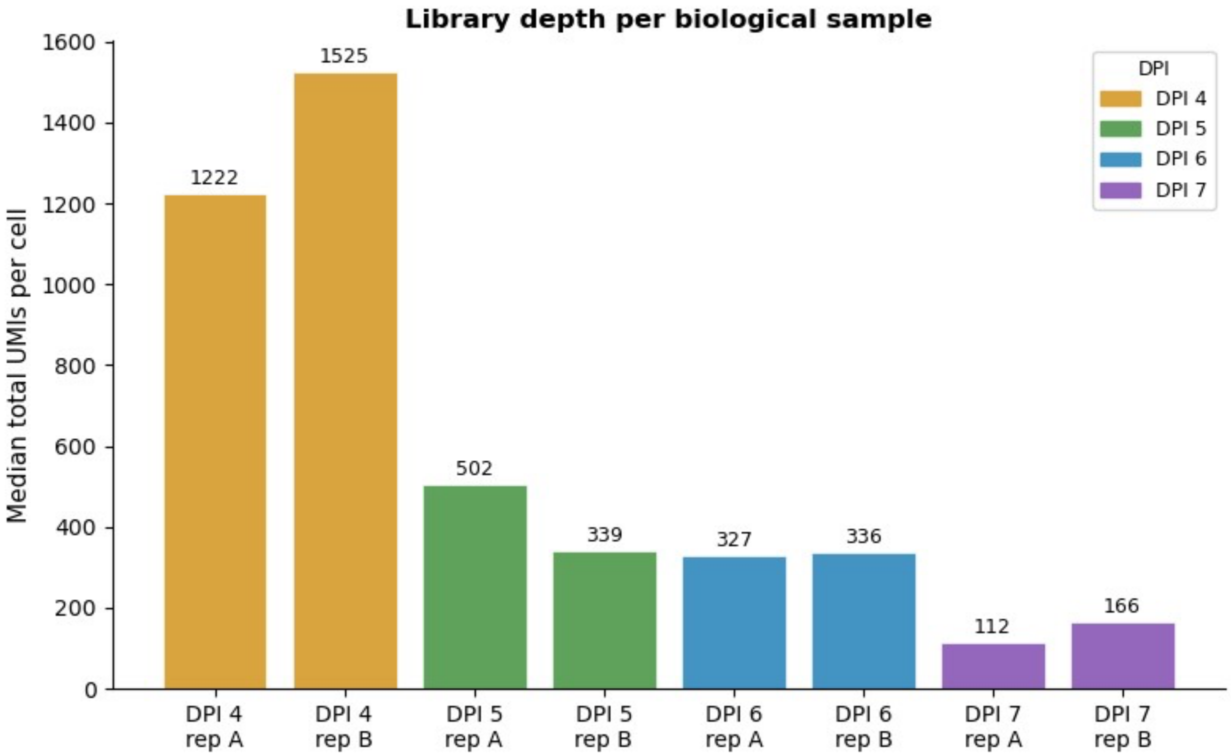


**Appendix Figure 7. Median library depth per biological sample across DPI 4–7.** Each bar shows the median total UMI count per QC-passing cell for one biological replicate (A or B). Colors indicate DPI. Median library depth declines from DPI 4 (> 1,000 UMIs/cell) to DPI 7 (< 500 UMIs/cell), consistent with the transition from exponential growth to stationary phase.

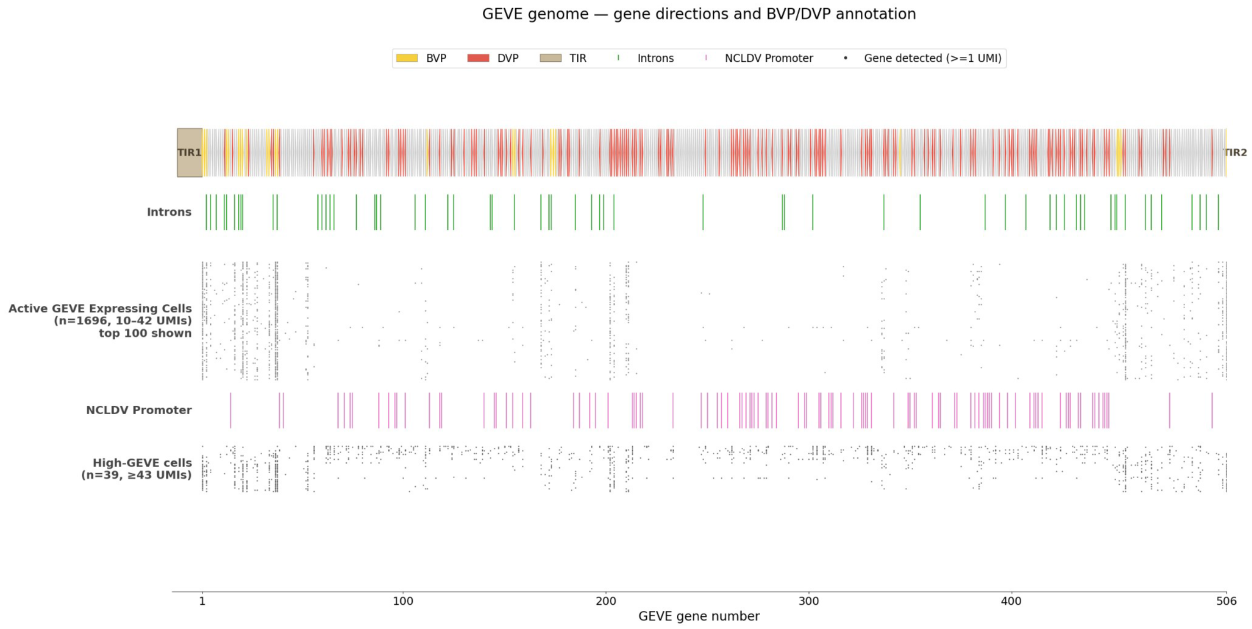


**Appendix Figure 8. Single-cell GEVE transcriptomes mapped onto the viral genome for Active GEVE Expressing Cells and High-GEVE cells.** Top panel: 506 annotated GEVE genes shown as directional arrows colored by cluster (BVP yellow, DVP red, Unclassified gray). TIR1 and TIR2 flanking rectangles. Introns track (green ticks; spliceosomal introns only); NCLDV Promoter track (pink ticks). Lower panels: each horizontal row is one cell; a dot indicates detection of ≥ 1 UMI for that gene. Active GEVE Expressing Cells (upper band; 100 highest-load cells shown out of n = 1,696) have fewer dots concentrated in the BVP region. High-GEVE cells (lower band) show broader detection across the DVP region.

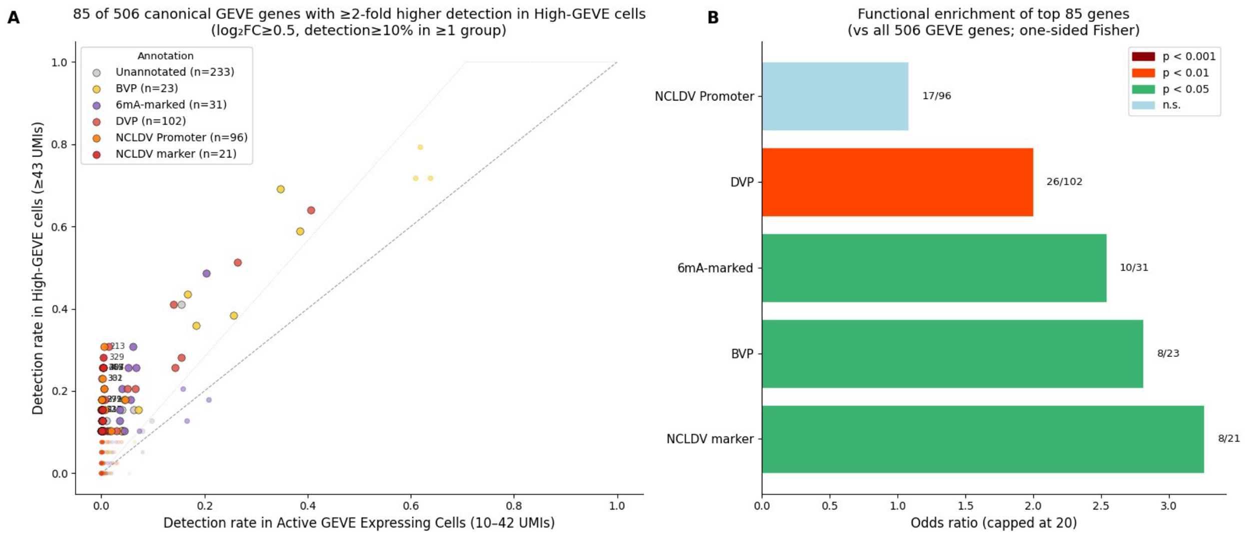


**Appendix Figure 9. Detection-rate comparison of GEVE genes between Active GEVE Expressing Cells and High-GEVE cells.** All 506 canonical GEVE genes ranked by $\log_2$ fold-change in detection rate (Active GEVE Expressing Cells: GEVE 10–42 UMIs, including 9 virocells, n = 1,696; High-GEVE: GEVE ≥ 43 UMIs, n = 39). (A) Scatter plot of detection rate in Active GEVE Expressing Cells (x-axis) versus High-GEVE cells (y-axis). Points colored by functional annotation. Filled circles with black outline = genes with substantially higher detection in High-GEVE ($\log_2FC \geq 0.5$, detection rate ≥ 10% in at least one group). (B) Functional enrichment of genes with substantially higher detection in High-GEVE cells (one-sided Fisher's exact test). Bars show OR for each functional category.

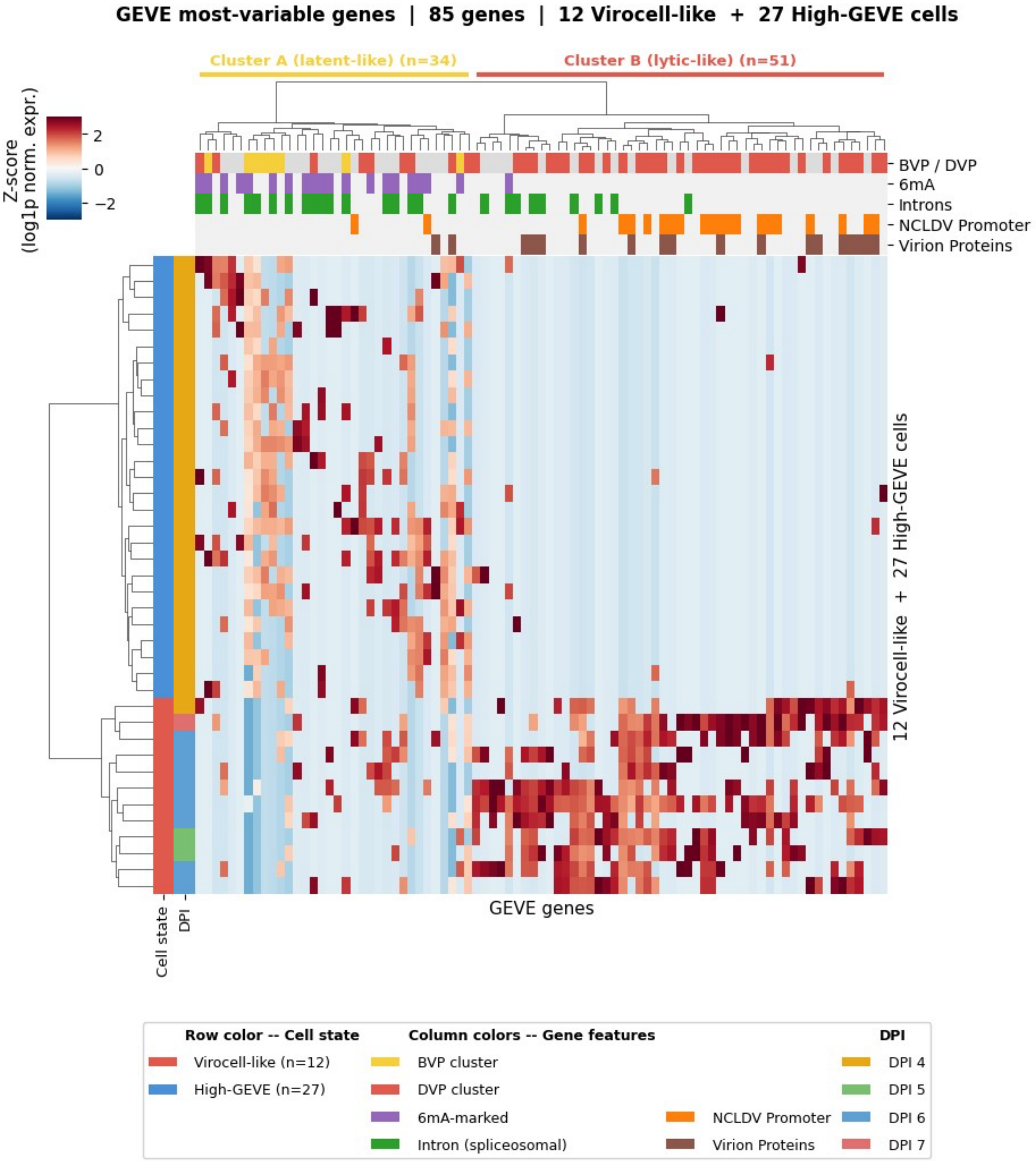


**Appendix Figure 10. Heatmap of most-variable GEVE genes across the 39 High-GEVE cells.** Rows: 39 High-GEVE cells (GEVE ≥ 43 UMIs). Columns: 85 GEVE genes with substantially higher detection in High-GEVE vs. Active GEVE Expressing Cells ($\log_2$FC ≥ 0.5, detection ≥ 10%). Values: $\log_2$(1 + 10k-normalized counts), Z-scored per gene, clipped at ±3. Both axes hierarchically clustered (Ward / Euclidean). Row color bar: red = 12 virocell cells; blue = remaining 27 High-GEVE cells. Column color bars: BVP/DVP cluster, 6mA, NCLDV promoter, spliceosomal intron. Gene labels include GEVE gene number plus NCLDV marker / Pfam / GVOG family where available.

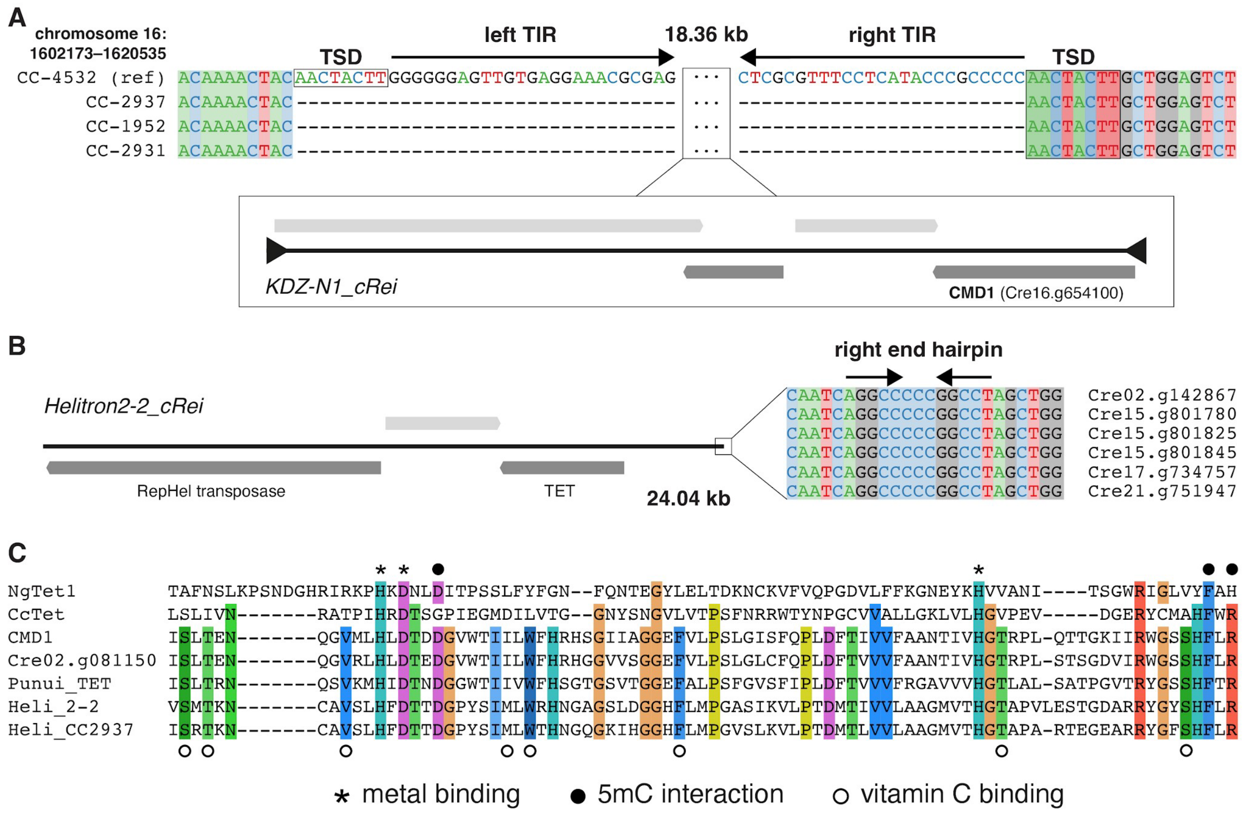


**Appendix Figure 11.** Transposon and Punuivirus TET genes of *Chlamydomonas reinhardtii*. (A) Example of polymorphic insertion of the *KDZ-N1_cRei* transposon carrying *CMD1* TET gene (Cre16.g654100) on chromosome 16 of the reference genome (strain CC-4532) relative to the genomes of the wild strains CC-2937, CC-1952 and CC-2931[2]. The 18.36 kb DNA transposon produces 8-bp target site duplications (TSD) upon insertion and carries four genes, although none are recognised as a transposase. It is tentatively classified as a nonautonomous transposon from the *KDZ* (*Kyakuja-Dileera-Zisupton*) superfamily based on the TSD length, terminal inverted repeat sequence and overall length of the element. Note that *CMD1* was originally described as the gene Cre12.g553400[3], although both genes are carried by copies of the same transposon and the encoded proteins are 100% identical. Also note that all of the genes contain introns, which are not shown. (B) Alignment of the right end region of the Helitron transposon *Helitron2-2_cRei* and schematic of three genes (introns not shown) carried by these elements. *Helitron2-2_cRei* is present in at least 8 copies in the reference strain and in two distinct copies in the CC-2937 genome. (C) Alignment of catalytic double-stranded β-helix (DSBH) domain of TET proteins from *C. reinhardtii* and punuivirus to characterised TET enzymes from *Naegleria fowleri* (NgTet1) and *Coprinopsis cinerea* (CcTet)[4,5]. Conserved amino acids involved in metal ion binding and 5mC interaction are highlighted above the alignment, and amino acids in CMD1 that are involved in binding the vitamin C co-substrate (Li, et al. 2021) are highlighted below (note that these are only conserved among *C. reinhardtii* and Punuivirus TETs, since NgTet1 and CcTet bind 2-oxoglutarate instead).

### Appendix Methods

*Single-cell reference genome and read processing*

A custom reference genome and annotation were built for single-cell processing from the *Chlamydomonas reinhardtii* CC-2937 genome and the integrated GEVE, using a GTF file that defined gene, exon, and intron coordinates for each gene. The reference was indexed using split-pipe (Parse Biosciences, version 1.1.2) in mkref mode. Raw reads were processed in all mode with chemistry v2 to align reads and assign transcripts to their sample of origin by combinatorial barcode, and the two sequenced sublibraries were merged using split-pipe in comb mode to generate a unified digital gene expression matrix.

*Expression normalization*

For analyses requiring normalized expression values, raw UMI counts were library-size normalized to a fixed total of 10,000 UMIs per cell (CP10K) and natural-log-transformed (log1p) [Scanpy 1.10.1][6,7], yielding log1p-normalized expression values used throughout the analyses below unless otherwise noted.

*Visualization of GEVE gene detection along the linear genome*

To visualize the distribution of GEVE gene detection along the linear genome, each GEVE-expressing cell was represented as a horizontal row in which a dot at a given genomic position indicates detection of at least one UMI for the corresponding GEVE gene. Cells were stratified into Active GEVE Expressing Cells (10–42 GEVE UMIs) and High-GEVE ($\geq 43$ UMIs) groups and plotted in register with the annotated GEVE genome track, showing gene arrows, BVP/DVP cluster assignment, spliceosomal intron positions, and NCLDV-type promoter elements (Appendix Figure 8). This visualization revealed that Active GEVE Expressing Cells predominantly detect BVP-cluster genes near the center of the genome, whereas High-GEVE cells detect a broader set of genes spanning the full DVP-enriched regions [Scanpy 1.10.1, AnnData 0.10.8, Matplotlib 3.8.4][6,7].

*GEVE expression threshold identification*

To define an empirical upper threshold separating High-GEVE cells from the bulk GEVE-expressing population, the per-cell GEVE UMI count distribution was modeled by fitting a log-normal distribution [SciPy 1.13.0][8,9] to all cells with GEVE > 4 UMIs (noise floor, below which per-cell GEVE counts are consistent with ambient RNA; $n = 4{,}688$; Appendix Figure 6). The fitted log-normal yielded a geometric mean of $\mu = 8.8$ UMIs; the upper tail defined a statistically motivated High-GEVE threshold at $\mu + 3\sigma = 43$ UMIs (one-tailed $p = 0.0013$ under the fitted distribution; 39 cells). Because no bimodal gap is visible in the distribution, this threshold does not reflect a natural break in expression but rather captures a statistically extreme subpopulation whose GEVE expression lies more than three standard deviations above the log-normal mean. An earlier manual threshold of 50 UMIs (corresponding to $\mu + 3.3\sigma$, $p = 0.0005$) selected a subset (24 cells) of the same outlier population. Median library depth (total UMIs per cell) was compared across samples and DPI (Appendix Figure 7). Depth was highest at DPI 4 (median 1,222–1,525 UMIs per cell), lower at DPI 5 (339–503) and several-fold lower at DPI 6–7 (112–336). Because 20 of 21 virocells come from the more shallowly sequenced DPI 5–7 samples, their high GEVE counts are not explained by sequencing depth. In contrast, 28 of the 39 High-GEVE cells, and all High-GEVE latent comparison cells, come from DPI 4, so comparisons between virocells and High-GEVE latent cells are confounded with depth and culture stage.

*Differential detection and lytic/latent module identification*

To identify GEVE genes with preferentially elevated expression in High-GEVE cells, detection rate (fraction of cells with at least one UMI) was computed for all 506 canonical GEVE genes separately in Active GEVE Expressing Cells (GEVE 10–42 UMIs; n = 1,696, including 9 virocells) and High-GEVE cells (GEVE ≥ 43 UMIs; n = 39) (Appendix Figure 9). Genes were ranked by $\log_2$ fold-change in detection rate between the two groups; 85 genes with $\log_2 FC \geq 0.5$ and detection rate ≥ 10% in at least one group were selected for hierarchical clustering. Z-scored log1p-normalized expression values for these 85 genes across the 39 High-GEVE cells were clustered using Ward linkage and Euclidean distance [SciPy 1.13.0, Seaborn 0.13.2][8,9] (Appendix Figure 10). Two stable gene clusters emerged: a lytic module of 51 genes (34 DVP-cluster, 17 unclassified) showing coordinated high expression in the 12 cells with the broadest GEVE expression profiles, and a latent module of 34 genes (8 BVP-cluster, 9 DVP-cluster, 17 unclassified) expressed at lower but more uniform levels across all 39 High-GEVE cells. The gene cluster with the higher mean Z-score in these 12 cells relative to the other 27 High-GEVE cells was designated the lytic module; the other cluster was designated the latent module. For each GEVE-expressing cell, the lytic/latent module ratio was then defined as the $\log_2$ ratio of mean log1p-normalized expression across the 51 lytic-module genes versus the 34 latent-module genes; this scalar quantity is used throughout to color-code cells by transcriptional program (Figure 2A and B). For cell-state classification (STAR Methods, Cell-state classification), the same ratio was computed on mean raw UMI counts per module gene, $\log_2$[(mean lytic UMIs + 1) / (mean latent UMIs + 1)].

### References


1. Erazo-Garcia, M.P., Sheyn, U., Barth, Z.K., Craig, R.J., Wessman, P., Jivaji, A.M., Ray, W.K., Svensson-Coelho, M., Cornwallis, C.K., Rengefors, K., et al. (2025). Cryptic infection of a giant virus in a unicellular green alga. Science *388*, eads6303.
2. López-Cortegano, E., Craig, R.J., Chebib, J., Balogun, E.J., and Keightley, P.D. (2023). Rates and spectra of *de novo* structural mutations in *Chlamydomonas reinhhardtii*. Genome Res *33*, 45–60.
3. Xue, J.-H., Chen, G.-D., Hao, F., Chen, H., Fang, Z., Chen, F.-F., Pang, B., Yang, Q.-L., Wei, X., Fan, Q.-Q., et al. (2019). A vitamin-C-derived DNA modification catalysed by an algal TET homologue. Nature *569*, 581–585.
4. Hashimoto, H., Pais, J.E., Dai, N., Corrêa, I.R., Jr, Zhang, X., Zheng, Y., and Cheng, X. (2015). Structure of Naegleria Tet-like dioxygenase (NgTet1) in complexes with a reaction intermediate 5-hydroxymethylcytosine DNA. Nucleic Acids Res *43*, 10713–10721.
5. Zhang, L., Mu, Y., Li, T., Hu, J., Lin, H., and Zhang, L. (2024). Molecular basis of an atypical dsDNA 5mC/6mA bifunctional dioxygenase CcTet from *Coprinopsis cinerea* in catalyzing dsDNA 5mC demethylation. Nucleic Acids Res *52*, 3886–3895.
6. Wolf, F.A., Angerer, P., and Theis, F.J. (2018). SCANPY: large-scale single-cell gene expression data analysis. Genome Biol *19*, 15.
7. Virshup, I., Rybakov, S., Theis, F.J., Angerer, P., and Alexander Wolf, F. (2024). anndata: Access and store annotated data matrices. Journal of Open Source Software *9*, 4371.

8. Waskom, M. (2021). seaborn: statistical data visualization. J. Open Source Softw. *6*, 3021.

9. Virtanen, P., Gommers, R., Oliphant, T.E., Haberland, M., Reddy, T., Cournapeau, D., Burovski, E., Peterson, P., Weckesser, W., Bright, J., et al. (2020). SciPy 1.0: fundamental algorithms for scientific computing in Python. Nat Methods *17*, 261–272.
